# Underwater photo-identification of marine megafauna: an identity card catalogue of sperm whales (*Physeter macrocephalus*) off Mauritius Island

**DOI:** 10.1101/2021.03.08.433909

**Authors:** Sarano Véronique, Sarano François, Girardet Justine, Preud’homme Axel, Vitry Hugues, Heuzey René, Sarano Marion, Delfour Fabienne, Glotin Hervé, Adam Olivier, Madon Bénédicte, Jung Jean-Luc

**Author notes:** Corresponding author. Mail.

## Abstract

The long-term monitoring of long-lived animal populations often requires individual identification. For cetacean populations, this identification is mostly based on morphological characters observable from a boat such as shape, spots and cuts of the back, caudal and dorsal fins. This is well suited for species easily displaying their caudal fins, such as the humpback whales *Megaptera novaeangliae*, or those whose skin pigmentation patterns enable individual identification.

However, for elusive or shier species such as the sperm whales *Physeter macrocephalus*, this approach may be more challenging as individuals display a rather uniform skin pigmentation. They also do not show very often their caudal fin that must be photographed perpendicularly to the water surface, vertically and fully emerged, uneasing the individual identification from a boat. Immature sperm whales that usually have a caudal fin without any distinctive marks may sometimes be excluded from photo-identification catalogues.

Within the framework of the Maubydick project, focusing on the long-term monitoring of sperm whales in Mauritius, passive underwater observation and video recording were used to identify long-lasting body markers (e.g., sex, ventral white markings, cut outs of fins) to improve individual identification. A catalogue of individual identity cards was developed and 38 individuals were recorded (six adult males, 18 adult females and 14 immatures). This catalogue was used in the field and enabled observers to record some nearly-daily and yearly recaptures. Advantages and disadvantages of this method are presented here.

Such catalogues represent a robust baseline for conducting behavioural, genetic and acoustic studies in marine megafauna social species. Benefits of such newly acquired knowledge are of first importance to implement relevant conservation plans in the marine realm.

## INTRODUCTION

The long-term study of long-lived animal population often requires individual identification, e.g. for abundance estimation in mark-recapture surveys, social behaviour understanding and for conservation purposes (Hammond et al. 1990, Würsig & Jefferson 1990, Gowans & Whitehead 2001, Möller et al. 2006, Calambokidis et al. 2008, Gero et al. 2014, Cantor & Whitehead 2016, Gero & Whitehead 2016, Augusto et al. 2017, Louis et al. 2017, Huisjer et al. 2020, Sarano et al. 2021).

This identification may be challenging in the marine environment and cetaceans are no exception, spending only a limited amount of time at the sea surface. The individual identification is then based on a reduced number of morphological characteristics captured on photographs taken from a boat or an unmanned aerial vehicle (Verfuss et al. 2019). The main morphological characteristics that can be observed are the coloring of the back, the shape of the dorsal fin and/or the distinct markings on the trailing edge of the caudal, the latter being only visible when the animal flukes (Arnbom 1987, Sears et al. 1990, Whitehead 1990, Dufault & Whitehead 1995, Gomez-Salazar et al. 2011). Algorithms have been developed to automate the fastidious task of visual inspection of photographs in the search for potential recaptures (*e*.*g*. Whitehead 1990, Huele & Udo de Haes 1998, White et al. 1998, Huele & Ciano 1999, Beekmans et al. 2005, Hillman et al. 2010, Levenson et al. 2015).

Underwater observation may be used to gather additional information that cannot be collected from a boat alone. Underwater devices such as unmanned underwater drones, for example, may be used. When the environment permits it, human observers can also film and perform accurate underwater individual identification. Such approaches have been used successfully in humpback whales *Megaptera novaeangliae* (Glockner-Ferrari & Ferrari 1990), dolphins *Tursiops truncatus* (Herzing 1997), manta rays *Mobula* spp (Town et al. 2013, Marshall & Holmberg 2018) or whale sharks *Rhincodon typus* (Pierce et al. 2018) to develop catalogues of individuals. To complete underwater video and photographic data, the observers can also collect samples and outcompete methods traditionally used for studies focused on genetics (Sarano et al. 2021).

For sperm whales (*Physeter macrocephalus*), a lot has been learnt through boat-based observations (see for instance Alessi et al. 2014, Carpinelli et al. 2014, Gero et al. 2014, Cantor & Whitehead 2016, Gero & Whitehead 2016, Cantor et al. 2019, Van der Linde & Eriksson 2020). Individual identification of sperm whales is in general based on mark patterns of the fluke (Arnborn 1987). Body marks can also be used (eg Alessi et al. 2014, Van der Linde & Eriksson 2020). But individual identification can sometime be difficult as some individuals have a dorsal fin barely distinctive (Van der Linde & Eriksson 2020) and as sperm whales have a caudal fin of uniform color unlike humpback whales for example (Mizroch et al. 1990). Young immature individuals rarely fluke, making their identification particularly difficult (Whitehead 2006, Gero et al. 2009). Their sexing is impossible from the surface as they do not show any apparent sexual dimorphism. Adult females and large immatures have similar sizes, and may therefore be difficult to distinguish (Gero et al. 2014). The capture probability may also differ between individuals, some spending less time at the surface or having no visible distinctive signs may escape identifications (Whitehead 2006). As a result, in photo-identification (photo-ID) monitoring of sperm whale populations, some individuals may remain unidentifiable from the sea surface (Whitehead 2006, Boys et al. 2019, Van der Linde and Eriksson 2020, Kobayashi et al. 2020). Underwater observation may therefore in some cases help to identify individual sperm whales, as more discriminating markers, e.g., located on the ventral part of the animals, could be observed. It would also help to infer gender with certainty.

This paper presents the results of a study based on an underwater photo-ID and video-identification (video-ID) protocol used to monitor sperm whales in Mauritius since 2015 by the French association Longitude 181, in the framework of the Maubydick program run by the Mauritian NGO Marine Megafauna Conservation Organization. In this long-term conservation program, the social organization and the dynamics of groups of sperm whales off west Mauritius are studied using video-recorded underwater observations, genetics analysis (Sarano et al. 2021) and acoustics (Ferrari et al. 2019, 2020). A unique catalogue of 38 sperm whales based on identity cards (ID-cards) displaying long-lasting and reliable morphological markers for each individual was created. These results should be of first interest in terms of conservation of the species in the Indian Ocean.

## MATERIAL AND METHODS

### Field observations

Sperm whales are common off the coast of the Mauritius Island (Mascarenes Islands, Indian Ocean). A protocol based on underwater observations through photography and video recording was implemented in 2011 for the Maubydick project led by the MMCO (Marine Megafauna Conservation Organization, Mauritius Island). In 2015, the protocol was standardized under the scientific lead of Longitude 181 association (France), and the sampling effort increased over the years since then (Table 1).

**Table 1:** Number of days of field work. The number of days of field work per year is indicated, as well as days with and without observation of sperm whales (in number of days and in percentage of days of field work). The protocol based on underwater observations was implemented in 2011 and standardized in 2015.

|  |  |  |  |  |  |  |  |  |  |  | Total numbers of field work days and of observations |  |
| --- | --- | --- | --- | --- | --- | --- | --- | --- | --- | --- | --- | --- |
| 2011-2014 field work |  |  |  |  | 2015-2020 Field work |  |  |  |  |  |  |  |
| Years | 2011 | 2012 | 2013 | 2014 | 2015 | 2016 | 2017 | 2018 | 2019 | 2020 | 2011-2020 | 2015-2020 |
| Numbers of days of field work | 13d | 5d | 10d | 12d | 36d | 40d | 54d | 70d | 81d | 36d | 357d | 317d |
| Days with no observations (numbers, percentage) | 2 | 1 | 0 | 0 | 7d, 19.4% | 6d, 15.0% | 4d, 7.4% | 6d, 8.6% | 13d, 16.1% | 15d, 41.7% | 54d, 15.1% | 51d, 16.1% |
| Days with observation of sperm whales (numbers, percentage) | 11 | 4 | 10 | 12 | 29d, 80.6% | 34d, 85% | 50d, 92.6% | 64d, 91.4% | 68d, 83.9% | 21d, 58.3% | 303d, 84.9% | 266d, 83.9% |

The study area is located on the west coast of the Mauritius Island, up to 15 km off the coast, between 20.465S 57.334E and 19.986S 57.605E (Sarano et al. 2021). The boat used for this survey is a 15-meter Mauritian motor vessel, chartered by MMCO and equipped for diving with a low rear platform, from which observers can immerse themselves by gently sliding into the water. All underwater observations were video-recorded, either with a Sony F55 4K, a Sony EXIR HD, a Nikon D800 Camera in Hugyfot housing or a GoPro camera Hero 4, 7 and 8.

### Ethical and legal aspects of the observations

According to Mauritius rules, observations were only performed during mornings (from 6.00 am to 12.00 pm). Out of respect for the cetaceans and their habitats, the observers strictly followed the ethical rules of the official Charter for responsible approach and observation of marine mammals and the Maritime zone regulations (Conduct of Marine Scientific Research/ Notice n°57 of 2017) promulgated by the Mauritius Government. This study was placed under the policies of the Mauritius Department for continental shelf, maritime zone administration and exploration, with appropriate permits to conduct underwater videos, underwater observations on sperm whales and marine scientific research.

### Underwater observations

The observation protocol was described in Sarano et al. (2021). Briefly, when a group of sperm whales was spotted from the boat, the animals were approached no closer than 100m and a small group of swimming observers, generally a scuba diver and 4 snorkelers, immerged themselves, upstream considering the movement direction of the sperm whale group. Observers were as passive as possible, typically not swimming towards the whales but waiting for the sperm whales to approach to film them. When sperm whales were static (e.g., socializing or sleeping), observers slowly and quietly approached. The scuba diver recorded videos and observations at a maximum diving depth of 40m, while the snorkelers performed observations from the surface and filmed the sperm whales at a maximum 20m depth.

The duration of observation varied between 20s to 10min when the animals were sleeping or socializing near the observers. The boat always stayed away and picked up the observers once the sperm whale group had moved away.

### Video processing

The identification of morphological markers to create the catalogue of ID-cards was based on meticulous analyses of the videos using VLC player (VideoLAN Organization, France). Slow motion mode was used to get the best screenshot for each of the body marks. These pictures were then used to illustrate the morphological markers on the catalogue.

### Morphological markers

The morphological markers retained for the ID-cards are illustrated in Figure 1. They include: sex, white spots, cuts with removal of material, scars from teeth marks (*i*.*e*. rake marks), shape of the fluke. Some of these marks can be observed from a boat (e.g., cuts on the caudal fin, cuts / callus on the dorsal fin), but the majority are visible only underwater (e.g., sex, cutting of the pectoral fins, clear spots of depigmentation on the ventral side, on the mandibular area and the cheeks, shape of the jaw, size of the teeth).

**Figure 1.**
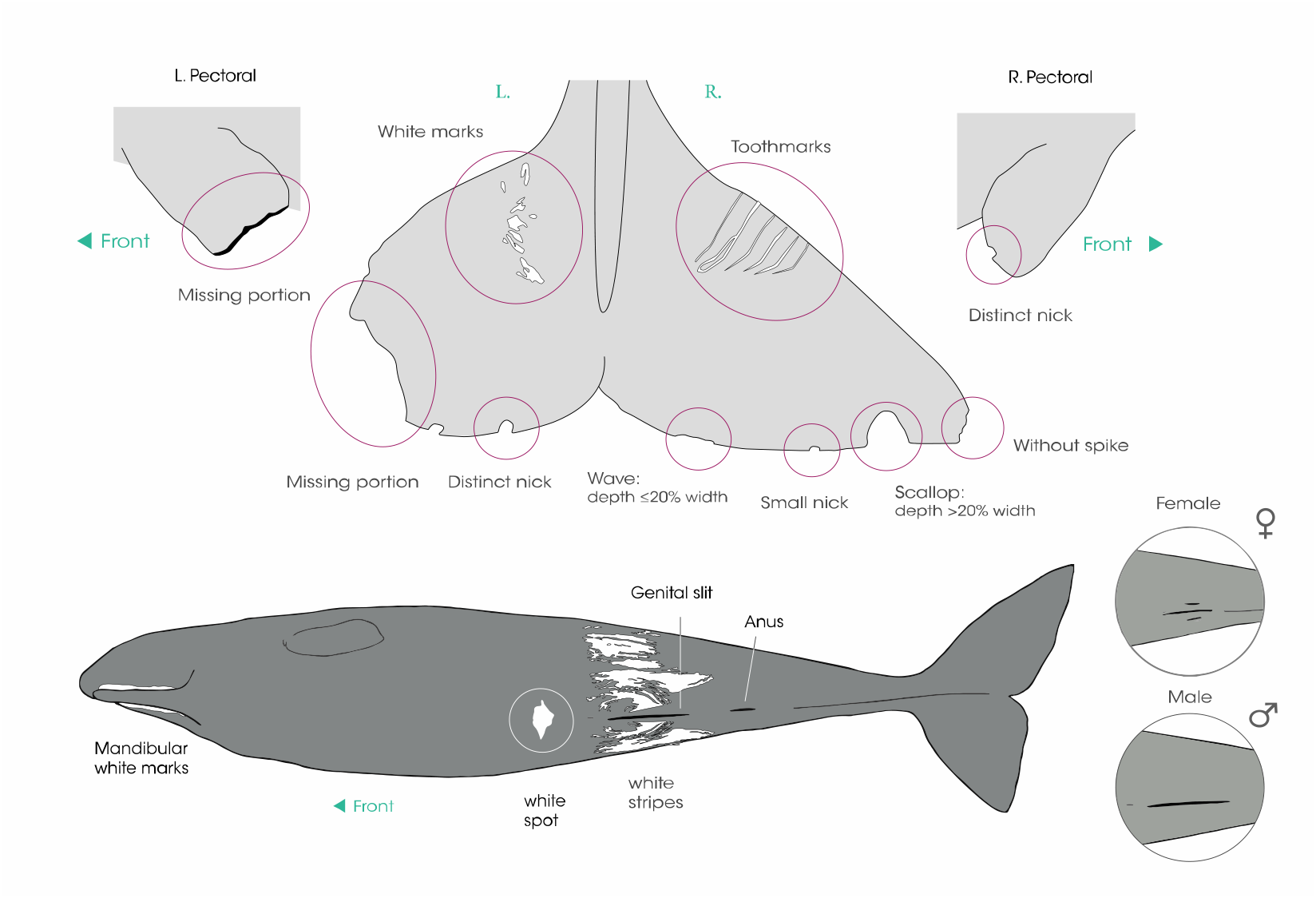
Morphological markers (sex, cutting patterns of the fins and depigmentation marks) used in this study to identify sperm whales.

#### Cutting Pattern of the fins

Types of cutting pattern of the fins (small nicks, distinct nicks, waves, scallops, missing portions, holes, tooth mark scars and calluses) have already been defined (Arnbom & Whitehead 1989, Whitehead 1990).

The features used in this study are (for those previously defined, descriptions are those of Arnbom & Whitehead 1989 when indicated; in the other cases, descriptions have been generalized to fit to underwater observations, Figure 1 and Table 2):

– Small nick: small indentation in edge of fin; only distinguished when the fin was relatively close (Arnbom & Whitehead 1989)
– Distinct nick: larger indentation sharply cut away (Arnbom & Whitehead 1989), which can be seen from a longer distance
– Wave: shallow smooth depression, with material removal, the depth of the missing part of the fin is ≤20% of its width
– Scallop: deep smooth depression, with removal of material, with depth of the missing part of the fin being ≥ 20% of its width
– Tip-missing: when only the tip of the fin is missing (fluke and pectoral)
– Missing portion: large part of the fin is sectioned (fluke and pectoral)
– Hole: small perforation of the fins
– Tooth mark: often seen as parallel scars
– Curled: tip of the fluke curled
– Callus: greyish or white deformity on the dorsal fin (Arnbom & Whitehead 1989)

**Table 2:**
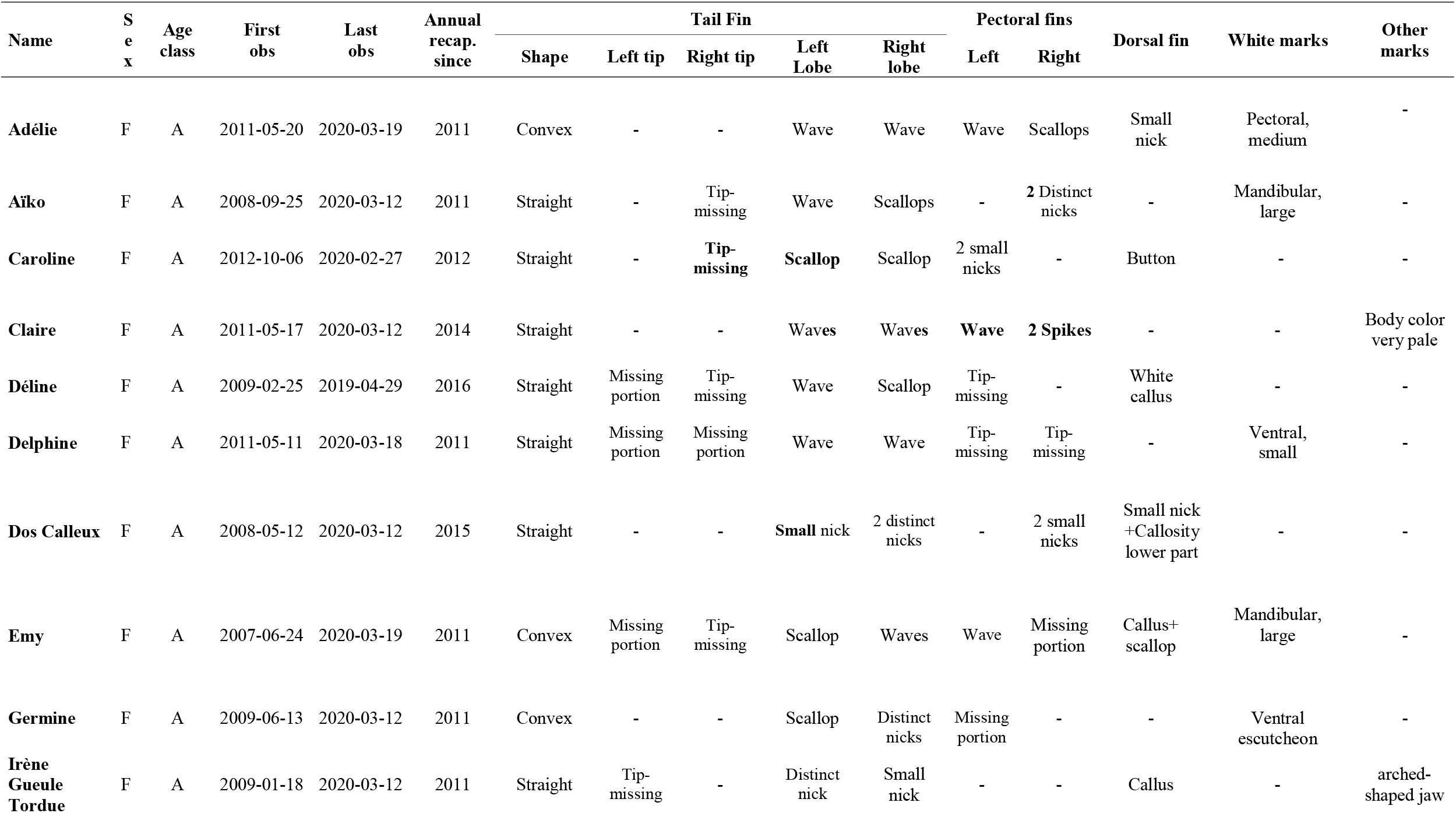
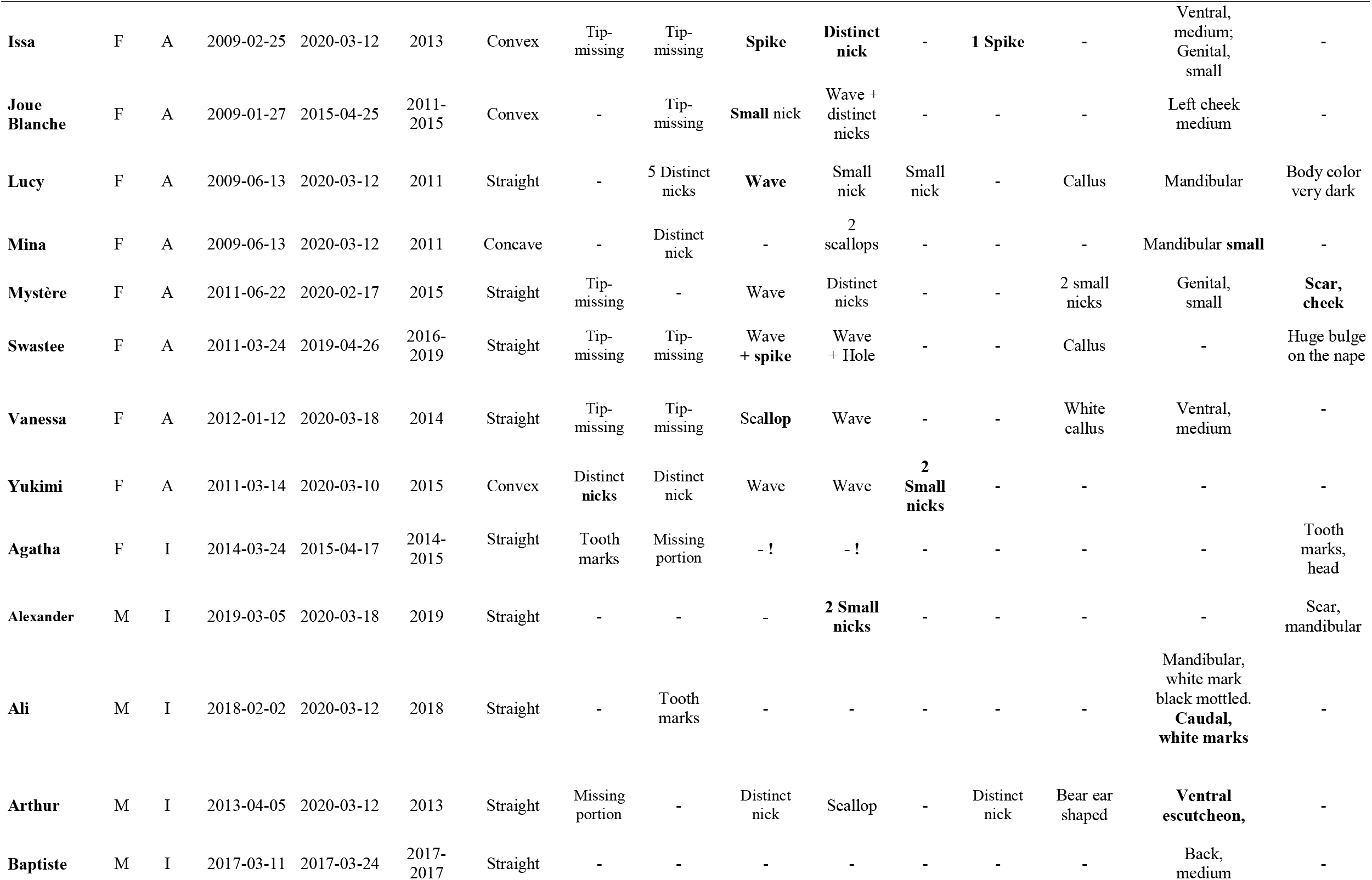
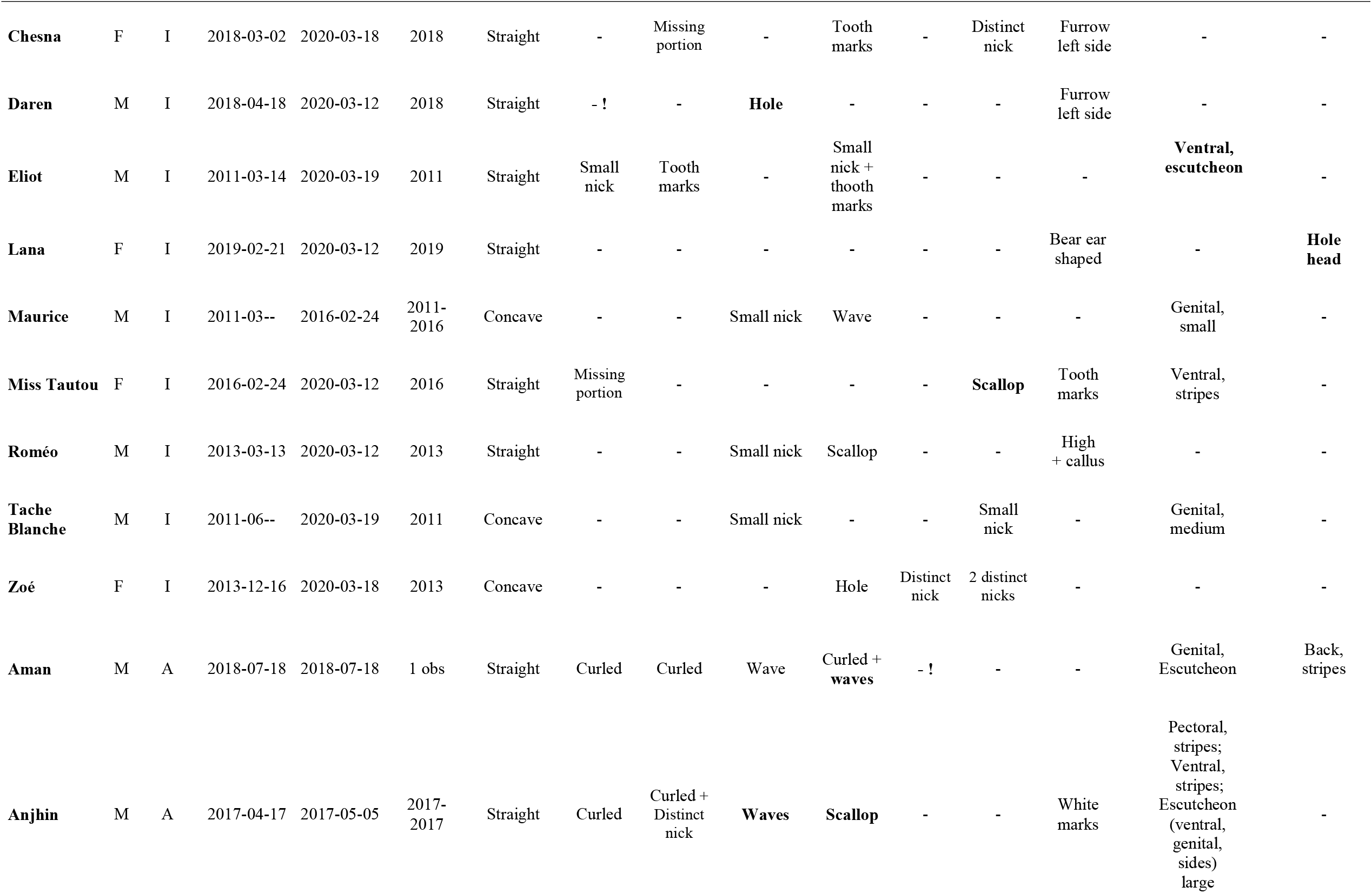
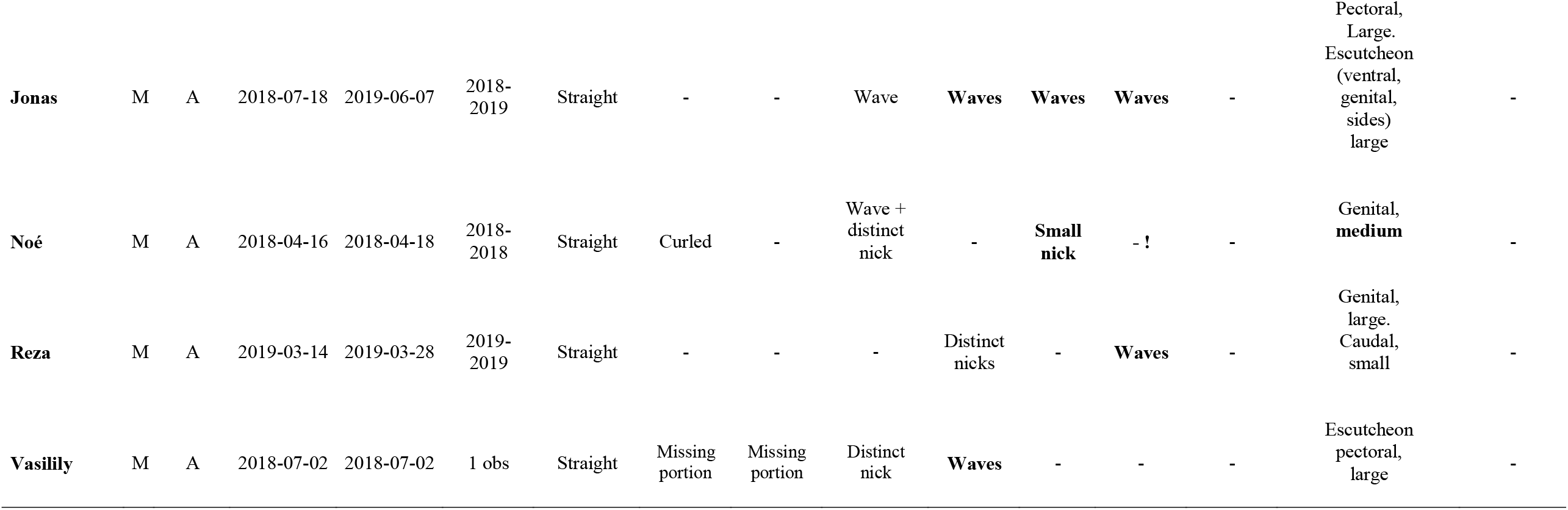
Marks used to identify specifically all the individual sperm whales represented in this study. A: Adult. I: Immature

| Name | Sex | Age class | First obs | Last obs | Annual recap. since | Tail Fin |  |  |  |  | Pectoral fins |  | Dorsal fin | White marks | Other marks |
| --- | --- | --- | --- | --- | --- | --- | --- | --- | --- | --- | --- | --- | --- | --- | --- |
|  |  |  |  |  |  | Shape | Left tip | Right tip | Left Lobe | Right lobe | Left | Right |  |  |  |
| **Adélie** | F | A | 2011-05-20 | 2020-03-19 | 2011 | Convex | - | - | Wave | Wave | Wave | Scallops | Small nick | Pectoral, medium | - |
| **Aïko** | F | A | 2008-09-25 | 2020-03-12 | 2011 | Straight | - | Tip-missing | Wave | Scallops | - | 2 Distinct nicks | - | Mandibular, large | - |
| **Caroline** | F | A | 2012-10-06 | 2020-02-27 | 2012 | Straight | - | **Tip-missing** | **Scallop** | Scallop | 2 small nicks | - | Button | - | - |
| **Claire** | F | A | 2011-05-17 | 2020-03-12 | 2014 | Straight | - | - | Waves | Waves | **Wave** | **2 Spikes** | - | - | Body color very pale |
| **Déline** | F | A | 2009-02-25 | 2019-04-29 | 2016 | Straight | Missing portion | Tip-missing | Wave | Scallop | Tip-missing | - | White callus | - | - |
| **Delphine** | F | A | 2011-05-11 | 2020-03-18 | 2011 | Straight | Missing portion | Missing portion | Wave | Wave | Tip-missing | Tip-missing | - | Ventral, small | - |
| **Dos Calleux** | F | A | 2008-05-12 | 2020-03-12 | 2015 | Straight | - | - | **Small** nick | 2 distinct nicks | - | 2 small nicks | Small nick +Callosity lower part | - | - |
| **Emy** | F | A | 2007-06-24 | 2020-03-19 | 2011 | Convex | Missing portion | Tip-missing | Scallop | Waves | Wave | Missing portion | Callus+scallop | Mandibular, large | - |
| **Germine** | F | A | 2009-06-13 | 2020-03-12 | 2011 | Convex | - | - | Scallop | Distinct nicks | Missing portion | - | - | Ventral escutcheon | - |
| **Irène Gueule Tordue** | F | A | 2009-01-18 | 2020-03-12 | 2011 | Straight | Tip-missing | - | Distinct nick | Small nick | - | - | Callus | - | arched-shaped jaw |
| **Issa** | F | A | 2009-02-25 | 2020-03-12 | 2013 | Convex | Tip-missing | Tip-missing | **Spike** | **Distinct nick** | - | **1 Spike** | - | Ventral, medium; Genital, small | - |
| **Joue Blanche** | F | A | 2009-01-27 | 2015-04-25 | 2011-2015 | Convex | - | Tip-missing | **Small nick** | Wave + distinct nicks | - | - | - | Left cheek medium | - |
| **Lucy** | F | A | 2009-06-13 | 2020-03-12 | 2011 | Straight | - | 5 Distinct nicks | **Wave** | Small nick | Small nick | - | Callus | Mandibular | Body color very dark |
| **Mina** | F | A | 2009-06-13 | 2020-03-12 | 2011 | Concave | - | Distinct nick | - | 2 scallops | - | - | - | Mandibular **small** | - |
| **Mystère** | F | A | 2011-06-22 | 2020-02-17 | 2015 | Straight | Tip-missing | - | Wave | Distinct nicks | - | - | 2 small nicks | Genital, small | **Scar, cheek** |
| **Swastee** | F | A | 2011-03-24 | 2019-04-26 | 2016-2019 | Straight | Tip-missing | Tip-missing | Wave + **spike** | Wave + Hole | - | - | Callus | - | Huge bulge on the nape |
| **Vanessa** | F | A | 2012-01-12 | 2020-03-18 | 2014 | Straight | Tip-missing | Tip-missing | **Scallop** | Wave | - | - | White callus | Ventral, medium | - |
| **Yukimi** | F | A | 2011-03-14 | 2020-03-10 | 2015 | Convex | Distinct **nicks** | Distinct nick | Wave | Wave | **2 Small nicks** | - | - | - | - |
| **Agatha** | F | I | 2014-03-24 | 2015-04-17 | 2014-2015 | Straight | Tooth marks | Missing portion | - ! | - ! | - | - | - | - | Tooth marks, head |
| **Alexander** | M | I | 2019-03-05 | 2020-03-18 | 2019 | Straight | - | - | - | **2 Small nicks** | - | - | - | - | Scar, mandibular |
| **Ali** | M | I | 2018-02-02 | 2020-03-12 | 2018 | Straight | - | Tooth marks | - | - | - | - | - | Mandibular, white mark black mottled. **Caudal, white marks** | - |
| **Arthur** | M | I | 2013-04-05 | 2020-03-12 | 2013 | Straight | Missing portion | - | Distinct nick | Scallop | - | Distinct nick | Bear ear shaped | **Ventral escutcheon,** | - |
| **Baptiste** | M | I | 2017-03-11 | 2017-03-24 | 2017-2017 | Straight | - | - | - | - | - | - | - | Back, medium | - |
| **Chesna** | F | I | 2018-03-02 | 2020-03-18 | 2018 | Straight | - | Missing portion | - | Tooth marks | - | Distinct nick | Furrow left side | - | - |
| **Daren** | M | I | 2018-04-18 | 2020-03-12 | 2018 | Straight | - ! | - | **Hole** | - | - | - | Furrow left side | - | - |
| **Eliot** | M | I | 2011-03-14 | 2020-03-19 | 2011 | Straight | Small nick | Tooth marks | - | Small nick + thooth marks | - | - | - | **Ventral, escutcheon** | - |
| **Lana** | F | I | 2019-02-21 | 2020-03-12 | 2019 | Straight | - | - | - | - | - | - | Bear ear shaped | - | **Hole head** |
| **Maurice** | M | I | 2011-03-- | 2016-02-24 | 2011-2016 | Concave | - | - | Small nick | Wave | - | - | - | Genital, small | - |
| **Miss Tautou** | F | I | 2016-02-24 | 2020-03-12 | 2016 | Straight | Missing portion | - | - | - | - | **Scallop** | Tooth marks | Ventral, stripes | - |
| **Roméo** | M | I | 2013-03-13 | 2020-03-12 | 2013 | Straight | - | - | Small nick | Scallop | - | - | High + callus | - | - |
| **Tache Blanche** | M | I | 2011-06-- | 2020-03-19 | 2011 | Concave | - | - | Small nick | - | - | Small nick | - | Genital, medium | - |
| **Zoé** | F | I | 2013-12-16 | 2020-03-18 | 2013 | Concave | - | - | - | Hole | Distinct nick | 2 distinct nicks | - | - | - |
| **Aman** | M | A | 2018-07-18 | 2018-07-18 | 1 obs | Straight | Curled | Curled | Wave | Curled + **waves** | - ! | - | - | Genital, Escutcheon | Back, stripes |
| **Anjhin** | M | A | 2017-04-17 | 2017-05-05 | 2017-2017 | Straight | Curled | Curled + Distinct nick | **Waves** | **Scallop** | - | - | White marks | Pectoral, stripes; Ventral, stripes; Escutcheon (ventral, genital, sides) large | - |
| **Jonas** | M | A | 2018-07-18 | 2019-06-07 | 2018-2019 | Straight | - | - | Wave | **Waves** | **Waves** | **Waves** | - | Pectoral, Large. Escutcheon (ventral, genital, sides) large | - |
| **Noé** | M | A | 2018-04-16 | 2018-04-18 | 2018-2018 | Straight | Curled | - | Wave + distinct nick | - | **Small nick** | - ! | - | Genital, **medium** | - |
| **Reza** | M | A | 2019-03-14 | 2019-03-28 | 2019-2019 | Straight | - | - | - | Distinct nicks | - | **Waves** | - | Genital, large. Caudal, small | - |
| **Vasilily** | M | A | 2018-07-02 | 2018-07-02 | 1 obs | Straight | Missing portion | Missing portion | Distinct nick | **Waves** | - | - | - | Escutcheon pectoral, large | - |

#### Skin depigmentation marks

The depigmentation of the skin in sperm whales result in white areas on the body that can be characterized according to their size (small, medium, large), their shape (spot, stripe, escutcheon) and their position on the body (caudal, genital, ventral, pectoral and mandibular areas, side and back). Finally, the presence of callus, fold or button on the dorsal fin are noted, as well as characteristics, rare but very discriminating, such as the crooked jaw or the bulge of the neck.

Temporary scratches and peeling spots were used only to help with daily recapture over a field season. These non-permanent markers were therefore, not retained in the catalog.

## Catalogue of ID-card creation

The catalogue developed through the underwater observation protocol consists of a series of individual ID cards for each sperm whale (Supplementary Information 1 and 2). For better recognition of individuals in the field and easier use of the catalogue, the ID-cards were designed using simplified standards (see Supplementary Information 1 and 2). For each individual, the distinctive markers were indicated on the ID-card (e.g., on the caudal), and/or detailed on dedicated zoomed photos (of pectoral, spots, mouth, …). Additional information was listed at the top of the card such as date of first observation, date of last observation, and years of successive observations. Each individual was given a name in an alphanumeric reference system to ease its identification in the field. Additional information, such as the availability of DNA samples, or information of kinship relations when known are enriched the ID-cards.

ID-cards were (and are) updated yearly with new elements in order to: (1) add new morphological markers, (2) take into account both the evolution over time of the markers, the growth and the presence of teeth, and (3) include any new information.

The number of recaptures for an individual is defined as the number of days that the individual is observed and filmed. Multiple daily resightings were ignored. This number is available for all individuals between 2011 and 2020 (Table 3).

**Table 3:** Numbers of days of observation per individual and percentage of days of observation per individual and per number of days of field work with sperm whale observation. **1st obs:** date of the first observation of each individual; **N**: number of days of observation of each individual per year; **%**: number of days of observation of each individual per number of days of field work with sperm whales observation; **Total 11-20**: total number of observations of the individual during 2011-2020 period; **Total 15-20**: total number of observations of the individual during 2015-2020 period. **Average**: average of % of observation per categories. For the missing individuals, days of last observations are indicated.

| Adult females | 1st OBS | 2011 | 2012 | 2013 | 2014 | 2015 |  | 2016 |  | 2017 |  | 2018 |  | 2019 |  | 2020 |  | Total 2015-2020 | Total 2011-2020 |
| --- | --- | --- | --- | --- | --- | --- | --- | --- | --- | --- | --- | --- | --- | --- | --- | --- | --- | --- | --- |
|  |  | N | N | N | N | N | % | N | % | N | % | N | % | N | % | N | % |  |  |
| Adélie | 2011 | 4 | 0 | 4 | 3 | 11 | 0.38 | 13 | 0.38 | 22 | 0.44 | 20 | 0.31 | 24 | 0.35 | 5 | 0.24 | 95 | 106 |
| Aïko | 2008 | 3 | 1 | 4 | 2 | 13 | 0.45 | 14 | 0.41 | 12 | 0.24 | 16 | 0.25 | 26 | 0.38 | 8 | 0.38 | 89 | 99 |
| Caroline | 2012 | 0 | 1 | 3 | 1 | 3 | 0.10 | 12 | 0.35 | 18 | 0.36 | 18 | 0.28 | 18 | 0.26 | 4 | 0.19 | 73 | 78 |
| Claire | 2011 | 1 | 1 | 0 | 1 | 6 | 0.21 | 13 | 0.38 | 12 | 0.24 | 15 | 0.23 | 15 | 0.22 | 5 | 0.24 | 66 | 69 |
| Déline | 2009 | 3 | 0 | 1 | 1 | 3 | 0.10 | 8 | 0.24 | 8 | 0.16 | 6 | 0.09 | 1 | 0.01 | 0 | 0.00 | 26 | 31 |
| Delphine | 2011 | 3 | 1 | 4 | 4 | 9 | 0.31 | 18 | 0.53 | 29 | 0.58 | 35 | 0.55 | 19 | 0.28 | 4 | 0.19 | 114 | 126 |
| Dos Calleux | 2008 | 1 | 0 | 0 | 0 | 2 | 0.07 | 7 | 0.21 | 20 | 0.40 | 14 | 0.22 | 15 | 0.22 | 8 | 0.38 | 66 | 67 |
| Emy | 2007 | 5 | 2 | 1 | 1 | 7 | 0.24 | 16 | 0.47 | 21 | 0.42 | 22 | 0.34 | 19 | 0.28 | 3 | 0.14 | 88 | 97 |
| Germine | 2009 | 10 | 5 | 4 | 4 | 23 | 0.79 | 21 | 0.62 | 31 | 0.62 | 32 | 0.50 | 36 | 0.53 | 11 | 0.52 | 154 | 177 |
| Irène | 2009 | 9 | 3 | 3 | 5 | 9 | 0.31 | 24 | 0.71 | 30 | 0.60 | 26 | 0.41 | 38 | 0.56 | 9 | 0.43 | 136 | 156 |
| Issa | 2009 | 1 | 0 | 1 | 3 | 3 | 0.10 | 13 | 0.38 | 10 | 0.20 | 14 | 0.22 | 14 | 0.21 | 5 | 0.24 | 59 | 64 |
| Lucy | 2009 | 4 | 1 | 3 | 3 | 11 | 0.38 | 23 | 0.68 | 18 | 0.36 | 27 | 0.42 | 21 | 0.31 | 4 | 0.19 | 104 | 115 |
| Mina | 2009 | 2 | 0 | 1 | 3 | 5 | 0.17 | 13 | 0.38 | 16 | 0.32 | 31 | 0.48 | 18 | 0.26 | 3 | 0.14 | 86 | 92 |
| Mystère | 2011 | 3 | 1 | 0 | 2 | 6 | 0.21 | 8 | 0.24 | 11 | 0.22 | 21 | 0.33 | 17 | 0.25 | 2 | 0.10 | 65 | 71 |
| Swastee | 2011 | 1 | 0 | 0 | 1 | 3 | 0.10 | 2 | 0.06 | 6 | 0.12 | 4 | 0.06 | 6 | 0.09 | 0 | 0.00 | 21 | 23 |
| Vanessa | 2012 | 0 | 1 | 1 | 4 | 5 | 0.17 | 11 | 0.32 | 26 | 0.52 | 33 | 0.52 | 14 | 0.21 | 3 | 0.14 | 92 | 98 |
| Yukimi | 2011 | 3 | 0 | 1 | 1 | 1 | 0.03 | 12 | 0.35 | 5 | 0.10 | 17 | 0.27 | 30 | 0.44 | 3 | 0.14 | 68 | 73 |
| **Averages** |  |  |  |  |  |  | **0.24** |  | **0.39** |  | **0.35** |  | **0.32** |  | **0.29** |  | **0.22** | **0.30** |  |
| Immatures | 1st OBS | N | N | N | N | N | % | N | % | N | % | N | % | N | % | N | % |  |  |
| Alexander | 2019 |  |  |  |  |  |  |  |  |  |  |  |  | 38 | 0.56 | 7 | 0.33 | 45 | 45 |
| Ali | 2018 |  |  |  |  |  |  |  |  |  |  | 47 | 0.73 | 45 | 0.66 | 11 | 0.52 | 103 | 103 |
| Arthur | 2013 |  |  | 3 | 6 | 21 | 0.72 | 29 | 0.85 | 30 | 0.60 | 41 | 0.64 | 43 | 0.63 | 11 | 0.52 | 175 | 184 |
| Chesna | 2018 |  |  |  |  |  |  |  |  |  |  | 39 | 0.61 | 31 | 0.46 | 6 | 0.29 | 76 | 76 |
| Daren | 2018 |  |  |  |  |  |  |  |  |  |  | 14 | 0.22 | 46 | 0.68 | 11 | 0.52 | 71 | 71 |
| Eliot | 2011 | 6 | 4 | 5 | 6 | 16 | 0.55 | 21 | 0.62 | 30 | 0.60 | 41 | 0.64 | 32 | 0.47 | 6 | 0.29 | 146 | 167 |
| Lana | 2019 |  |  |  |  |  |  |  |  |  |  |  |  | 43 | 0.63 | 6 | 0.29 | 49 | 49 |
| Miss Tautou | 2016 |  |  |  |  |  |  | 25 | 0.74 | 34 | 0.68 | 35 | 0.55 | 44 | 0.65 | 11 | 0.52 | 149 | 149 |
| Roméo | 2013 |  |  | 2 | 5 | 19 | 0.66 | 28 | 0.82 | 24 | 0.48 | 35 | 0.55 | 41 | 0.60 | 8 | 0.38 | 155 | 162 |
| Tache Blanche | 2011 | 3 | 3 | 5 | 5 | 11 | 0.38 | 15 | 0.44 | 25 | 0.50 | 41 | 0.64 | 32 | 0.47 | 7 | 0.33 | 131 | 147 |
| Zoé | 2013 |  |  | 1 | 4 | 12 | 0.41 | 21 | 0.62 | 23 | 0.46 | 31 | 0.48 | 27 | 0.40 | 6 | 0.29 | 120 | 125 |
| **Averages** |  |  |  |  |  |  | **0.54** |  | **0.68** |  | **0.55** |  | **0.56** |  | **0.56** |  | **0.39** | **0.55** |  |
| Missing individuals | 1st OBS | N | N | N | N | N | % | N | % | N | % | Last Obs |  |  |  |  |  |  |  |
| Joue Blanche | 2009 | 0 | 2 | 0 | 3 | 3 | 0.10 | - | - | - | - | 2015-04-25 |  |  |  |  |  | 3 | 8 |
| Agatha (Juv) | 2014 |  |  |  | 5 | 13 | 0.45 | - | - | - | - | 2015-04-17 |  |  |  |  |  | 13 | 18 |
| Baptiste (Juv) | 2017 |  |  |  |  |  |  |  |  | 9 | 0.18 | 2017-03-24 |  |  |  |  |  | 9 | 9 |
| Maurice (Juv) | 2011 | 1 | 4 | 3 | 6 | 15 | 0.52 | 1 | 0.03 | - | - | 2016-02-24 |  |  |  |  |  | 16 | 29 |

## RESULTS

Between 2015 and 2020, the team went out in the field on average 53 days per year (min 36d in 2015 and 2020, max 81d in 2019), mainly between February and May (Table 1). In 2020, the fieldwork season was shortened due to bad weather and the Covid19 pandemic.

The ID-cards were created from about 250 hours of underwater video recording between 2015 and 2020, for a total number of 317 days of observation (see Table 1). Sperm whales were observed in 83.9% of the fieldtrips.

### Catalogue of individual ID-cards

A total of 38 ID-cards corresponding to 38 identified individuals are presented in this study: 18 adult females, 14 immatures (9 males, 5 females) and 6 adult males (Table 2 and Supplementary Information 1 and 2). Table 2 presents all morphological markers identified for each of these 38 individuals. They are described according to their position on the body (e.g., sex, caudal, pectoral, dorsal, back, head) and, for white marks, according to their location on the ventral parts of the animal.

Gender was the first identification criterion used in the field to identify the individuals. Then the individual-specific body markers were used to narrow down the identification at the individual level. The number of body marks typically increased with age, young individuals displaying very few (e.g., Ali, Alexander, Daren, Lana) to more than 10 marks in older individuals (e.g., up to 14 marks for the adult male Anjhin, Table 2). Some of these body markers were unique enough to enable direct identification of the individuals: e.g., distinct missing portions on the fluke (e.g., Arthur, Chesna, Miss Tautou, Agatha) or on the pectoral fin (e.g., Germine), white markers (e.g., Adélie, Tache blanche, Issa, Joue Blanche) or arched-shaped jaw (e.g., Irène’ s twisted jaw). For other individuals, the observation of several body markers was required to make the identification. Overall, the body marks presented in Table 2 enabled field observers to unambiguously identify these 38 individuals.

### Marker persistence over time

All body markers used to draw the ID-cards were persistent over time, i.e., no body marker disappeared during the present study (*i*.*e*., 9 years): white skin pigmentation appeared stable over time as well as markers resulting from a wound with flesh removal: *e*.*g*., Eliot’s clear ventral escutcheon (Figure 2), the white spot of Tache Blanche, the sectioned pectoral of Germine or Irène’ s twisted jaw were recaptured on the videos, either from birth (for immature: Eliot and Tache Blanche), or since their first observations in 2011 (for adult females: Germine and Irène).

**Figure 2.**
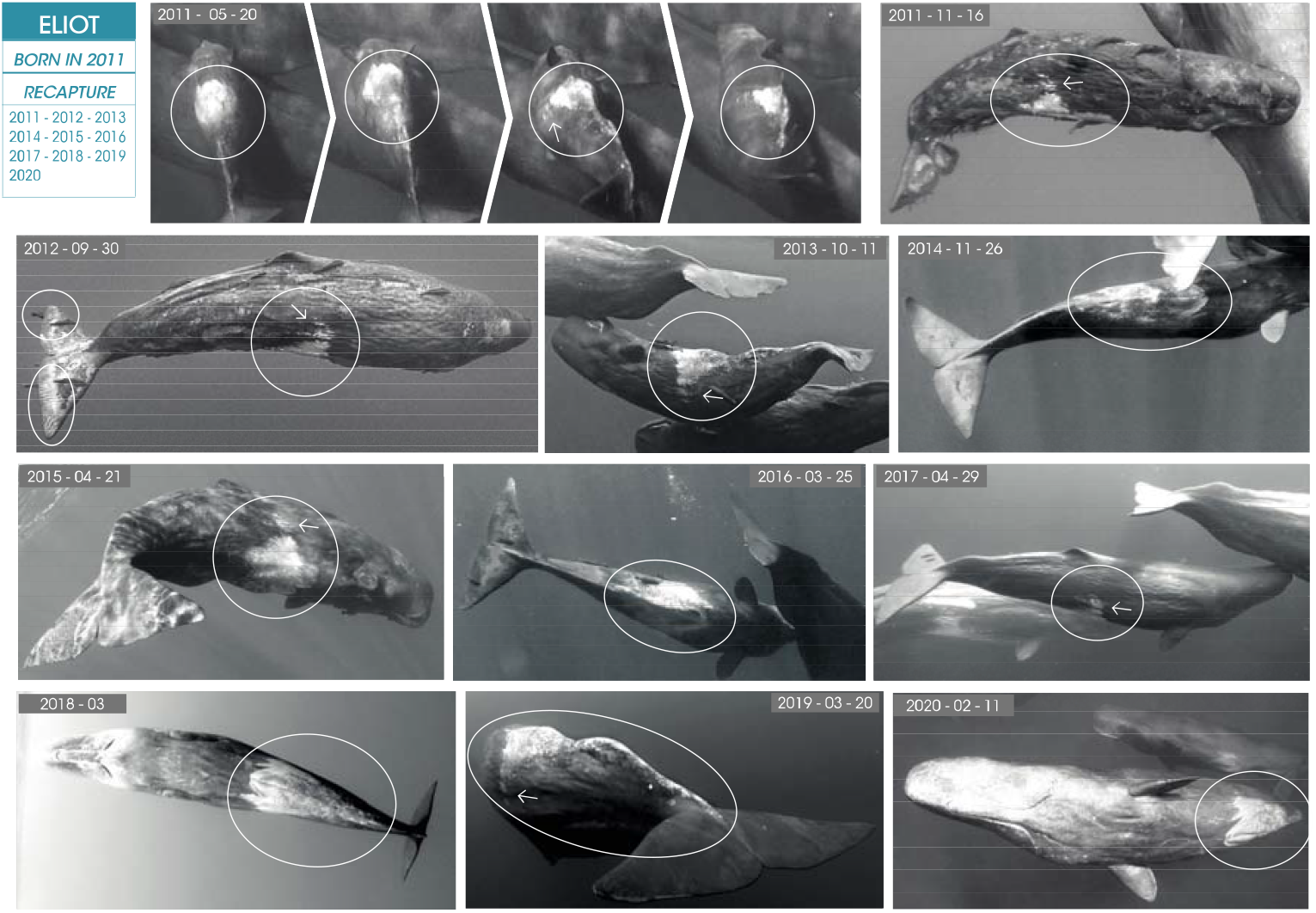
Example of recapture of a body marker over 9 years: the first 4 photos (taken from a sequence where the newborn Eliot turns around) show the shape of the escutcheon captured at different angles. The other photos were taken every year from its birth in 2011 until 2020. The escutcheon being unique, it allows the direct identification of this immature, while its caudal fin shows only traces of teeth and tiny notches almost indistinguishable.

### Recapture rate of females and immatures

The primary aim of the ID-card catalogue was to enable underwater field identification of individual whales from 2015 to 2020. The ID-cards were also used to analyze field videos recorded between 2011 and 2014, as well as some older underwater photographs taken in 2007 and 2009, in order to identify the individuals. The observation effort was therefore divided in 2 periods, one until 2014 and the second starting in 2015 (Tables 1 and 3).

Over the period 2011-2020, 17 adult females identified were recaptured 1,542 times with an average rate of 91 recaptures per individual (min=23, max=177). One adult female, Joue Blanche, was seen only 8 times, and no more since 2015 (Table 3). Among the adult females, 2 had few recaptures (Déline *n* = 31 and Swastee *n* = 23) although they were easily identified thanks to their distinct markers (for Swastee, a huge bulge on the nape; for Déline, a big cut on the caudal fin -see Supplementary Information 1 and 2). The most recaptured females were Germine (n = 177) and Irène (n=156), observed during more than 50% of the days of field work. Immatures were more often recaptured, even those presenting limited distinctive markers such as Roméo, Ali or Daren (between 39% to 68% of the days of observation depending on the individual, 55% on average) than adult females (between 22% to 39%, 30% of the days of observation on average). The most recaptured immature was Arthur (n=184).

During the 2015-2020 period, some individuals disappeared. They have been collated at the bottom of Table 3, i.e., 2 immature males (Maurice, 5 years-old and Baptiste, 3 weeks-old), probably dead, and a immature female (Agatha, 1 year-old) with her assigned mother Joue Blanche (observed since 2009) who both disappeared (or left the group) in April 2015.

## DISCUSSION

This study presents an underwater observation protocol based on morphological body markers in a sperm whale population off Mauritius that led to the development of a robust catalogue of ID-cards enabling the unique identification and monitoring of 38 sperm whales.

### Advantages of underwater monitoring

#### Direct gender assignation

Platform-based observers (e.g., on a boat) can hardly assign gender to immatures showing no global sexual dimorphism (Arnbom & Whitehead 1989, Gero et al. 2013). Adult females and large immatures have similar sizes, and are also often classified together (Matthews et al. 2001; Gero et al. 2014). Except for adult male sperm whales, easily identifiable (Arnbom & Whitehead 1989), skin biopsies and molecular sexing are therefore necessary to determine the genders (Gero et al. 2008, 2009, 2014). Underwater observation allows to observe the genital slit, and thus to distinguish between males and females, even before they reach sexual maturity. Here gender assignment was possible for 14 immatures, some of them from the day they were born.

#### Identification of immatures without any distinctive markers on the fluke

Underwater observation provides access to a range of body markers that a platform-based observer can only occasionally see but whose utility on sperm whale individual identification has been proved (Van der Linde & Eriksson 2020). These markers are, in particular, relevant for individuals without any distinctive markers on the fluke (Figure 3) and for the very young individuals that seldom fluke. These markers are, for instance, the indentations on the pectoral fins, the shape of the jaw or the pigmentation patterns on the ventral side, the flanks and the mandibular area. The presence / pattern of colored markings is often used for humpback whales (Glockner-Ferrari & Ferrari 1990) or for dolphins (Herzing 1997). Three immatures with an intact caudal fin and therefore impossible to identify from a boat were identified this way: Zoé, Tache blanche and Eliot (Figure 3). Underwater, the observer can notice that Zoé has a distinct nick on each pectoral and is a female, Eliot, a male, has a white ventral escutcheon and Tache blanche, another male, has a white spot on the belly as well as a small nick on the right pectoral (Figure 3).

**Figure 3.**
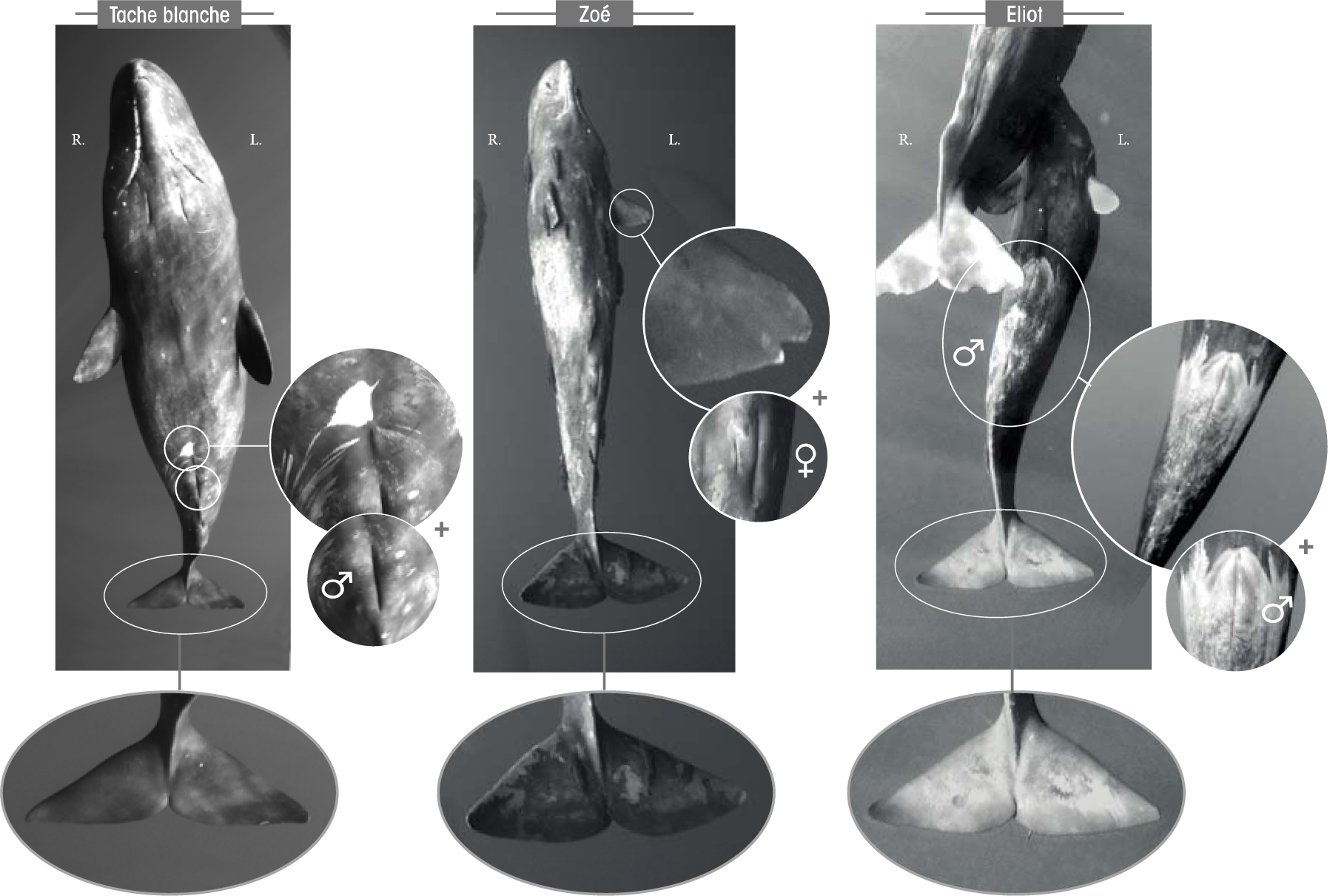
Differences between 3 immatures with intact caudal (from left to right): Tache Blanche has a white spot on the belly, a small nick on the right pectoral and is a male; Eliot has a white ventral escutcheon and is a male; Zoé has a nick on each pectoral and is a female.

This underwater method provided additional information on the immatures and could therefore improve the knowledge on this cohort, *e*.*g*., by allowing to determine mortality rates including calves (Gero & Whitehead 2016).

#### Recapture rate

The recapture rate is increased by underwater observation as compared to boat-based studies. Over a period of 9 years, the method presented here has resulted in numerous recaptures for all the individuals identified (mean rate for adult females = 91, Table 3). As a comparison, another sperm whale study in Mauritius, based on boat-observations, identified 101 different sperm whales among which 32 where sighted more than once over 5 years (Huijser et al. 2020). Another 28-year study in the West Indies identified 419 individuals, of which 175 individuals were recaptured 2 to 14 times (Gero et al. 2014).

#### Identification and behavioural observation of deeply-immerged individuals

Koyabashi et al. (2020) noticed that fluking was more often observed during foraging than after social interactions, which could lower the possibility of identifying individuals socializing thanks to fluke marks. Underwater studies enable to observe and capture several behaviours and social interactions that may be difficult to record from a boat, like underwater gathering (playing, socializing, swimming together), suckling (Johnson et al. 2010) or sleeping behaviour (figure 4).

**Figure 4.**
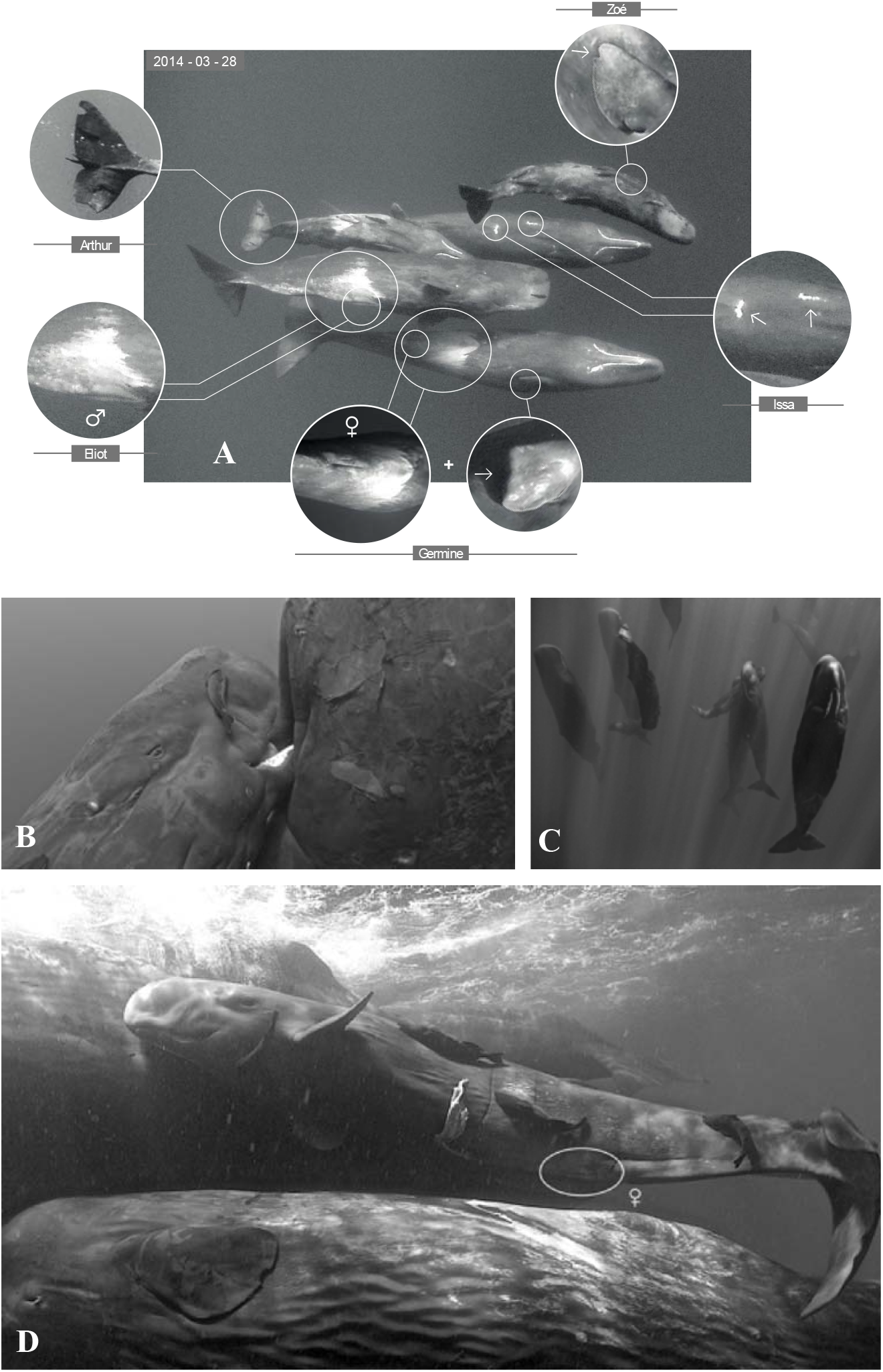
Different examples of behaviours and social interactions recorded underwater: **A**: gathering (note that all the individuals are recognized through body marks), **B**: suckling by the mouth (Johnson et al. 2010), **C**: sleeping, **D**: sex identification of a newborn.

#### Non-invasive sampling and individual acoustic signature

Some studies have used sloughed skin samples released by sperm whales as a source of non-invasive sampling for DNA analysis (Richard et al. 1996). These samples were taken from the sea surface. Underwater, and as the sperm whales can be identified in the field or later on video recording, skin samples can be taken in an individual specific manner by the snorkelers, allowing individual specific genotype determination, of first use for kin relationship determination for instance (Sarano et al. 2021).

Moreover, this visual identification of each individual may allow to perform individual specific recording, which is the key for research on individual acoustic signature. Current work joins video and audio labels from fine acoustic localization using high velocity hydrophone array recording (Ferrari et al. 2019).

### Disadvantages and limits of the underwater approach

#### Ethical and legal constraints

This method can only be used in areas where swim-with activities are legally allowed, ethically acceptable, and with appropriate permits from the Authorities. Our protocol implies that, once the snorkelers and scuba divers are in the water, the boat goes away. This also contributes to lower the human presence: other protocols, using on boat observations, involve that the boat follows the sperm whale until it flukes (Arnbom 1987). However, swimming regularly with marine mammals might impact their behaviour with a possible habituation or sensitization in the long term (Bejder et al. 2009): targeted animals tend to increase their avoidance behaviours (Constantine 2001, Delfour 2007, Filby et al. 2014), to change their activity budget and aerial behaviours (Peters et al. 2013) and to modify their sound productions (Scarpaci et al. 2000). However recent studies showed that the animals’ responses might be species-specific (Cecchetti et al. 2019, Pagel et al. 2017). Richter et al. (2006) showed an impact of whale-watching tour boats on sperm whales’ ventilation, vocalization patterns and swimming direction changes. The potential impacts of swim-with activities on sperm whales’ behaviours will have to be analyzed in the next years.

#### Water conditions

In this study, the underwater visibility (around 20m) enabled to easily identify morphological markers and white spot patterns on the body. But in terms of logistics linked to climatic conditions, it is clear that this underwater method cannot be implemented everywhere, e.g., it is much more complicated to perform underwater observations in polar waters for example, which are relatively dark and where the temperature may be near 0°C. Additional equipment adapted to these conditions would then be necessary. High turbidity can also reduce the visibility to a few meters (due to high primary production or turbid rainwater coming from inland). In those cases, this underwater method cannot be implemented, and only well-marked sperm whales (i.e. with white spots or large missing portions) are identifiable.

#### Identification of isolated individuals

When sperm whales are not grouped underwater, the identification of individuals which are not in the range of the camera is impossible because big-lense camera cannot be used underwater. However, several immersions, in compliance with the cetacean approach charter, when possible, can permit to overcome this issue in order to identify all individuals. Otherwise, underwater observations can be supplemented by observations from the boat.

## Importance of underwater observations for sperm whale conservation

Although many cetacean species are highly mobile, and show great dispersal capacities, their intraspecific diversity strongly vary, some species display local cultures and some populations may show high site fidelity (eg Gero & Whitehead 2016, Louis et al. 2017, Richard et al. 2018). Conservation priorities cannot then be defined at the species level, but rather at the population level (*e*.*g*., Clapham et al. 1999, Baker et al. 2013, Gero et al. 2016, Louis et al. 2017, Richard et al. 2018). Small-scale studies have therefore to be performed, taking into account and focusing on the local characteristics of the groups or populations. Such studies need to be able to estimate the level of differentiation of the studied group in the species, its connectivity with surrounding individuals and/or groups of the same species, the global health of the group, and its trends over years. The sperm whale is listed as vulnerable by the IUCN (Taylor et al. 2019). Whitehead (2002) estimated that sperm whale numbers have been reduced to about 32% of their original abundance by commercial whaling. The species was predicted to have recovered since the end of commercial whaling in 1986. But local trends have been shown to vary, to be locally slightly increasing (Moore & Barlow 2014) or not (Carrol et al. 2014), and to be worrying in some places (Reeves & Notarbartolo Di Sciara 2006, Gero & Whitehead 2016). Thanks to a long-term monitoring of well known social groups, Gero & Whitehead (2016) highlighted the disturbing situation for sperm whales in the West Indies. But the authors stress that these negative trends have been difficult to highlight, as immigration from surrounding regions may hide local mortality (Gero & Whitehead, 2016).

In the Indian Ocean, Kirkwood et al. (1980) estimated a global abundance of around 30,000 sperm whales, but no recent estimation is available. Anthropic activities are nowadays well known to negatively impact marine mammals in general (*eg* Jung & Madon *in press*), and sperm whales in particular (Gero and Whitehead 2016). For instance, collisions with ships (Laist et al. 2001) and ingestion of plastic debris (Jacobsen et al. 2010, de Stephanis et al. 2013, Unger et al. 2016) have demonstrated direct lethal effects on sperm whales. Marine debris accumulation has been recently evidenced in the Indian Ocean (Duhec et al. 2015, Lavers et al. 2019), as well as the direct impact of by-catch on cetaceans (Anderson et al. 2020). The expected recovery of sperm whales in the Indian Ocean needs thus to be carefully analyzed, and long-term localized monitoring of sperm whales are therefore strongly needed.

In conjunction with boat-based observation, allowing to identify high number of individuals (Huisjer et al. 2020), the underwater approach presented in this study will strongly help to determine the trends of the studied sperm whale populations. First, by increasing the accuracy and the frequency of individual identification, and the number of individual recaptures. Second, by bringing the opportunity to differentiate calves one from each other. Calves are in fact particularly important, as only their precise count allows to determine the real rate of increase of the population (e.g. Gero and Whitehead 2016). Local trends of sperm whale populations will be more precisely determined, which will be of first importance to define conservation concerns, priorities, and to measure the benefits of newly implemented protection plans. Other cetaceans could obviously strongly benefit of such careful individual specific studies, in particular when local groups of small size are known, whose trends can vary depending on impact of local anthropogenic activities (e.g. Louis et al. 2017).

## CONCLUSION

The protocol based on underwater videos has already proven to be highly robust and widely-used for other marine megafauna species (e.g., Glockner-Ferrari & Ferrari 1990, Herzing 1997, Marshall & Holmberg 2018, Pierce et al. 2018). It has been applied here for the identification of sperm whales in Mauritius, based on underwater observations. The relevance of this approach is evidenced by quasi-daily recaptures of females and immatures, over the field seasons and from one year to another. These recaptures were carried based on markers that can hardly be observed from the sea surface. The markers used proved to be stable and reliable over the 9 years of the study. This underwater observation approach using video recordings enables to identify individuals with intact caudal fins and to sex the entire group, including young and newborns, without using biopsies and molecular sexing. Like any catalogue, it requires annual updates of the ID-cards to take into account the possible evolution of morphological markers. It will also soon be extended by the ID-cards of around sixty more individuals observed off the coast of Mauritius.

## ACKNOWLEDGEMENT

Mauritian public authorities greatly helped the Maubydick project, in particular the Mauritian Prime Minister Office, the Marine Continental Shelf Exploration and Administration (MCSEA, Dr Réza Badal and his team), the Albion Fisheries Research Center (AFRC, Chief Scientific officer Mr Satish Kadhun), the Mauritius Film Development Corporation (MFDC, Mr. Sachin Jootun et Miss Eliana Timol) and the Tourism Authority (TA, Miss Khoudijah Boodoo, ex-Director). We thank Navin Rishinand Boodhonee and the Blue water diving center (Trou aux Biches, Mauritius), as well as all the ecovolunteers of MMCO and of the program Maubydick for their valuable participation in field work. A special thanks to late Cindy Vandebreucq for her involvement in 2013-2014 work. Stéphane Granzotto, Fabrice Guérin, Bernard Kirchhofer and Vanessa Mignon participated to the video recording and photography taking.

The Association Longitude 181 and Daniel Jouannet, Réseau-TERIA (Vitry sur Seine, France), supported the drawing and filling of the Identity cards.

## AUTHORISATIONS

To respect cetaceans and their habitats, the observers strictly followed the ethical rules of the official Charter for responsible approach and observation of marine mammals and the Maritimes zones regulations (Conduct of Marine Scientific Research/ Notice n°57 of 2017) promulgated by the Mauritius Government. This study was placed under the policies of the Mauritius Department for continental shelf, maritime zone administration and exploration, with appropriate permits to conduct underwater videos, underwater observations on sperm whales and marine scientific research on sperm whales.

