## Supplementary information 1 for "Underwater photo-identification of marine megafauna: an identity card catalogue of sperm whales (*Physeter macrocephalus*) off Mauritius Island"

### MALES ♂

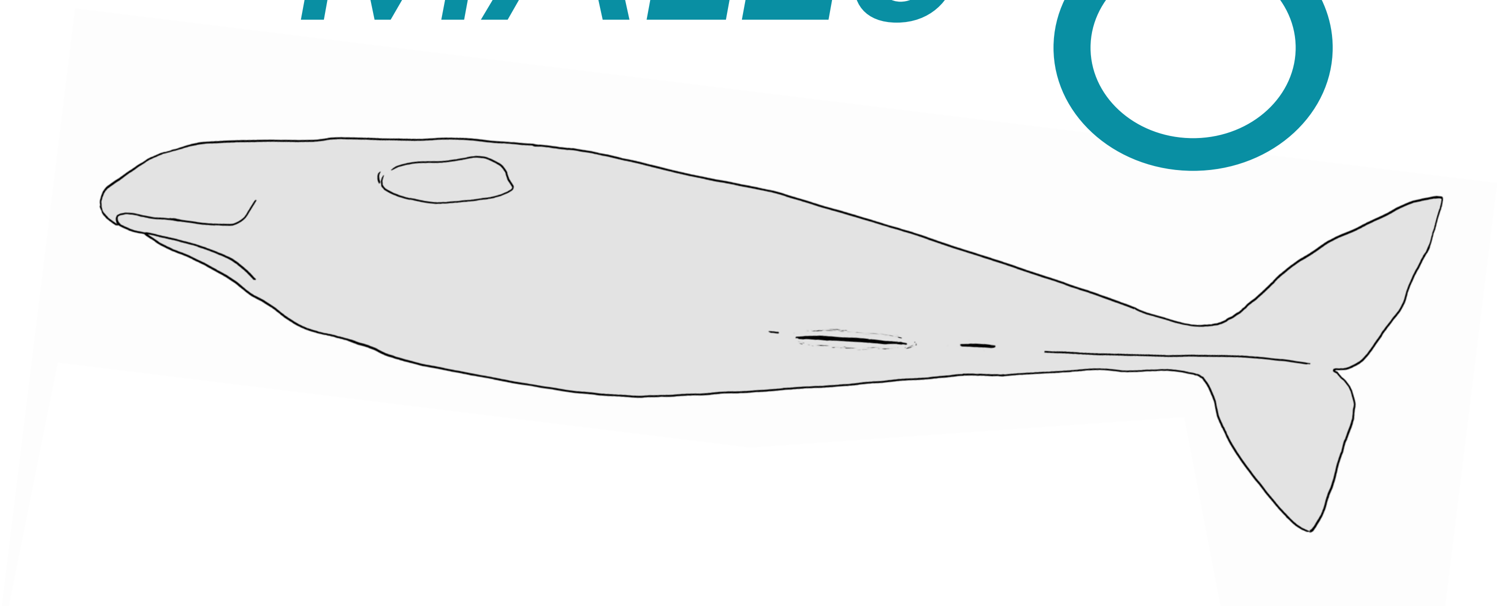

**CONCEPT** : François Sarano

**GRAPHIC DESIGN & ILLUSTRATIONS** : Marion Sarano

**PICTURES** : Stéphane Granzotto, Fabrice Guérin, René Heuzey, Daniel Jouannet, Bernard Kirchhofer, Vanessa Mignon, Axel Preud'homme, Véronique Sarano, François Sarano, Hugues Vitry.

© Longitude 181 - Creation 2015 / Update 2020

LONGITUDE 181
La Voix de l'Océan

IDENTITY CARDS
**SPERM WHALES OF MAURITIUS I.**

un océan de vie
www.unoceandevie.com
TERIA
Marine Megafauna Conservation Organisation

### AMAN

*1ST OBS. : 2018 - 07 - 18 // LAST OBS. : 2018 - 07 - 18*

| *ADULT* | *SEEN* | |
|---|---|---|
| *DNA* | ... - 2018 | *ADDITIONAL INFORMATION* |
| ♂ | | • Large white ventral escutcheon<br>• Curled R. & L. caudal tips |

L. pectoral

Aman

◀ Front

L. pectoral

◀ Front

2018 - 07 - 18

Ventral view // 2018 - 07 - 18

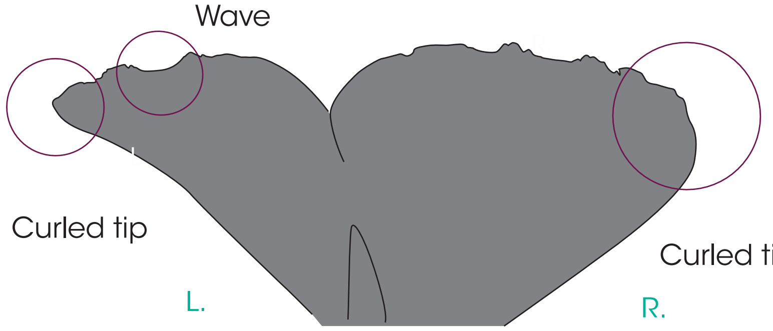

R. pectoral

**UNHARMED**

Front ▶

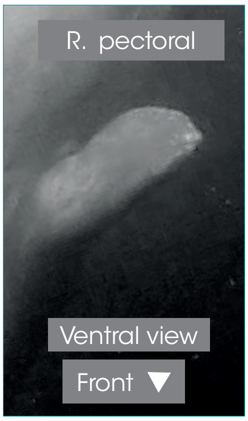

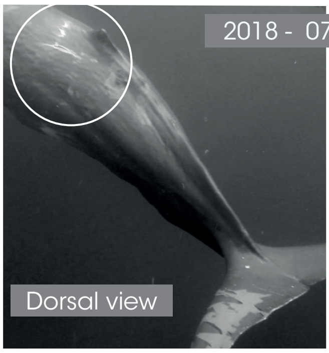

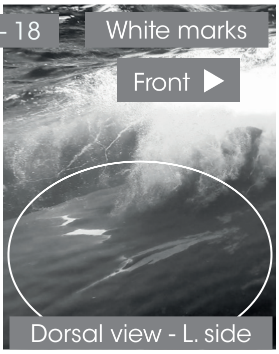

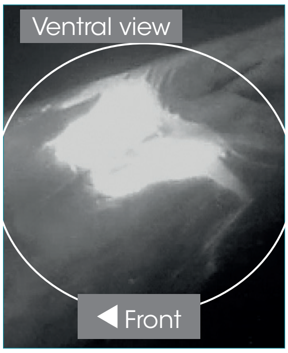

L. side

Ventral view

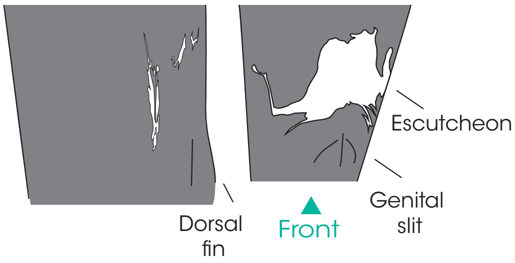

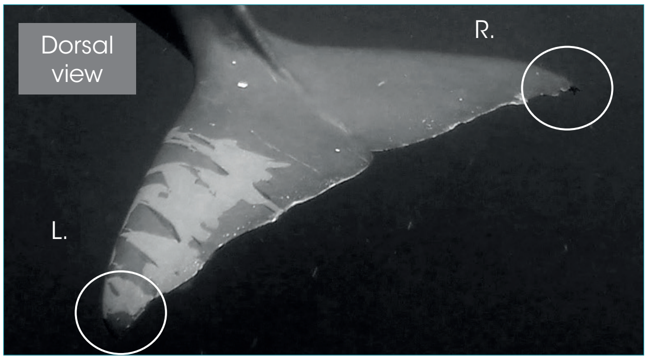

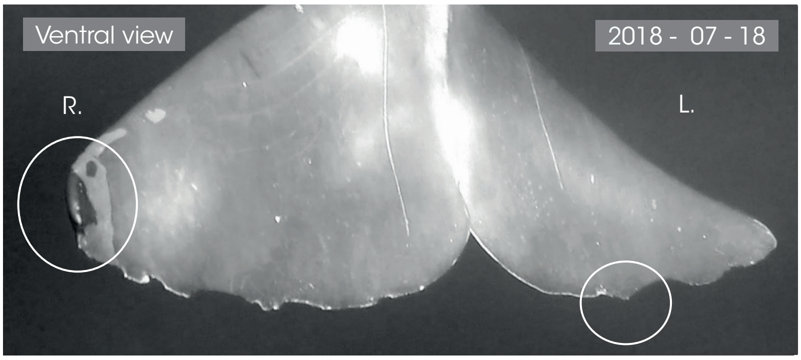

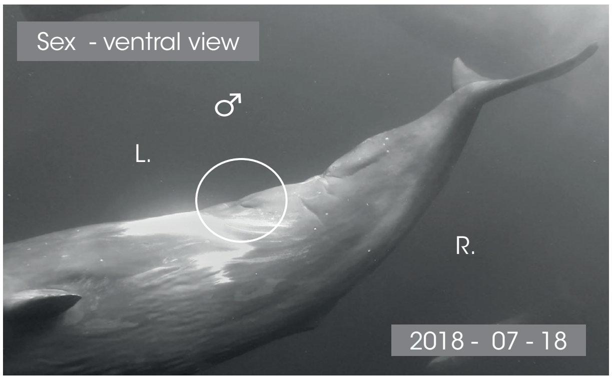

L. side

◀ Front

R. side

Front ▶

© Longitude 181 - Creation 2015 / Update 2020 - Concept : F. Sarano - Graphic design & illustrations : M. Sarano - © Pictures : S. Granzotto, F. Guérin, R. Heuzey, A. Preud'homme, V. & F. Sarano.

### ANJHIN

**1[ST] OBS. : 2017 - 04 - 17 // LAST OBS. : 2017 - 05 - 02**

| ADULT | SEEN | |
|---|---|---|
| DNA | 2017 | ADDITIONAL INFORMATION |
| ♂ | | • Many horizontal white stripes on the back and the sides, in the area of the dorsal.<br>• Curled R. & L. caudal tips. |

L. pectoral

**UNHARMED**

◀ Front

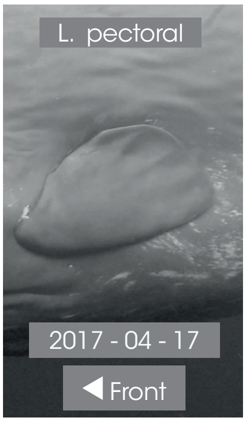

R. pectoral

**UNHARMED**

Front ▶

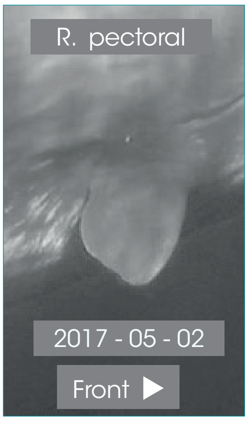

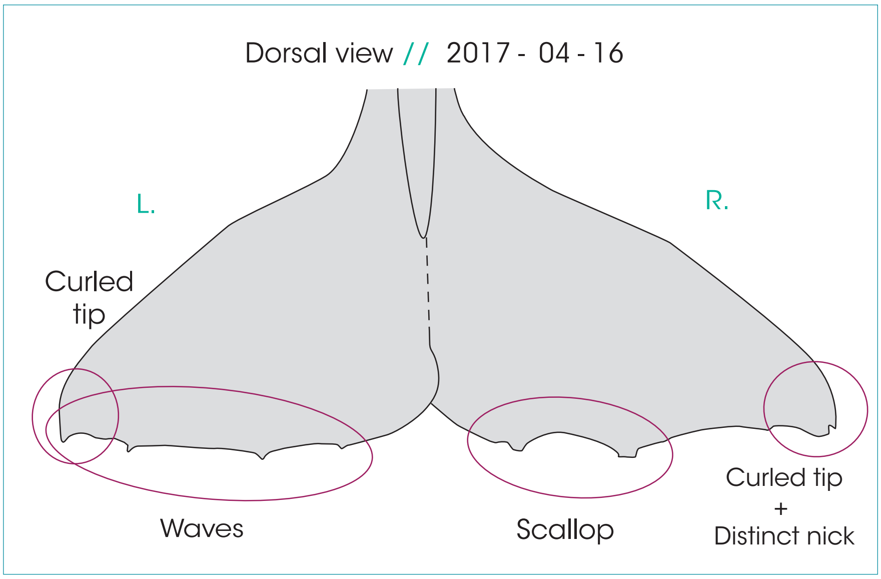

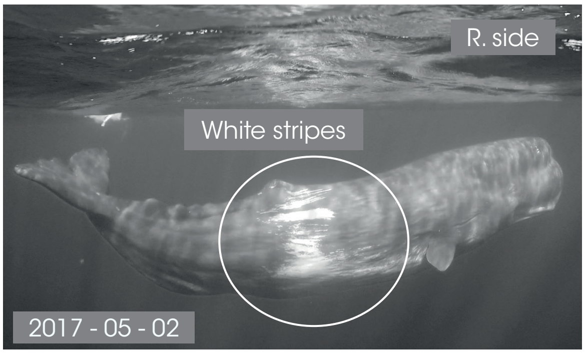

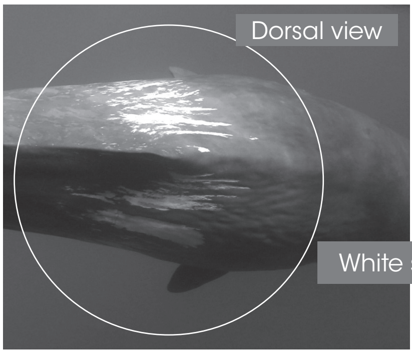

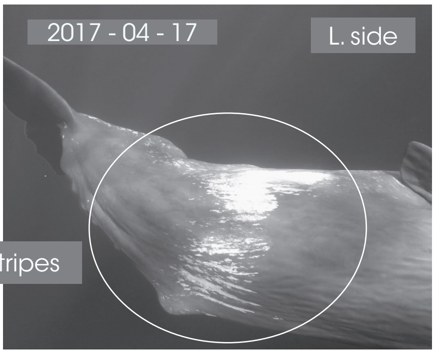

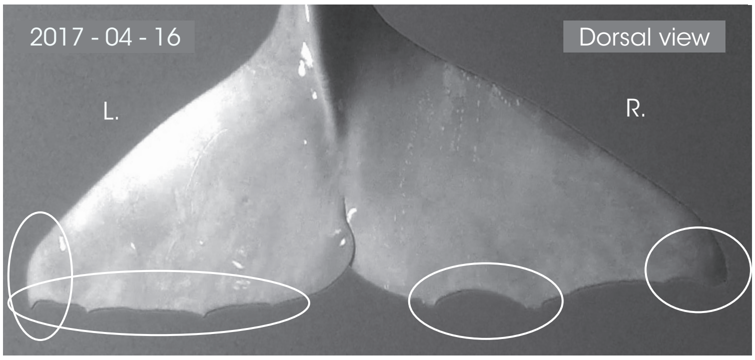

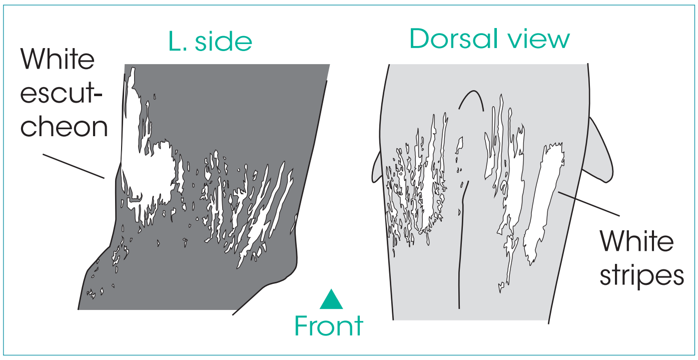

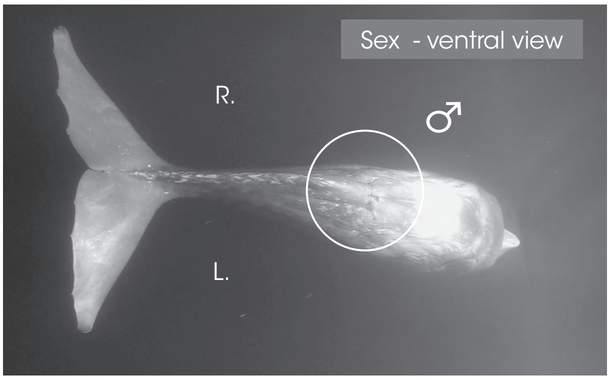

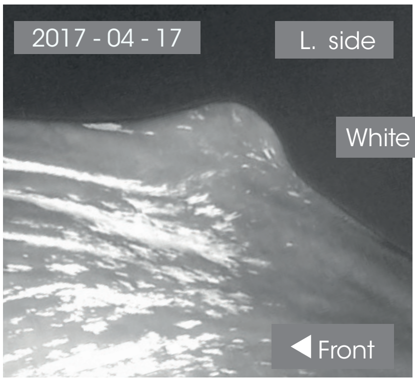

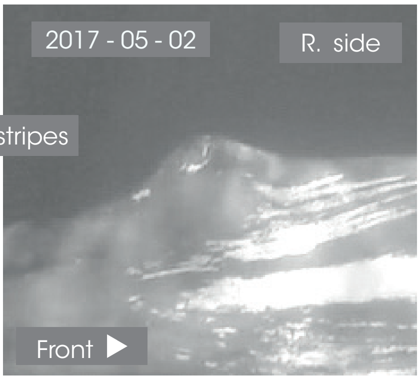

© Longitude 181 - Creation 2015 / Update 2020 - Concept : F. Sarano - Graphic design & illustrations : M. Sarano - © Pictures : S. Granzotto, F. Guérin, R. Heuzey, A. Preud'homme, V. & F. Sarano.

### JONAS

**1ST OBS. : 2018 - 07 - 18 // LAST OBS. : 2019 - 06 - 07**

| ADULT | SEEN | |
|---|---|---|
| DNA | 2018 - 2019 | ADDITIONAL INFORMATIONS |
| ♂ | | • Caroline's father, Zoé & Alexander's grandfather.<br>• Daren's father |

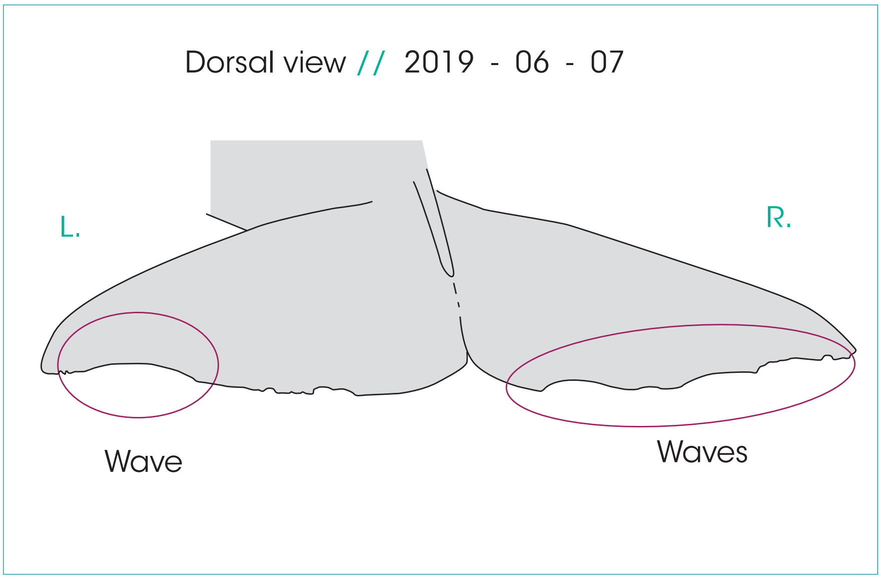

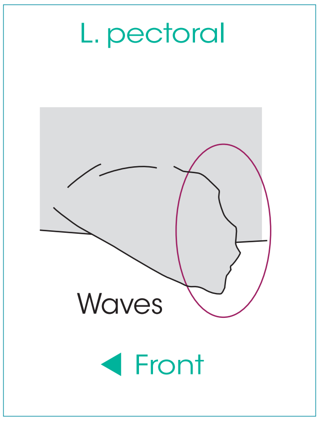

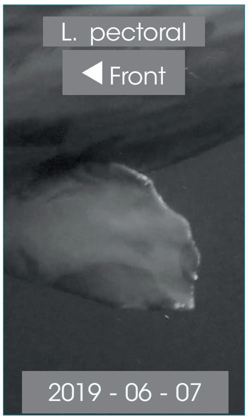

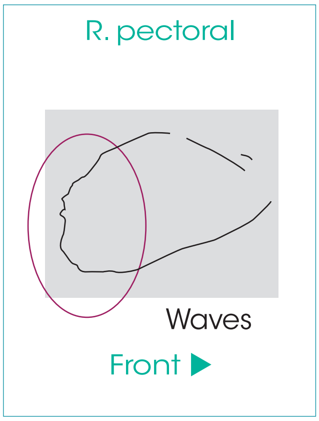

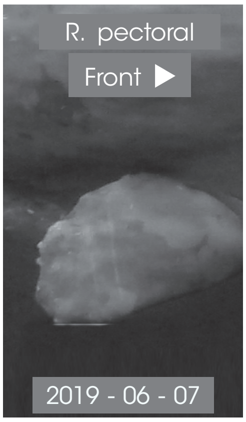

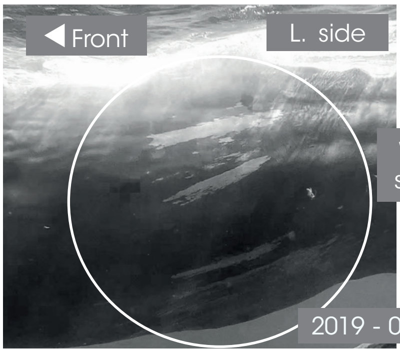

White stripes

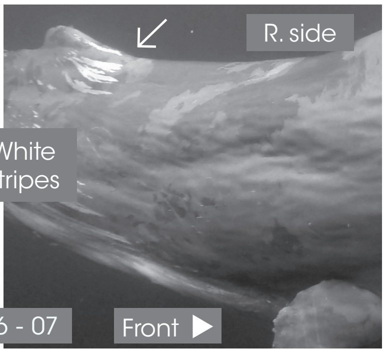

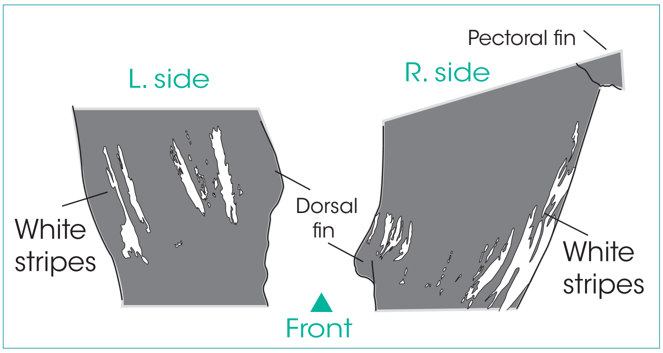

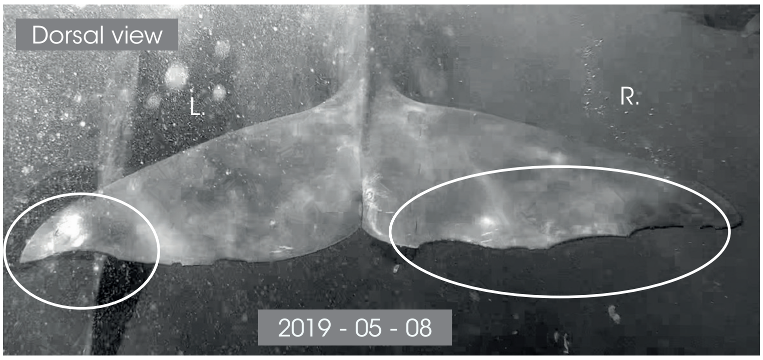

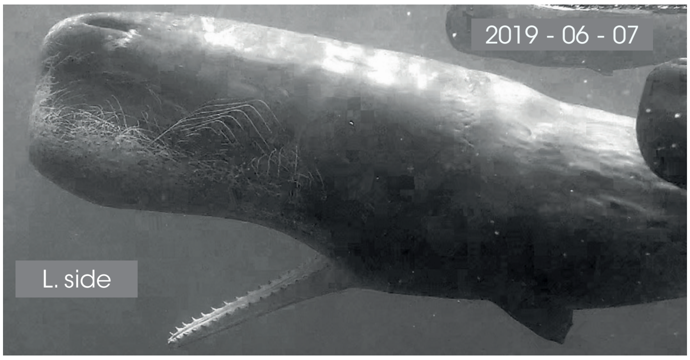

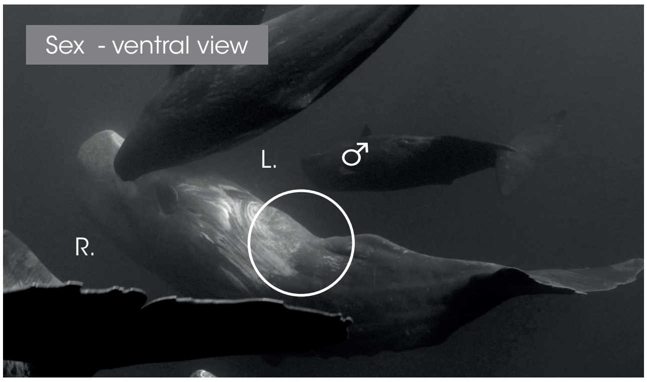

L. side

Front

R. side

Front

© Longitude 181 - Creation 2015 / Update 2020 - Concept : F. Sarano - Graphic design & illustrations : M. Sarano - © Pictures : S. Granzotto, F. Guérin, R. Heuzey, A. Preud'homme, V. & F. Sarano.

### NOÉ

*1ST OBS. : 2018 - 04 -16 // LAST OBS. : 2018 - 06 -25*

| *ADULT* | *SEEN* | |
|---|---|---|
| *DNA* | 2018 | *ADDITIONAL INFORMATION* |
| ♂ | | • Curled left caudal tip.<br>• Ker2007's grandson<br>• Lana's father |

L. pectoral

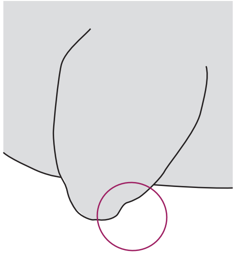

◀ Front

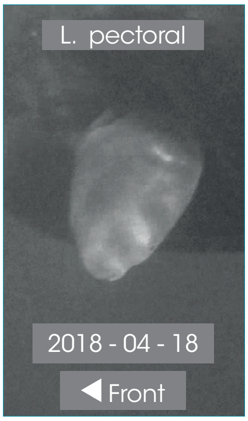

Dorsal view // 2018 - 04 - 16

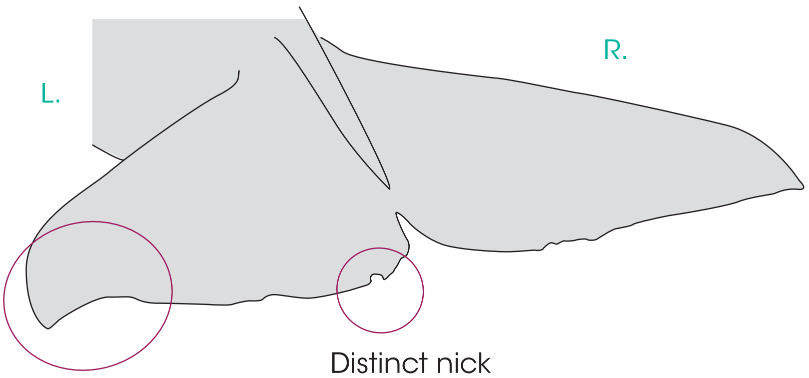

R. pectoral

Noé

Front ▶

R. pectoral

Front ▶

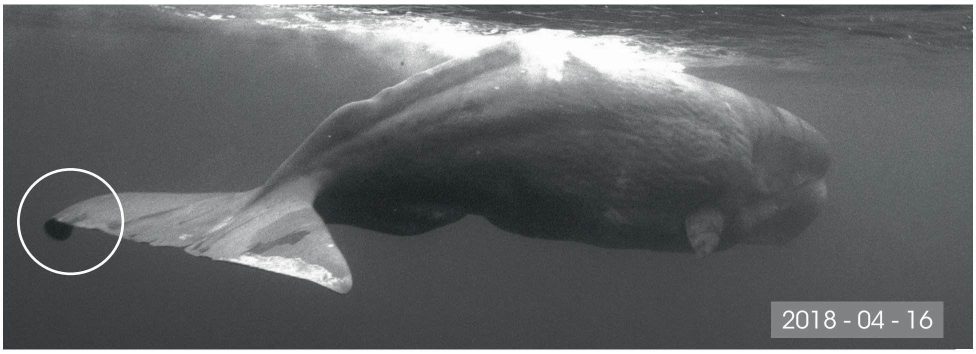

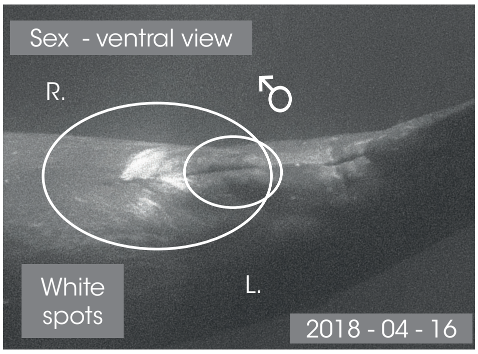

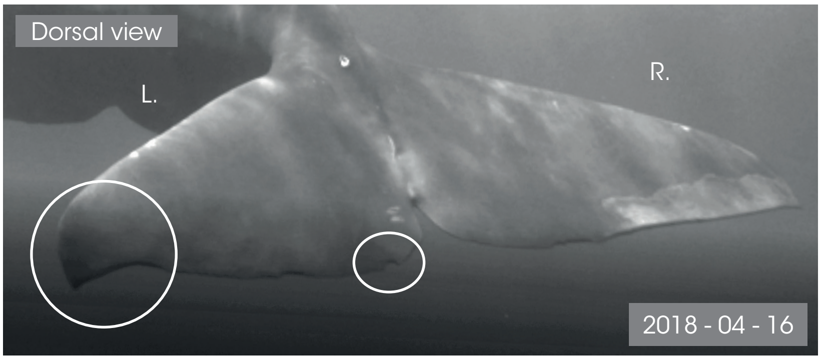

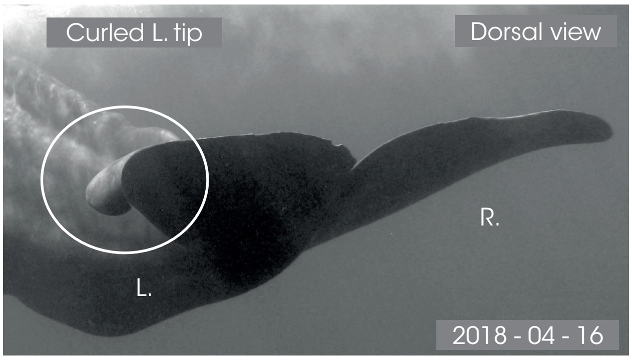

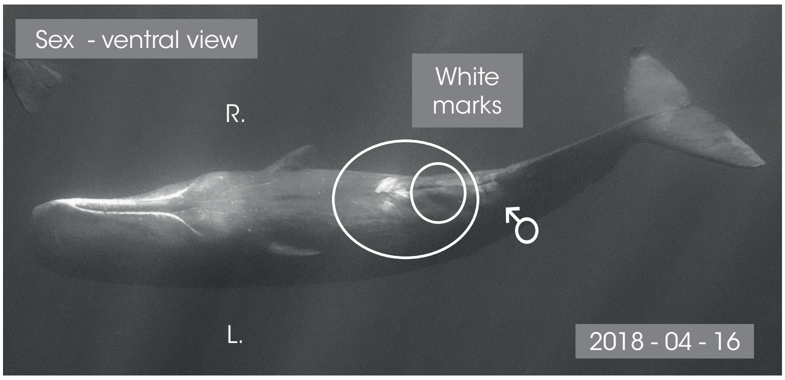

L. side

◀ Front

R. side

Front ▶

© Longitude 181 - Conception : François Sarano - Graphisme & illustration : Marion Sarano - © Photos : Stéphane Granzotto, Fabrice Guérin, René Heuzey, , Axel Preud'homme, Véronique & François Sarano, Hugues Vitry.

### REZA

**1ST OBS. : 2019 - 03 - 14 // LAST OBS. : 2019 - 03 - 28**

| ADULT | SEEN | |
|---|---|---|
| DNA | 2019 | ADDITIONAL INFORMATION |
| ♂ | | • White marks on caudal fin |

Dorsal view // 2019 - 03 - 19

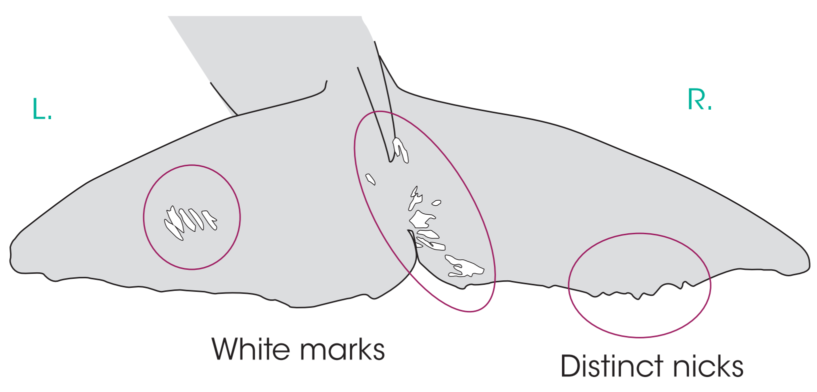

L. pectoral

**UNHARMED**

◀ Front

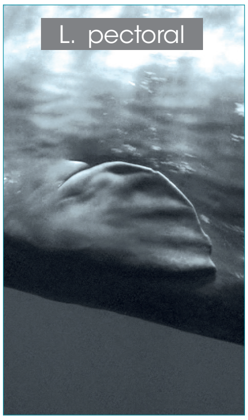

R. pectoral

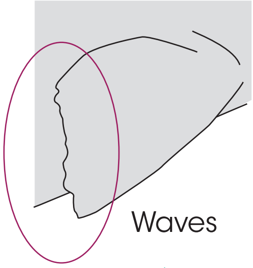

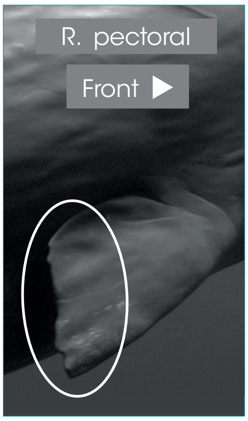

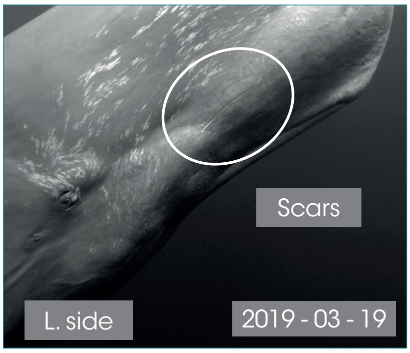

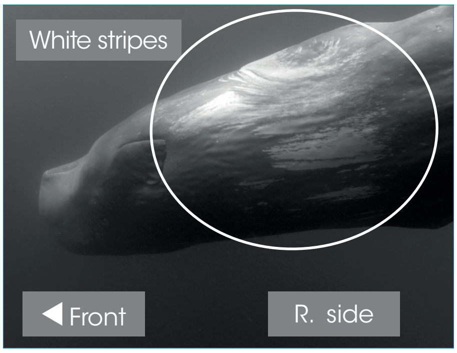

R. side

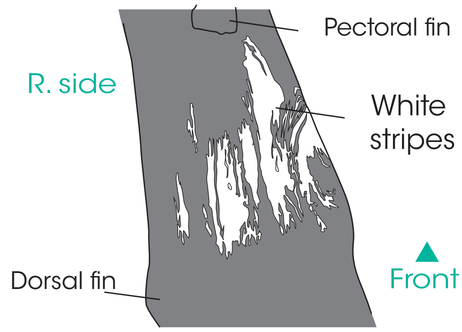

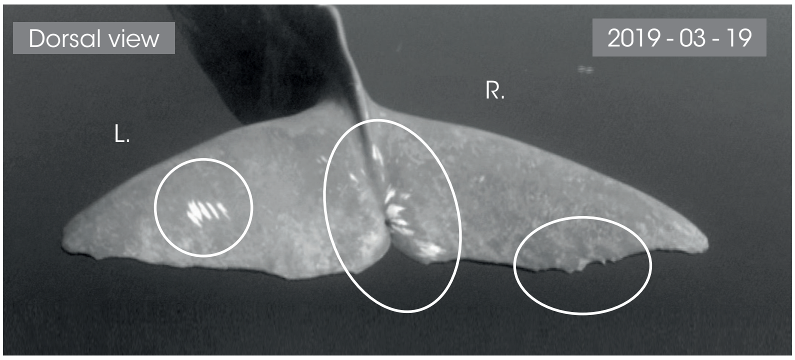

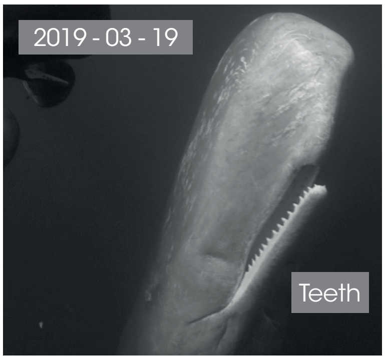

Sex - ventral view

L. side

◀ Front

R. side

Front ▶

© Longitude 181 - Creation 2015 / Update 2020 - Concept : F. Sarano - Graphic design & illustrations : M. Sarano - © Pictures : S. Granzotto, F. Guérin, R. Heuzey, A. Preud'homme, V. & F. Sarano.

### VASILILY

1ST OBS. : 2018 - 07 - 02 // LAST OBS. : 2018 - 07 - 02

| ADULT | SEEN | |
|---|---|---|
| DNA | 2018 | ADDITIONAL INFORMATION |
| ♂ | | • Missing portion on both caudal tips<br>• Large white escutcheon |

Dorsal view // 2018 - 07 - 02

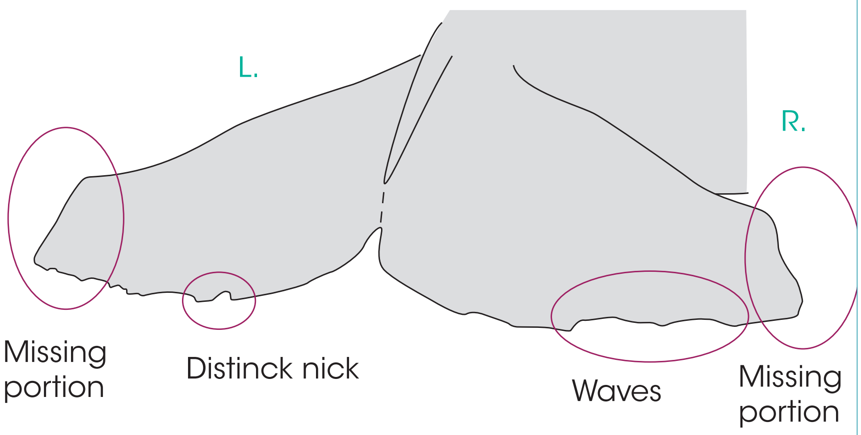

L. pectoral

UNHARMED

◀ Front

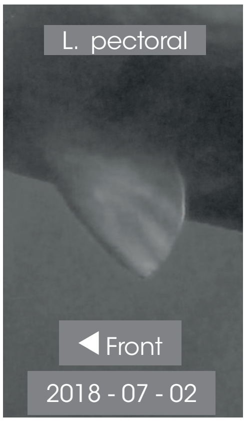

R. pectoral

UNHARMED

Front ▶

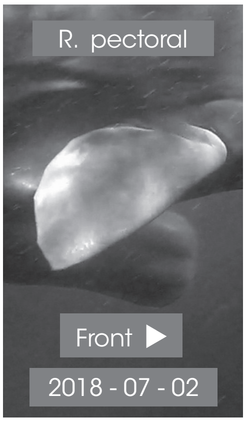

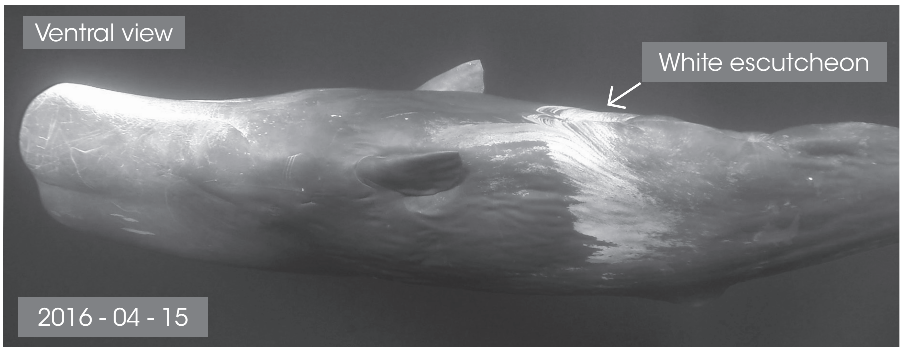

R. side

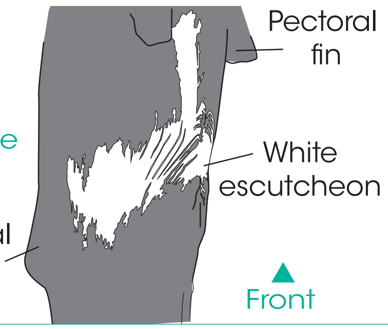

▲ Front

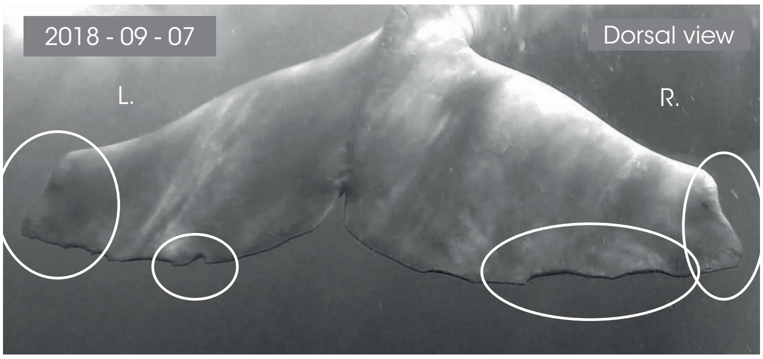

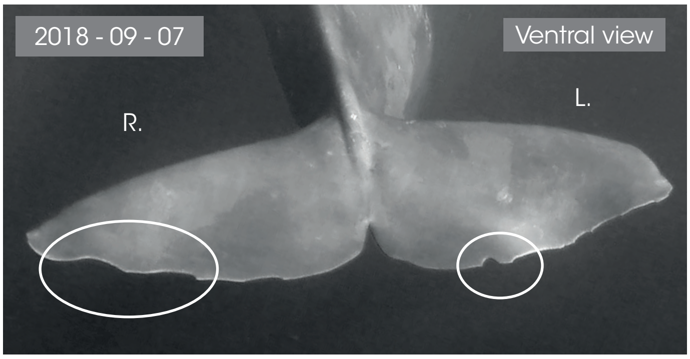

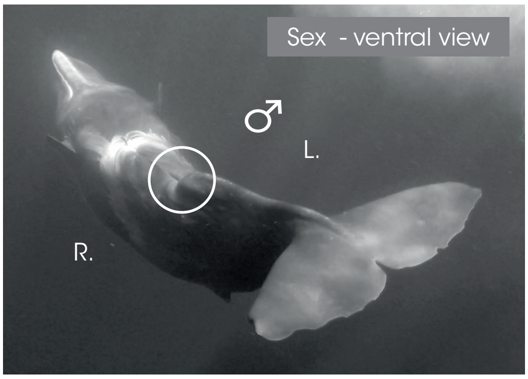

L. side

◀ Front

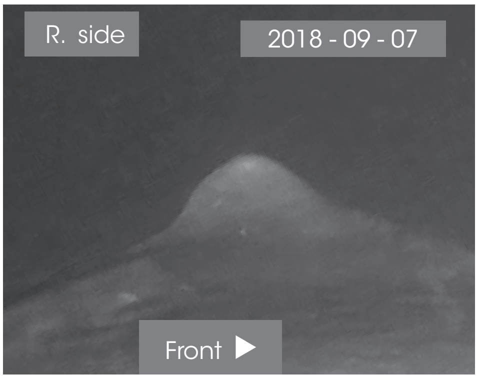

© Longitude 181 - Creation 2015 / Update 2020 - Concept : F. Sarano - Graphic design & illustrations : M. Sarano - © Pictures : S. Granzotto, F. Guérin, R. Heuzey, A. Preud'homme, V. & F. Sarano.
