## Supplementary information 2 for "Underwater photo-identification of marine megafauna: an identity card catalogue of sperm whales (*Physeter macrocephalus*) off Mauritius Island"

### FEMALES ♀

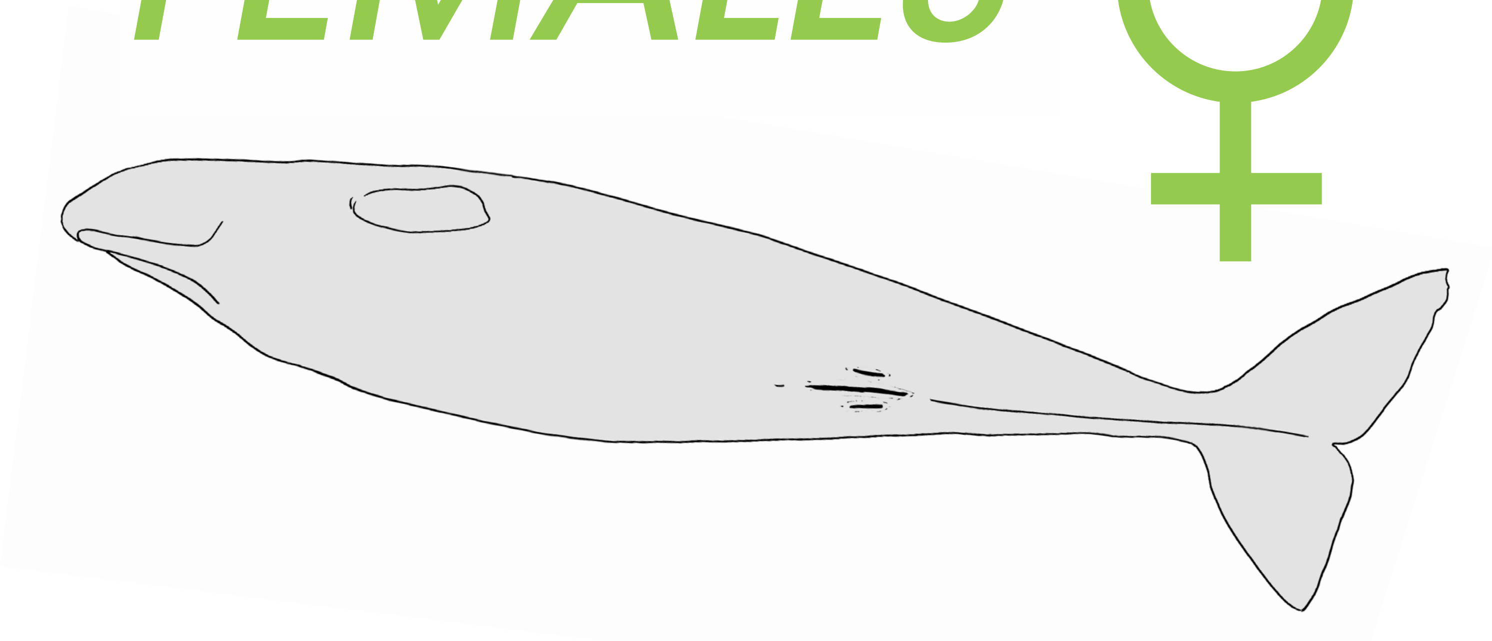

CONCEPT : François Sarano

GRAPHIC DESIGN & ILLUSTRATIONS : Marion Sarano

PICTURES : Stéphane Granzotto, Fabrice Guérin, René Heuzey, Daniel Jouannet, Bernard Kirchhofer, Vanessa Mignon, Axel Preud'homme, Véronique Sarano, François Sarano, Hugues Vitry.

© Longitude 181 - Creation 2015 / Update 2020

### ADÉLIE

**1ST OBS. : 2011 - 05 - 20 // LAST OBS. : 2020 - 03 - 19**

| ADULT | SEEN | |
|---|---|---|
| DNA | 2011 - 2012<br>2013 - 2014<br>2015 - 2016<br>2017 - 2018<br>2019 - 2020 | ADDITIONAL INFORMATION |
| ♀ | | • White escutcheon between pectoral fins.<br>• Convex caudal.<br>• Eliot's mother. |

Dorsal view // 2018 - 04 - 01

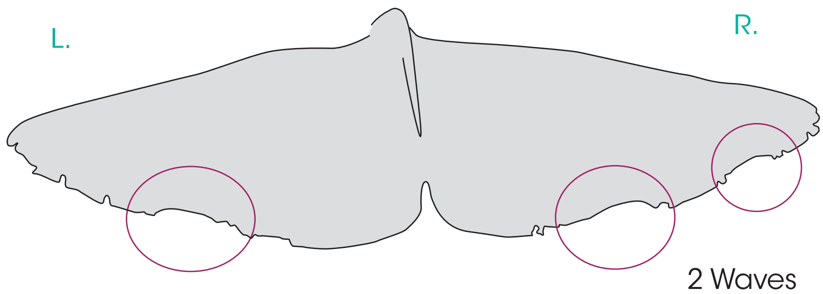

L. pectoral

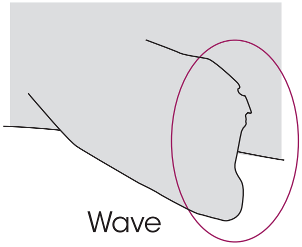

◀ Front

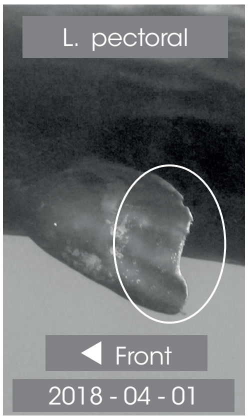

R. pectoral

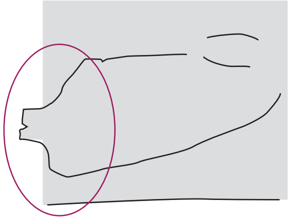

Front ▶

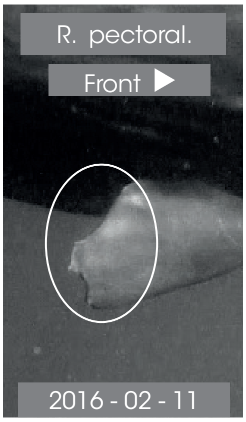

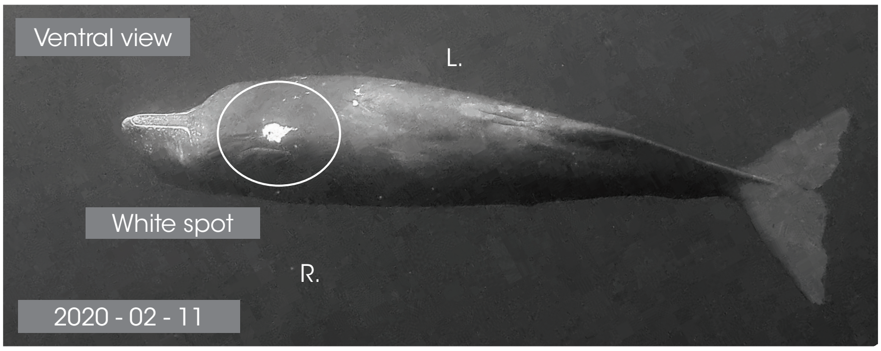

Ventral view

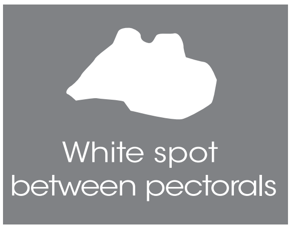

Front ▶

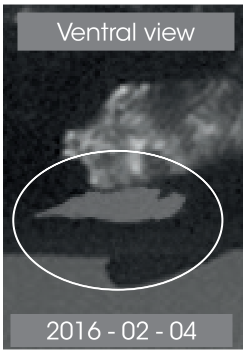

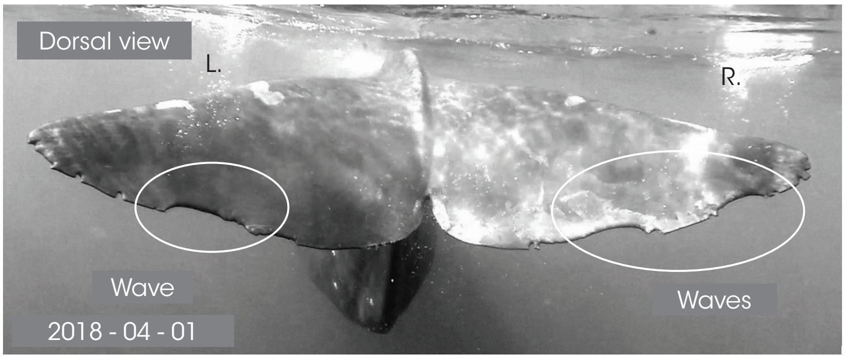

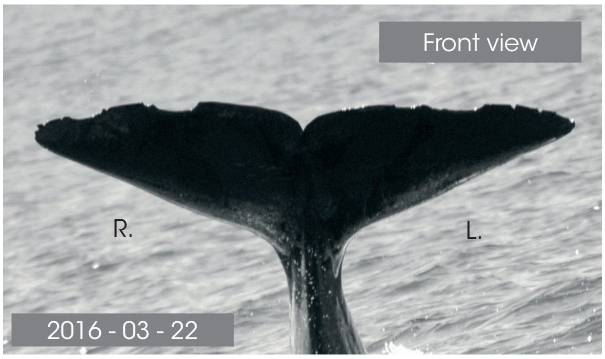

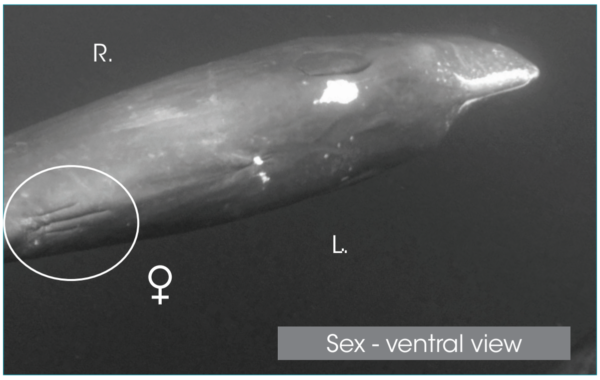

Dorsal - L. side

◀ Front

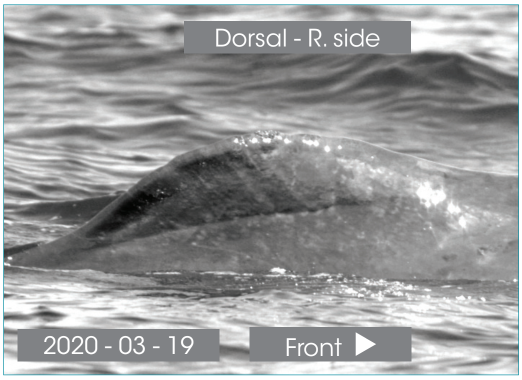

© Longitude 181 - Creation 2015 / Update 2020 - Concept : F. Sarano - Graphic design & illustrations : M. Sarano - © Pictures : S. Granzotto, F. Guérin, R. Heuzey, A. Preud'homme, V. & F. Sarano.

### AÏKO

**1ST OBS. : 2008 - 09 - 25 // LAST OBS. : 2020 - 03 - 12**

| ADULT | SEEN | |
|---|---|---|
| DNA | 2011 - 2012<br>2013 - 2014<br>2015 - 2016<br>2017 - 2018<br>2019 - 2020 | ADDITIONAL INFORMATION |
| ♀ | | • Large female.<br>• Only 1 distinct nick on R. pectoral in 2011. |

Dorsal view // 2018 - 03 - 22

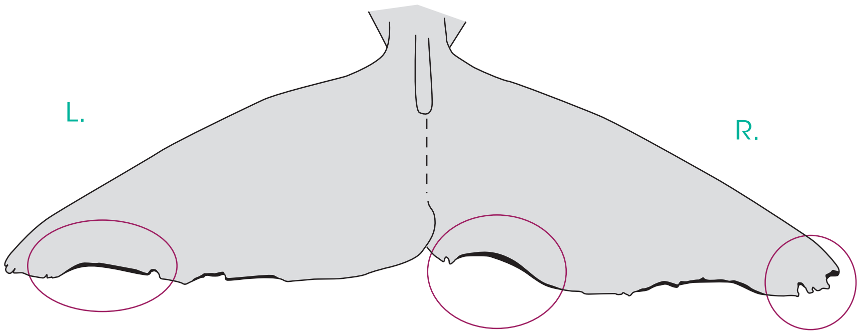

L. pectoral

UNHARMED

◀ Front

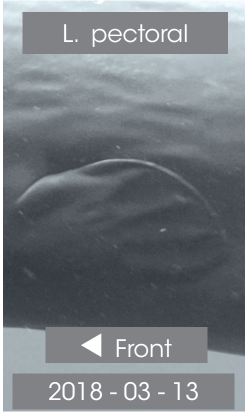

R. pectoral

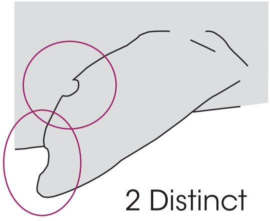

2 Distinct nicks

Front ▶

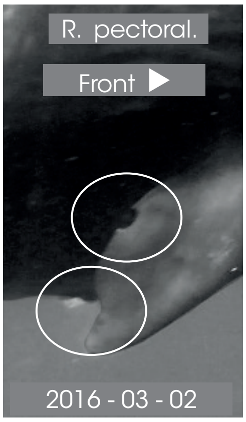

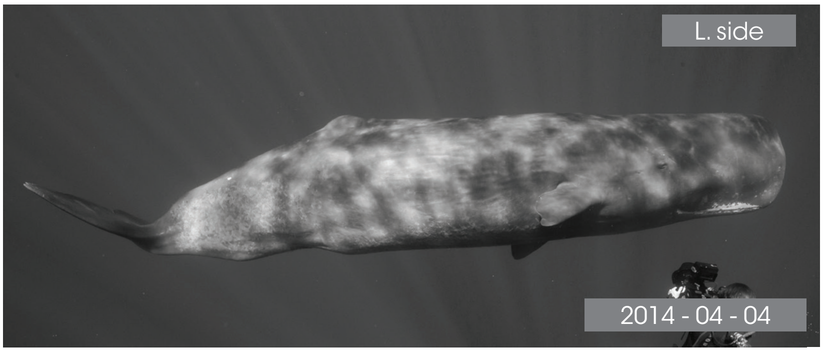

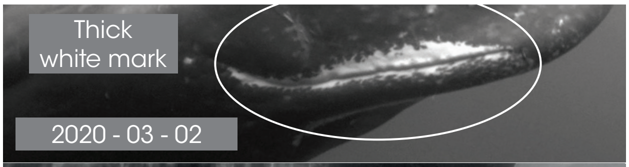

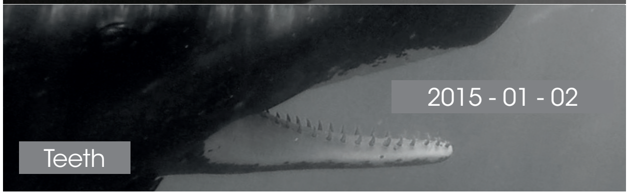

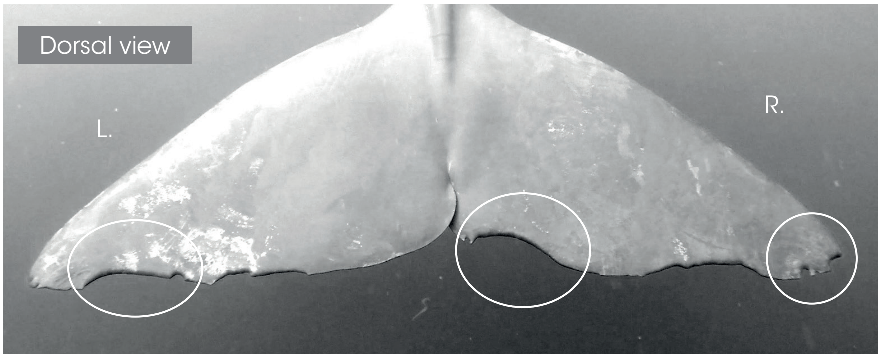

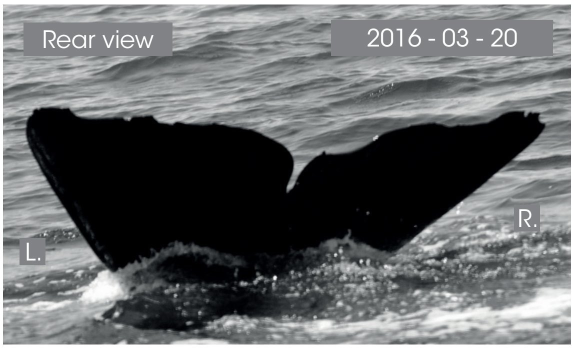

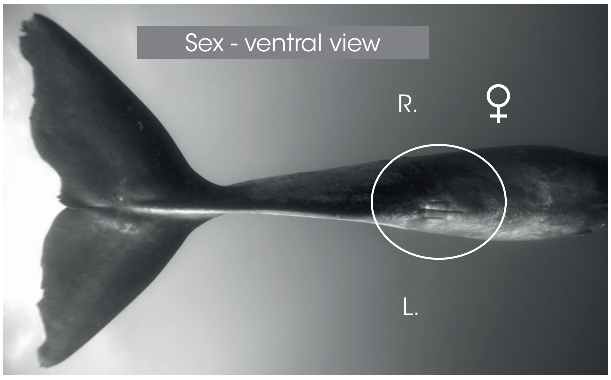

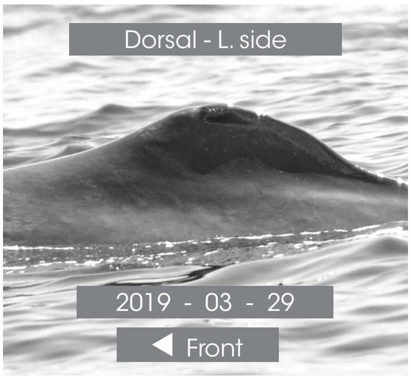

Dorsal - R. side

Front ▶

© Longitude 181 - Creation 2015 / Update 2020 - Concept : F. Sarano - Graphic design & illustrations : M. Sarano - © Pictures : S. Granzotto, F. Guérin, R. Heuzey, A. Preud'homme, V. & F. Sarano.

### CAROLINE

1ST OBS. : 2012 - 10 - 06 // LAST OBS. : 2020 - 02 - 25

| ADULT | SEEN | |
|---|---|---|
| DNA | - | ADDITIONAL INFORMATION |
| ♀ | 2012 2013<br>- ?<br>2015 - 2016<br>2017 - 2018<br>2019 - 2020 | • Zoé's mother & Alexander's mother. |

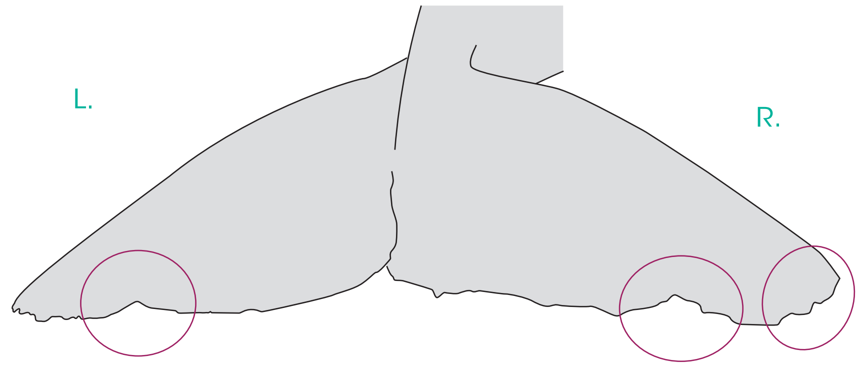

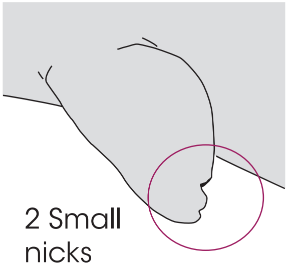

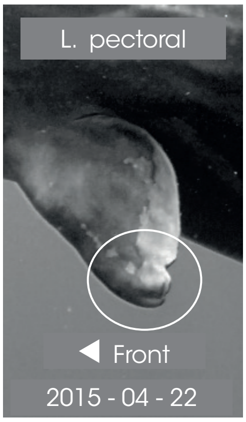

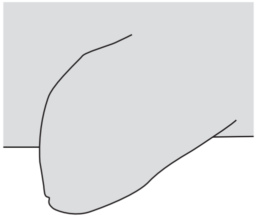

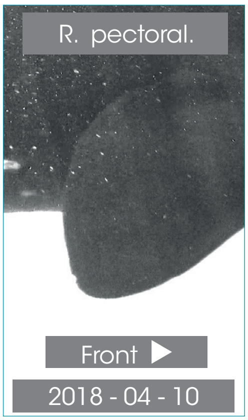

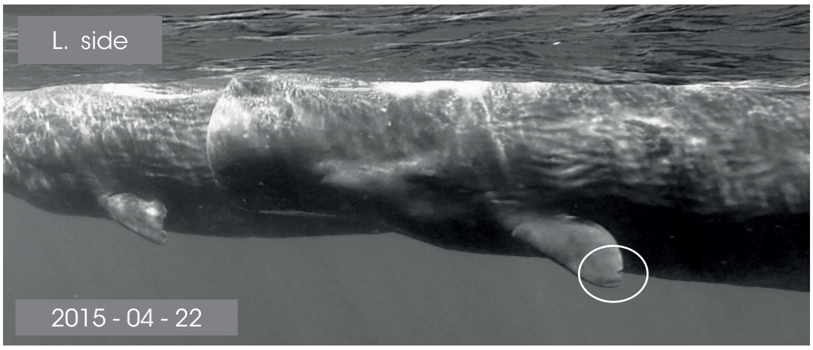

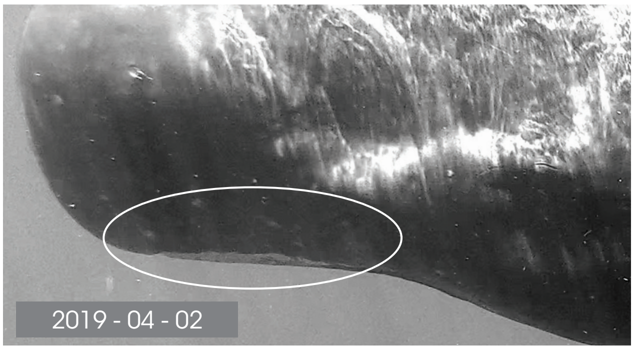

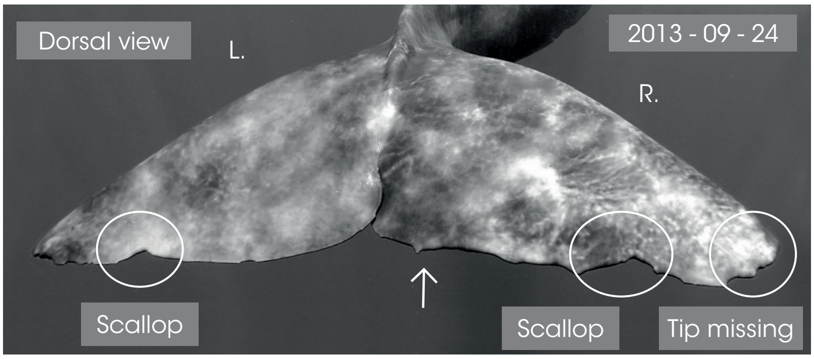

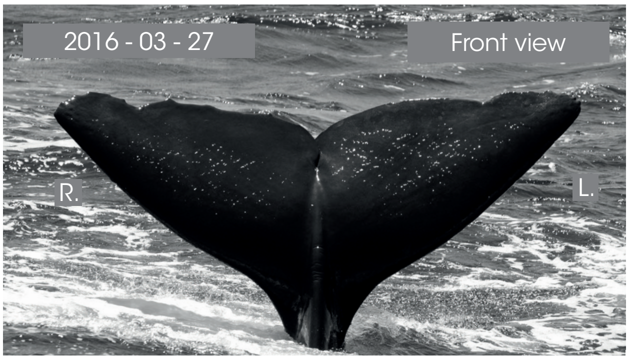

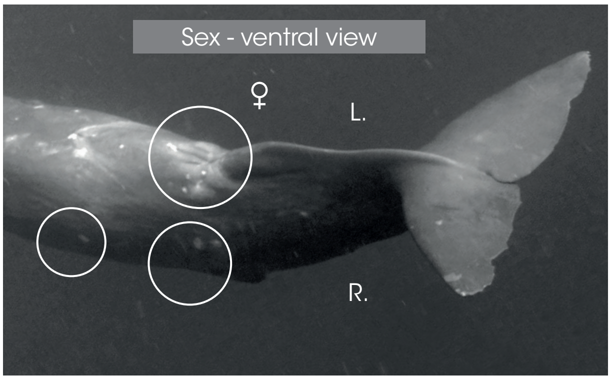

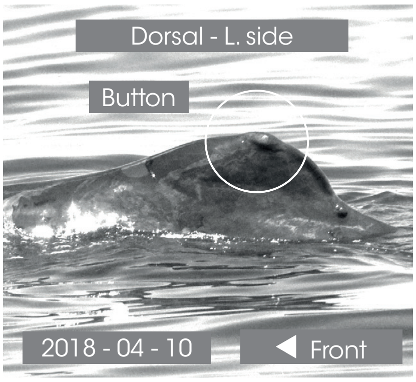

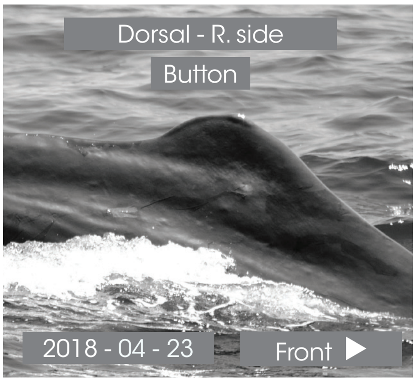

© Longitude 181 - Creation 2015 / Update 2020 - Concept : F. Sarano - Graphic design & illustrations : M. Sarano - © Pictures : S. Granzotto, F. Guérin, R. Heuzey, A. Preud'homme, V. & F. Sarano.

### CLAIRE

*1ST OBS. : 2011 - 05 - 17 // LAST OBS. : 2020 - 03 - 12*

| *ADULT* | *SEEN* | |
|---|---|---|
| *DNA* | 2011 - 2012<br>? - 2014<br>2015 - 2016<br>2017 - 2018<br>2019 - 2020 | *ADDITIONAL INFORMATION* |
| ♀ | | • Very pale color of the body.<br>• Haplotype different from the haplotype of the other adult females. |

Dorsal view // 2015 - 04 - 28

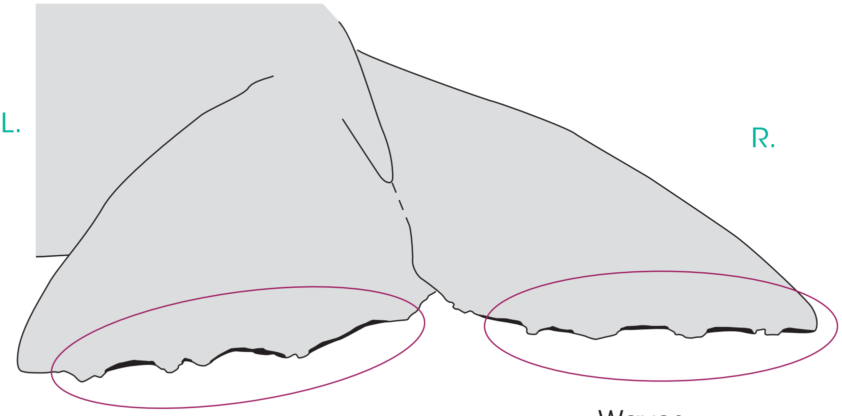

L. pectoral

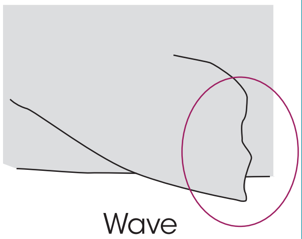

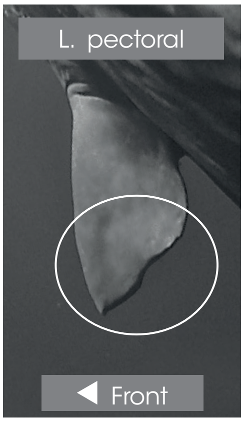

R. pectoral

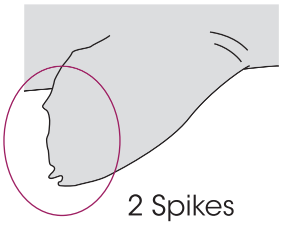

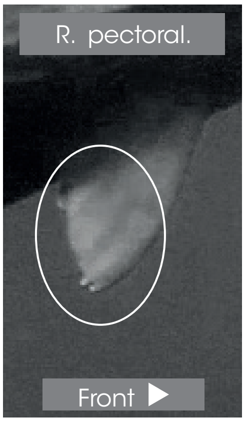

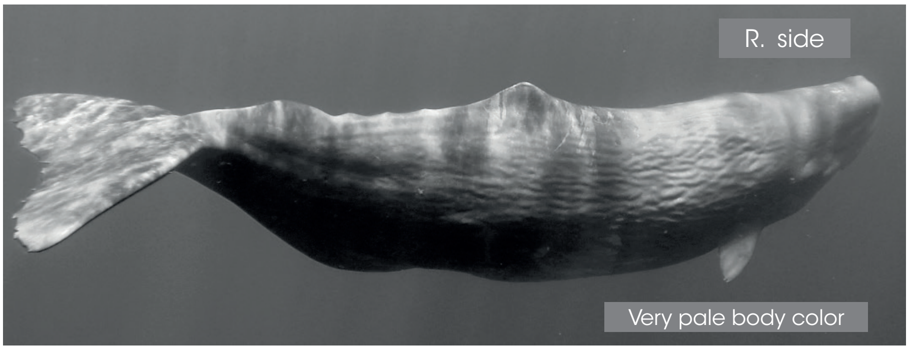

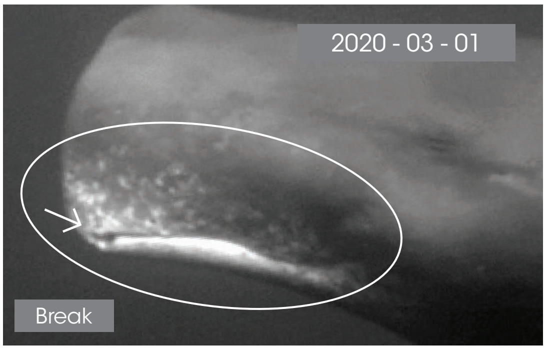

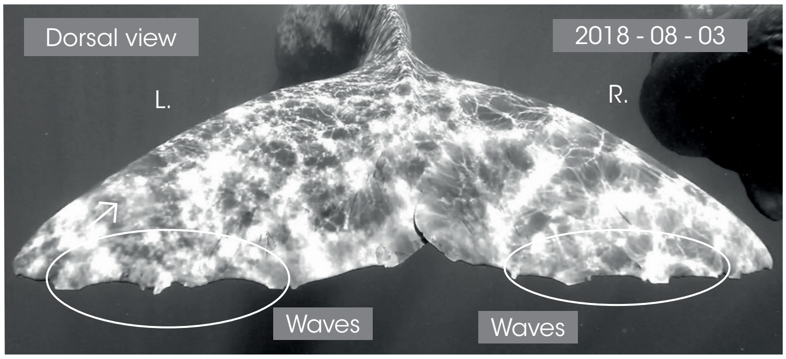

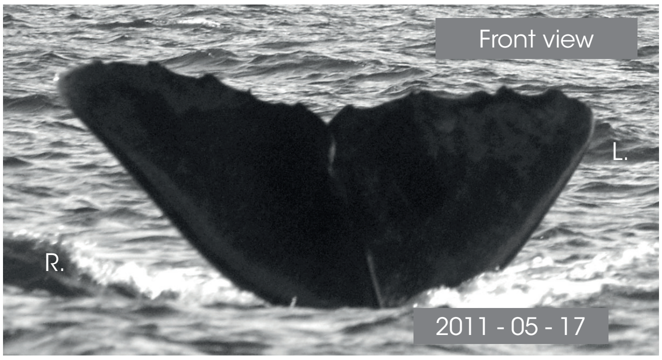

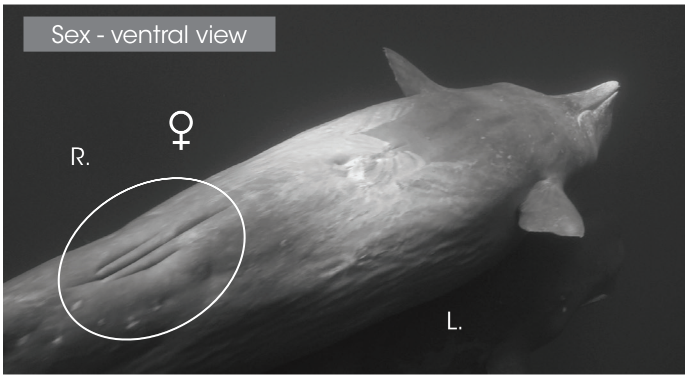

Dorsal - L. side

Front

2018 - 04 - 22

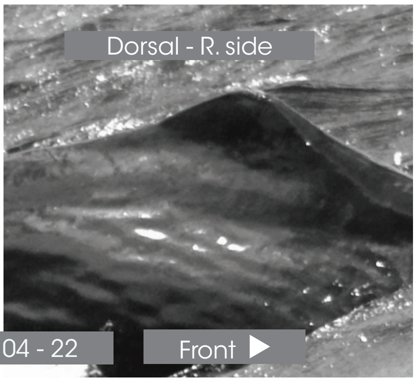

© Longitude 181 - Creation 2015 / Update 2020 - Concept : F. Sarano - Graphic design & illustrations : M. Sarano - © Pictures : S. Granzotto, F. Guérin, R. Heuzey, A. Preud'homme, V. & F. Sarano.

### DÉLINE

*1ST OBS. : 2009 - 02 - 25 // LAST OBS. : 2019 - 04 - 29*

| *ADULT* | *SEEN* | |
|---|---|---|
| - | 2011 - ?<br>? - ?<br>? - 2016<br>2017 - 2018<br>2019 | *ADDITIONAL INFORMATION* |
| ♀ | | • CAUTION: do not confuse Déline's dorsal with white callus with Vanessa's dorsal.<br>• CAUTION: caudal fin similar to Aïko's one but the pectorals are unharmed. |

L. pectoral

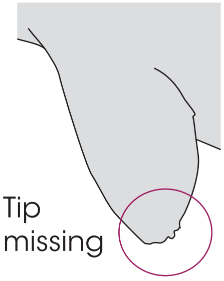

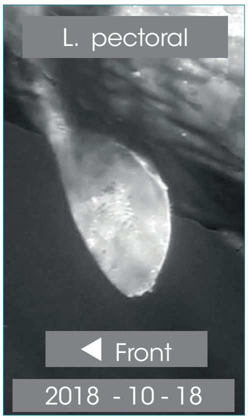

R. pectoral

**UNHARMED**

Front ▶

R. pectoral.

Front ▶

Dorsal view // 2016 - 05 - 06

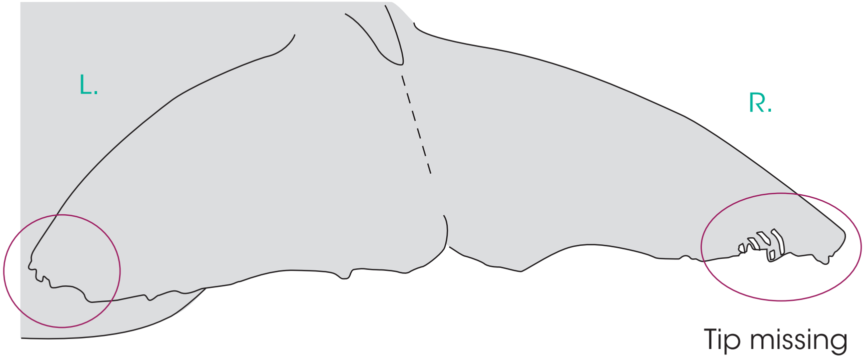

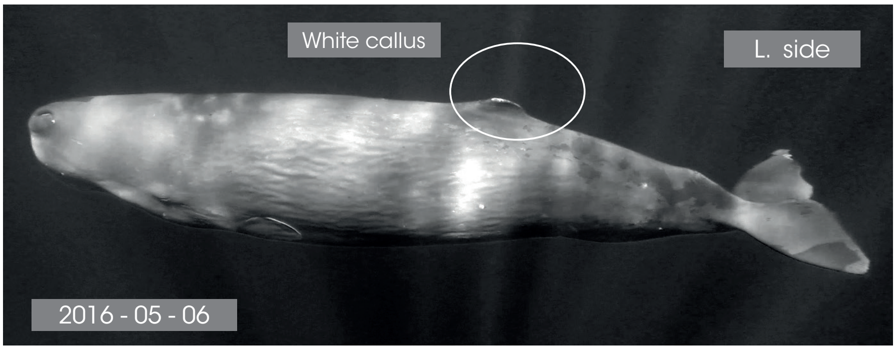

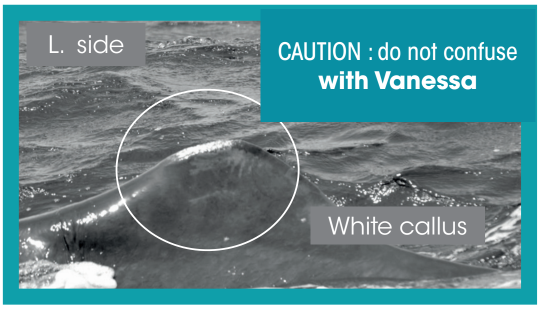

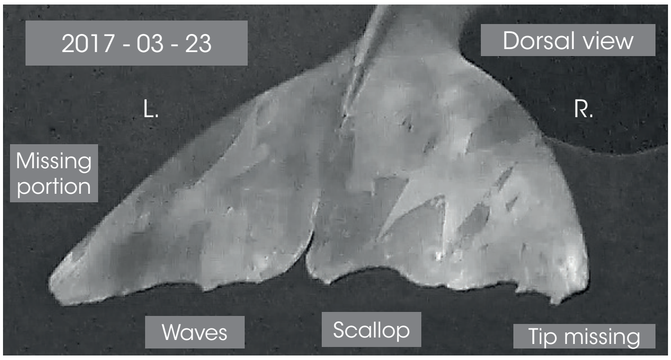

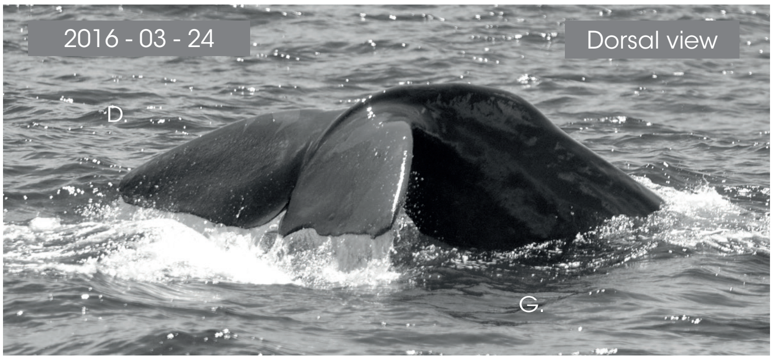

Sex - ventral view

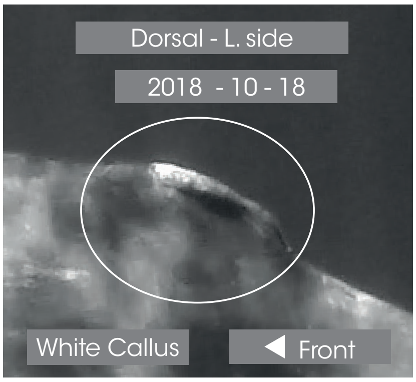

Dorsal - R. side

Front ▶

© Longitude 181 - Creation 2015 / Update 2020 - Concept : F. Sarano - Graphic design & illustrations : M. Sarano - © Pictures : S. Granzotto, F. Guérin, R. Heuzey, A. Preud'homme, V. & F. Sarano.

### DELPHINE

**1ST OBS. : 2011 - 05 - 16 // LAST OBS. : 2020 - 03 - 18**

| ADULT | SEEN | |
|---|---|---|
| DNA | 2011 - 2012<br>2013 - 2014<br>2015 - 2016<br>2017 - 2018<br>2019 - 2020 | ADDITIONAL INFORMATION |
| ♀ | | • Very dark mandibular zone, without white mark.<br>• 3 white stripes close to the navel.<br>• Pregnant with Tache Blanche : 20th May 2011<br>• Pregnant with Chesna : 14th Feb. 2018 |

Dessus // 2018 - 03 - 02

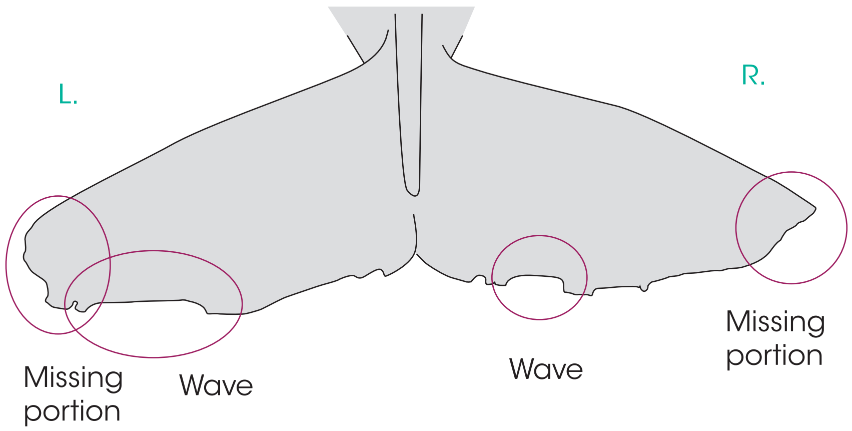

L. pectoral

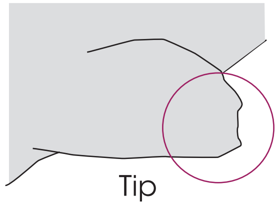

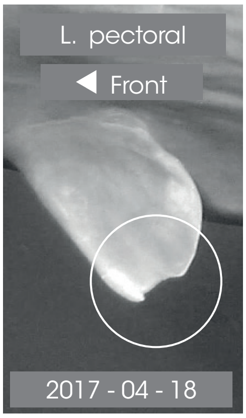

R. pectoral

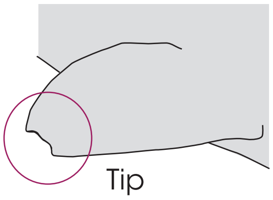

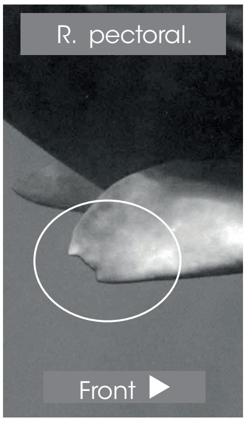

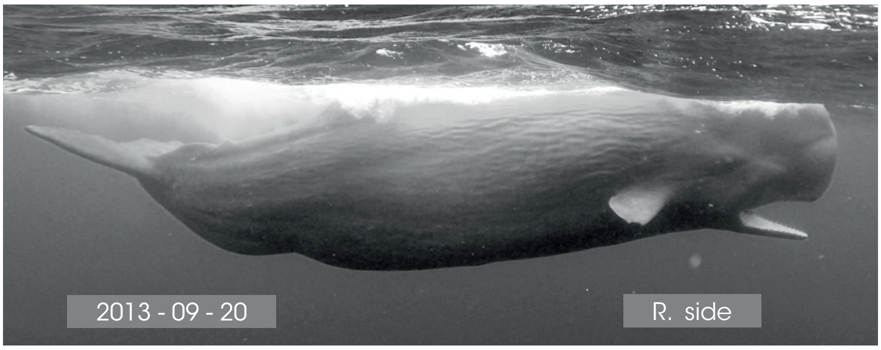

Ventral view

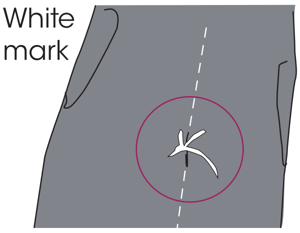

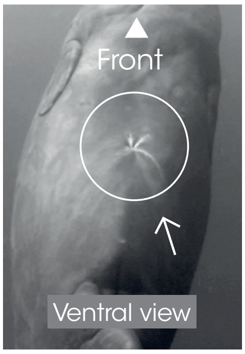

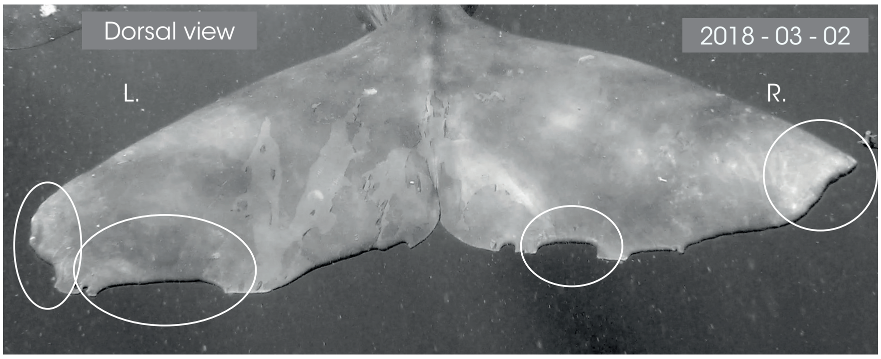

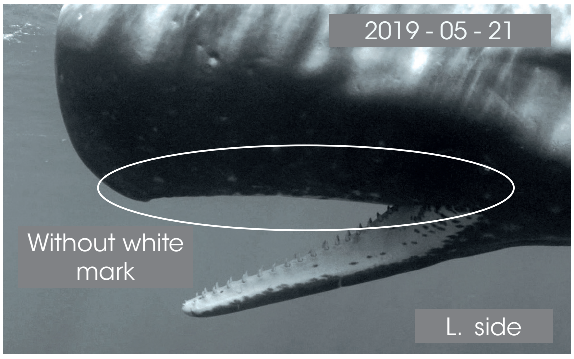

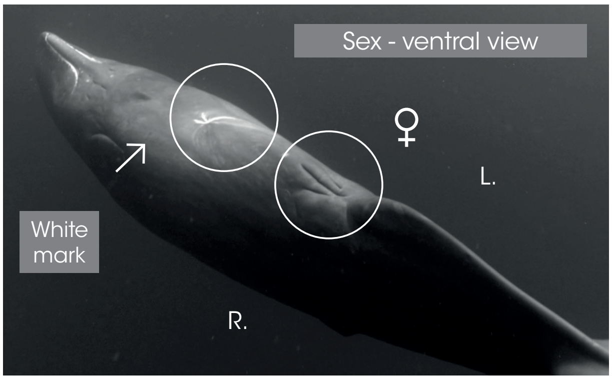

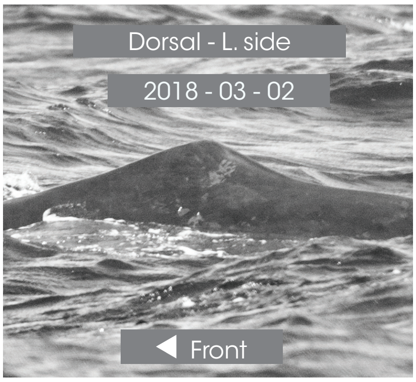

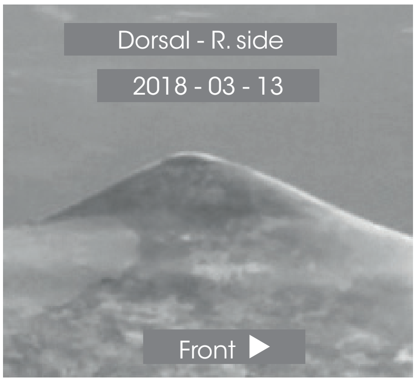

© Longitude 181 - Creation 2015 / Update 2020 - Concept : F. Sarano - Graphic design & illustrations : M. Sarano - © Pictures : S. Granzotto, F. Guérin, R. Heuzey, A. Preud'homme, V. & F. Sarano.

LONGITUDE 181
La Voix de l'Océan

IDENTITY CARDS
**SPERM WHALES OF MAURITIUS I.**

### DOS CALLEUX

*1ST OBS. : 2008 - 05 - 12 // LAST OBS. : 2020 - 03 - 12*

| ADULT | SEEN | |
|---|---|---|
| DNA | 2011 - 2012<br>? - ?<br>2015 - 2016<br>2017 - 2018<br>2019 - 2020 | ADDITIONAL INFORMATION |
| ♀ | | • Large callosity on the lower part of the dorsal.<br>• Dark mandibular zone, without white mark.<br>• Pregnant with Baptiste: 27th Feb 2017.<br>• Mina's mother & Lucy's mother<br>• Suckles Ali, her grandson, in 2018. |

L. pectoral

DOS CALLEUX

◀ Front

L. pectoral

◀ Front

Dorsal view // 2015 - 04 - 25

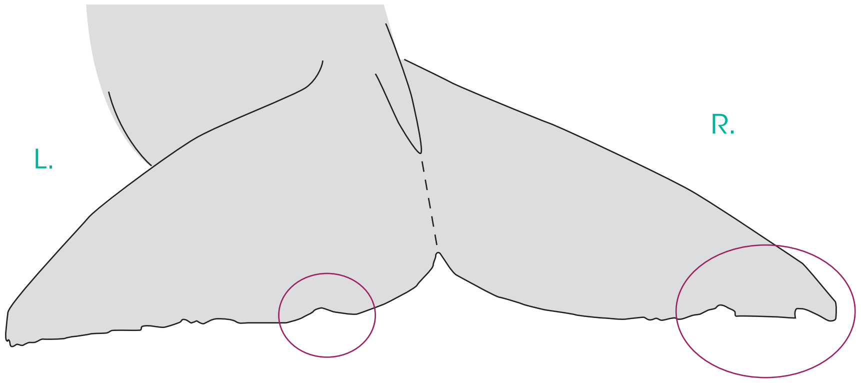

R. pectoral

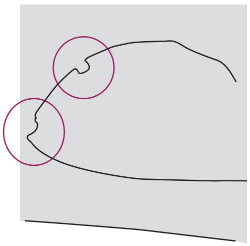

Front ▶

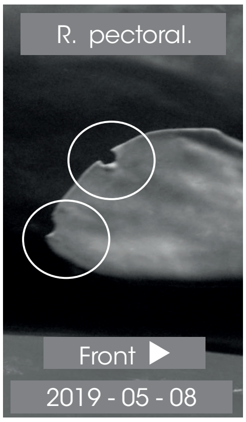

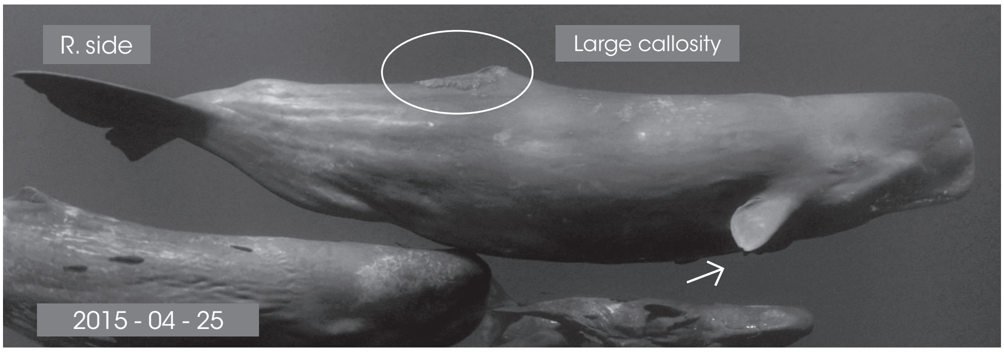

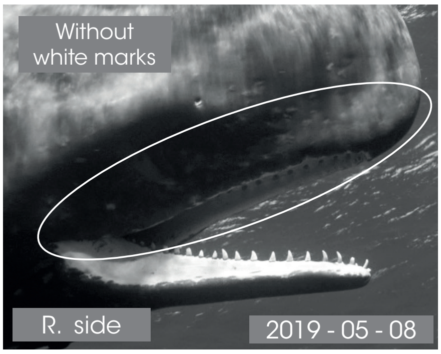

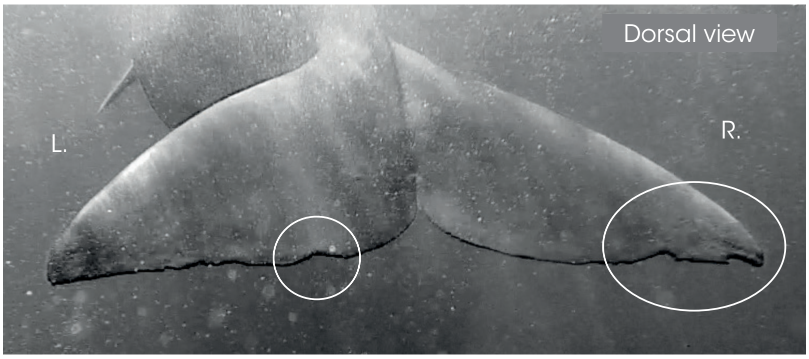

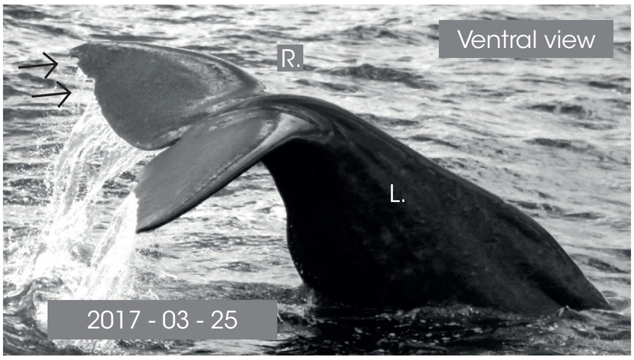

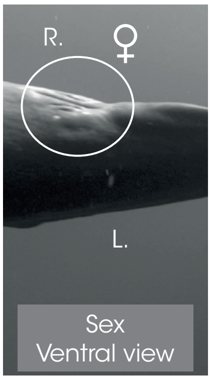

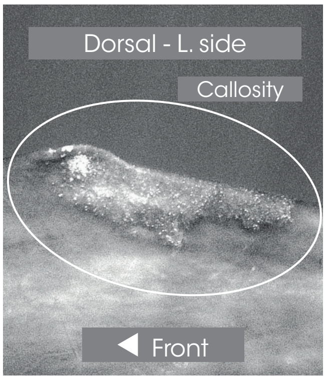

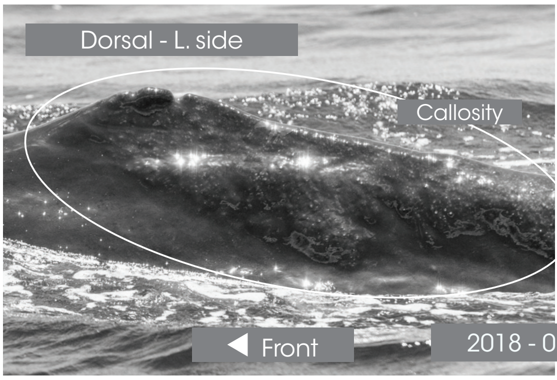

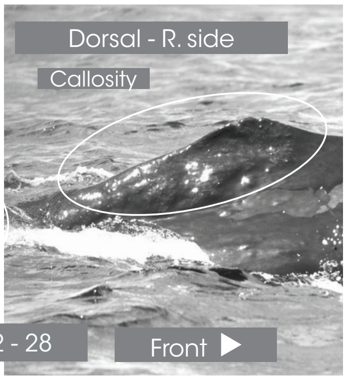

© Longitude 181 - Creation 2015 / Update 2020 - Concept : F. Sarano - Graphic design & illustrations : M. Sarano - © Pictures : S. Granzotto, F. Guérin, R. Heuzey, A. Preud'homme, V. & F. Sarano.

© Longitude 181 - Creation 2015 / Update 2020 - Concept : F. Sarano - Graphic design & illustrations : M. Sarano - © Pictures : S. Granzotto, F. Guérin, R. Heuzey, A. Preud'homme, V. & F. Sarano.

#### EMY

*1ST OBS. : 2007 - 06 - 24 // LAST OBS. : 2020 - 03 - 19*

| *ADULT* | *SEEN* | |
|---|---|---|
| *DNA* | 2011 - 2012<br>2013 - 2014<br>2015 - 2016<br>2017 - 2018<br>2019 - 2020 | *ADDITIONAL INFORMATION*<br>• L. caudal tip : missing portion.<br>• Scallop on the back side of the dorsal which was unharmed until March 2020.<br>• Chesna's nurse & Alexander's nurse |
| ♀ | | |

Dorsal view // 2017 - 04 - 18

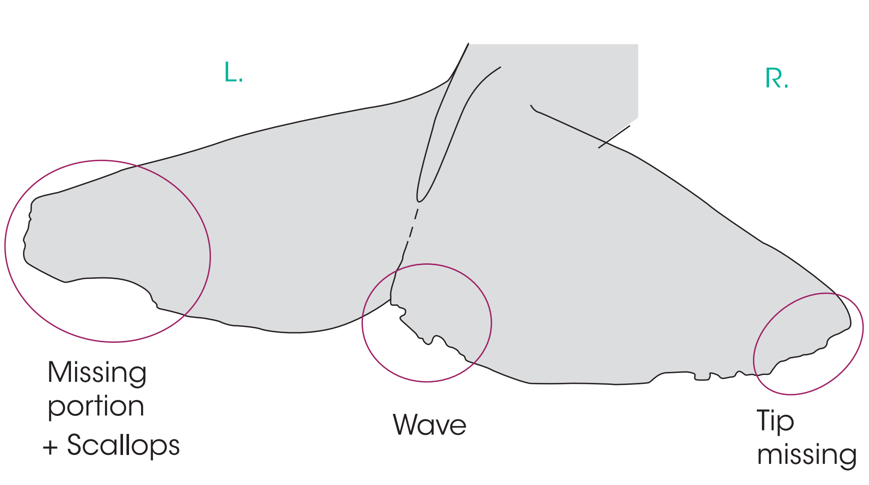

L. pectoral

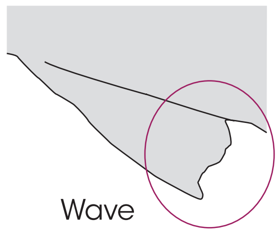

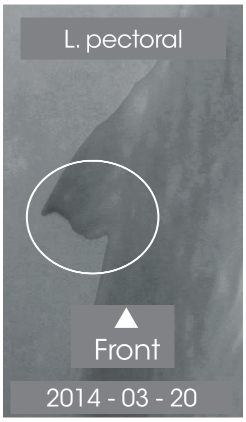

R. pectoral

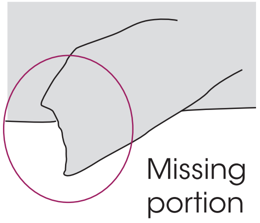

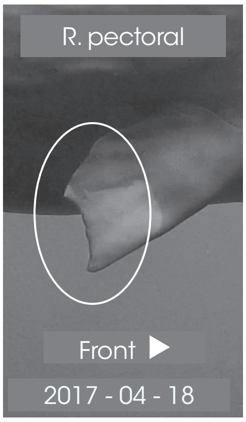

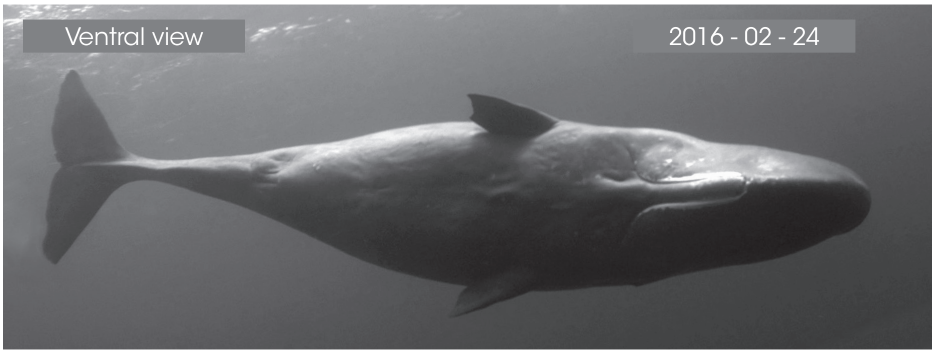

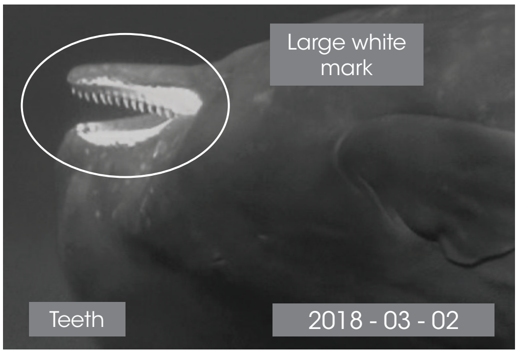

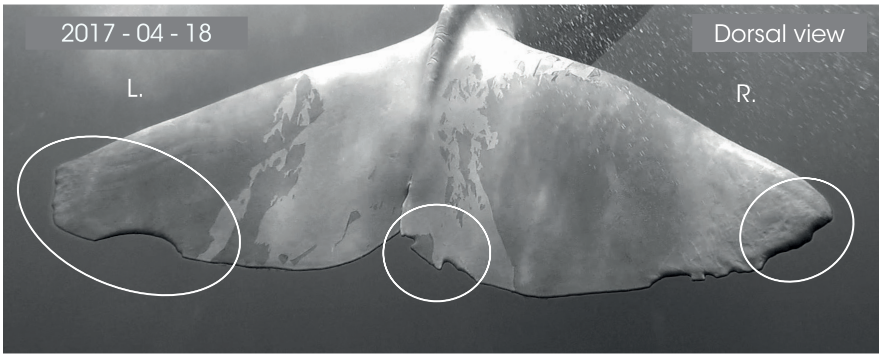

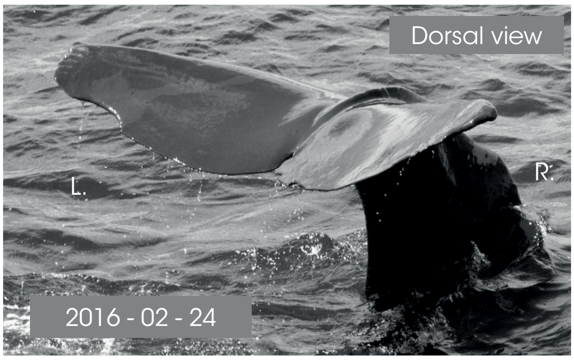

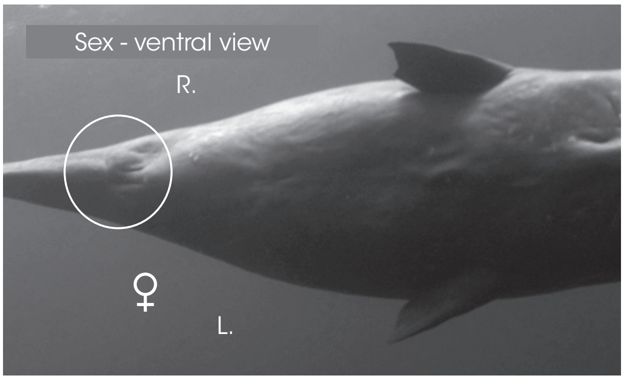

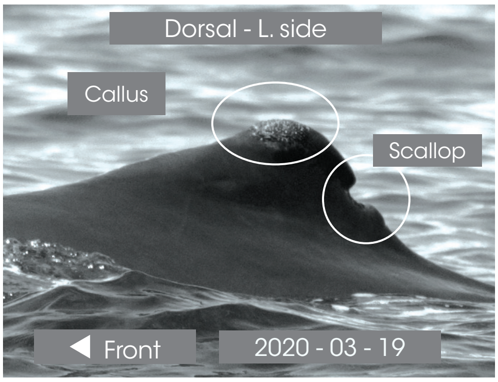

Dorsal - R. side

Front

### GERMINE

1ST OBS. : 2009 - 06 - 13 // LAST OBS. : 2020 - 03 - 12

| ADULT | SEEN | |
|---|---|---|
| DNA | 2011 - 2012<br>2013 - 2014<br>2015 - 2016<br>2017 - 2018<br>2019 - 2020 | ADDITIONAL INFORMATION |
| ♀ | | • Convex caudal.<br>• Ventral white escutcheon.<br>• Nurse of all juveniles, from Eliot (20th May 2011) until Lana (2020). |

Dorsal view // 2019 - 11 - 13

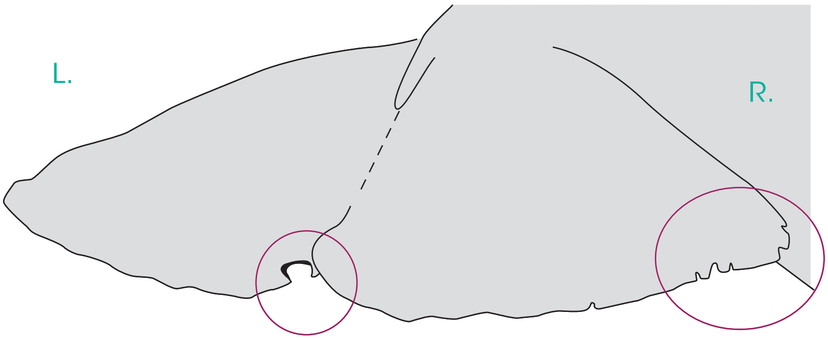

L. pectoral

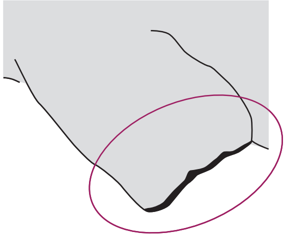

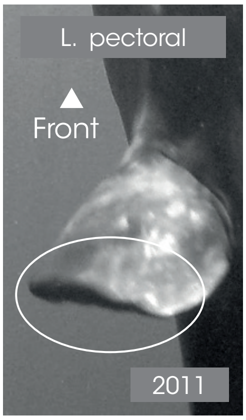

R. pectoral

**UNHARMED**

Front ▶

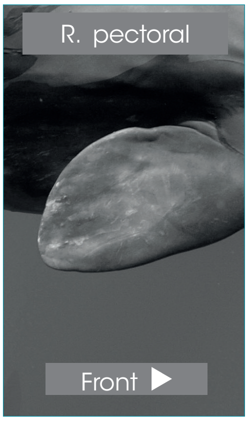

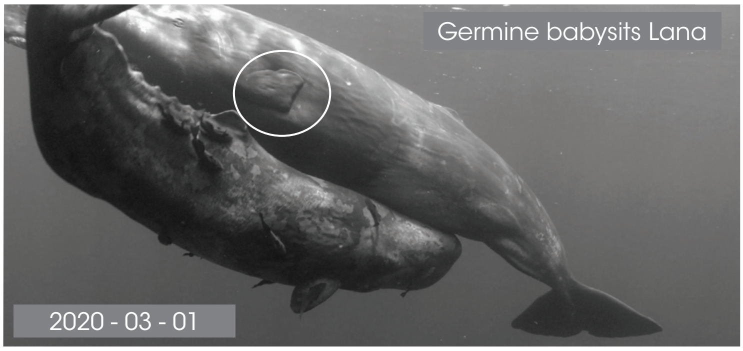

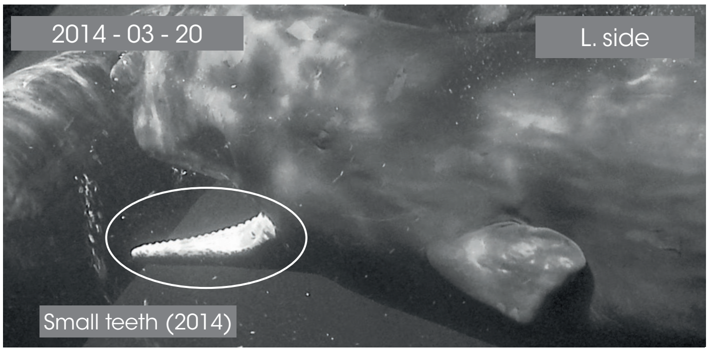

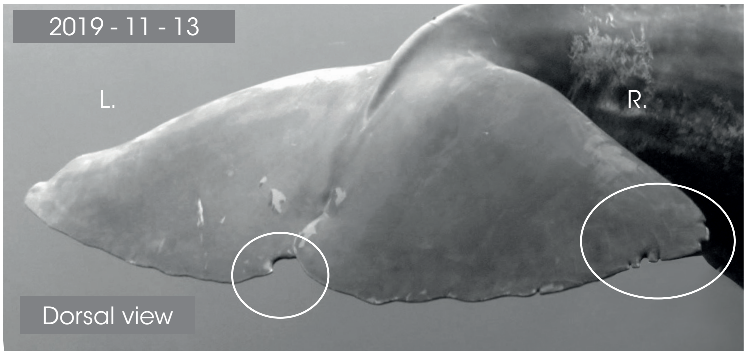

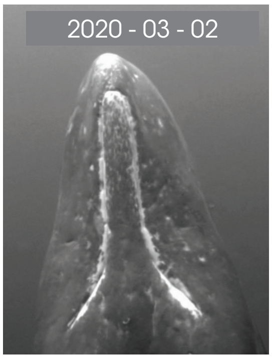

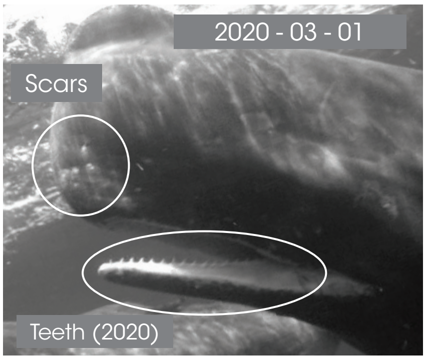

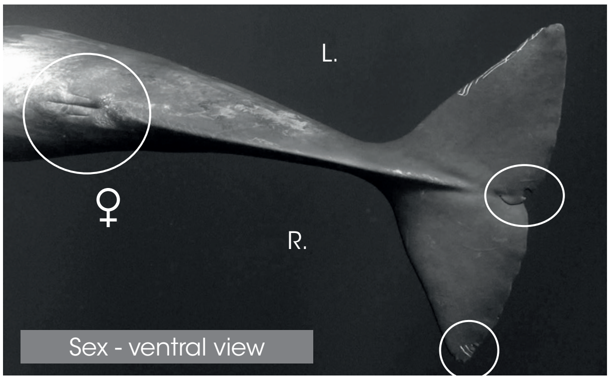

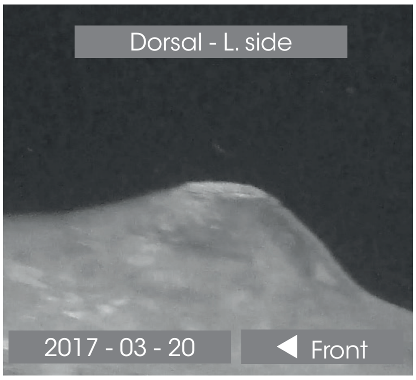

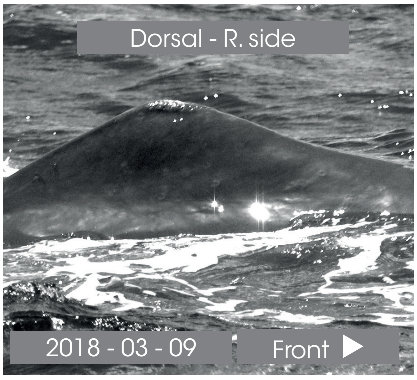

© Longitude 181 - Creation 2015 / Update 2020 - Concept : F. Sarano - Graphic design & illustrations : M. Sarano - © Pictures : S. Granzotto, F. Guérin, R. Heuzey, A. Preud'homme, V. & F. Sarano.

### IRÈNE GUEULE TORDUE

**1ST OBS. : 2009 - 01 - 18 // LAST OBS. : 2020 - 03 - 12**

| ADULT | SEEN | |
|---|---|---|
| DNA | 2011 - 2012<br>2013 - 2014<br>2015 - 2016<br>2017 - 2018<br>2019 - 2020 | ADDITIONAL INFORMATION |
| ♀ | | • Arched-shaped jaw.<br>• Arthur's mother. Parturition: probably on 5th April 2013.<br>• Ali's nurse.<br>• Lana's mother. Parturition: 20-21st Feb. 2019. |

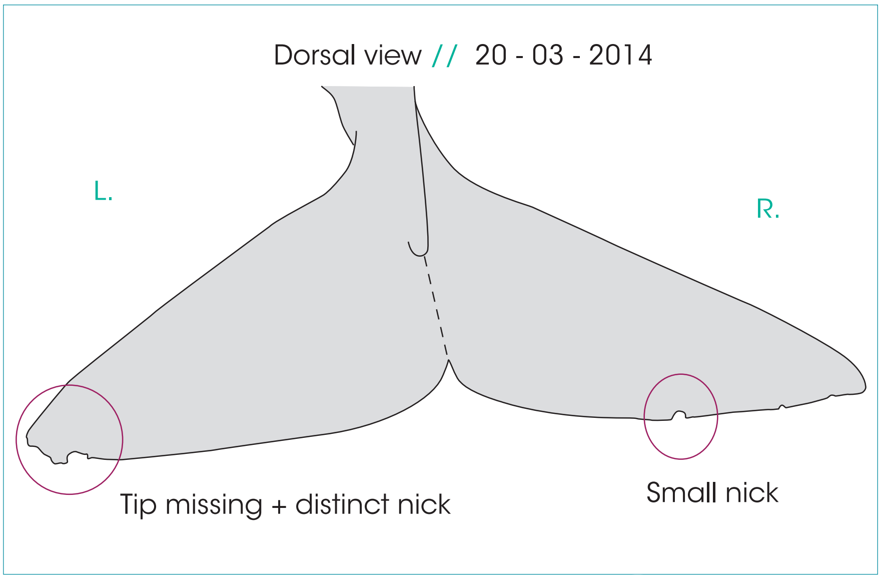

L. pectoral

INTACTE

◀ Front

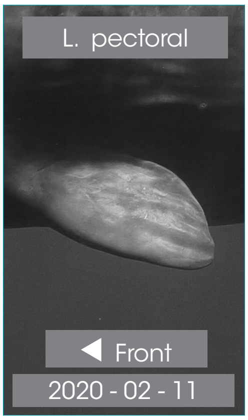

R. pectoral

INTACTE

Front ▶

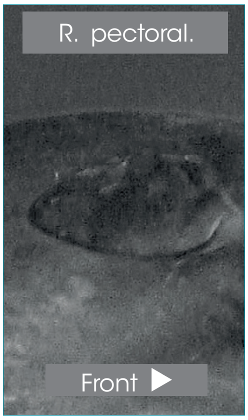

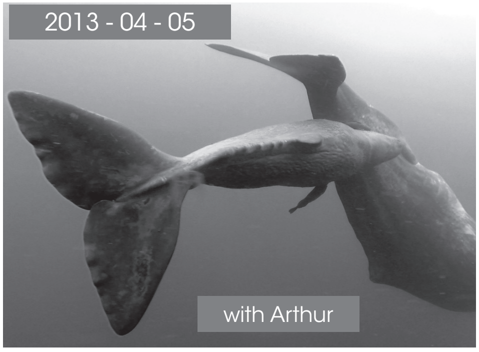

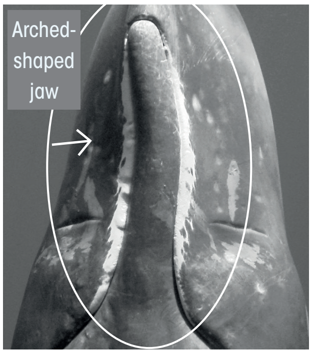

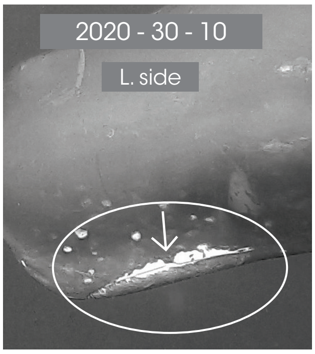

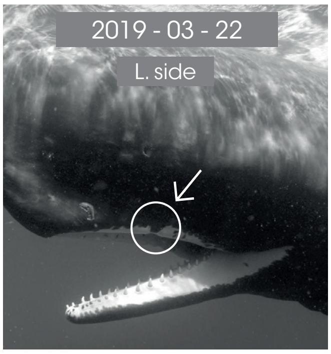

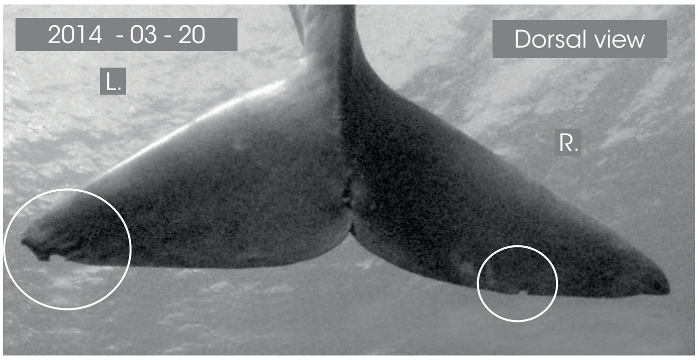

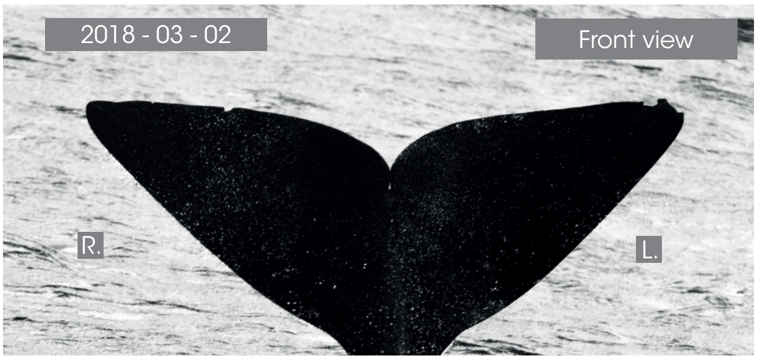

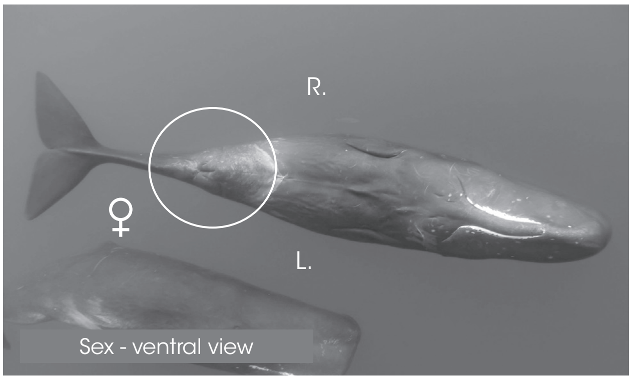

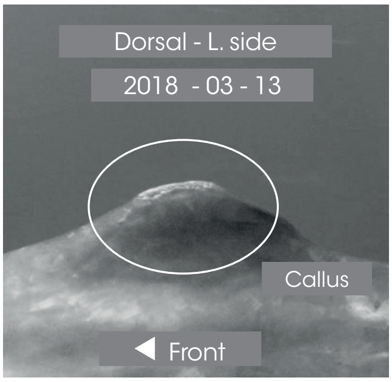

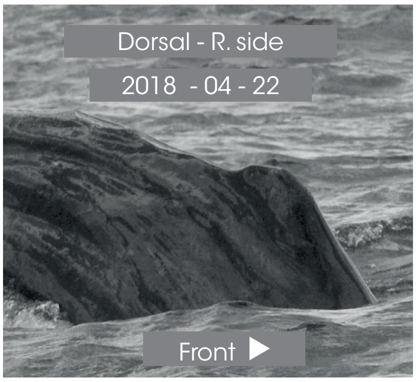

© Longitude 181 - Creation 2015 / Update 2020 - Concept : F. Sarano - Graphic design & illustrations : M. Sarano - © Pictures : S. Granzotto, F. Guérin, R. Heuzey, A. Preud'homme, V. & F. Sarano.

### ISSA

1<sup></sup>ST OBS. : 2009 - 02 - 25 // LAST OBS. : 2020 - 03 - 12

| ADULT | SEEN | |
|---|---|---|
| DNA | 2011 - ?<br>2013 - 2014<br>2015 - 2016<br>2017 - 2018<br>2019 - 2020 | ADDITIONAL INFORMATION |
| ♀ | | • White spot (butterfly-shaped) close to the navel.<br>• Wound between pectorals (17th April 2017).<br>• Convex caudal.<br>• Miss Tautou's mother. Probable parturition: 22th Feb 2016.<br>• Germine's mother. |

Dorsal view // 2018 - 02 - 06

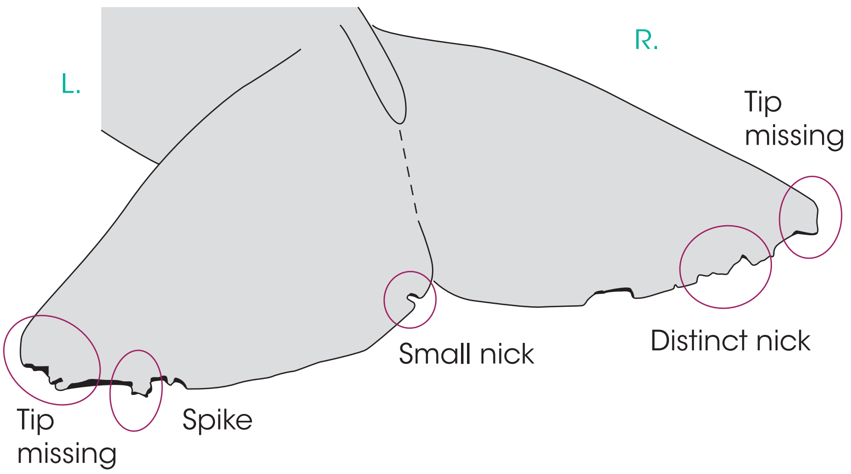

L. pectoral

Issa

◀ Front

L. pectoral

◀ Front

R. pectoral

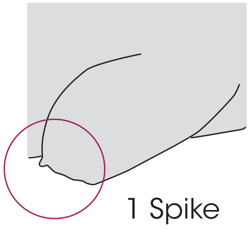

Front ▶

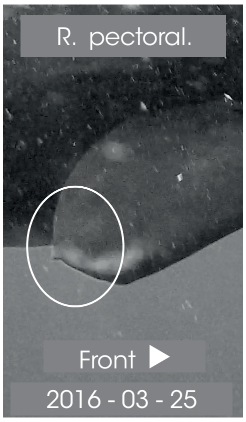

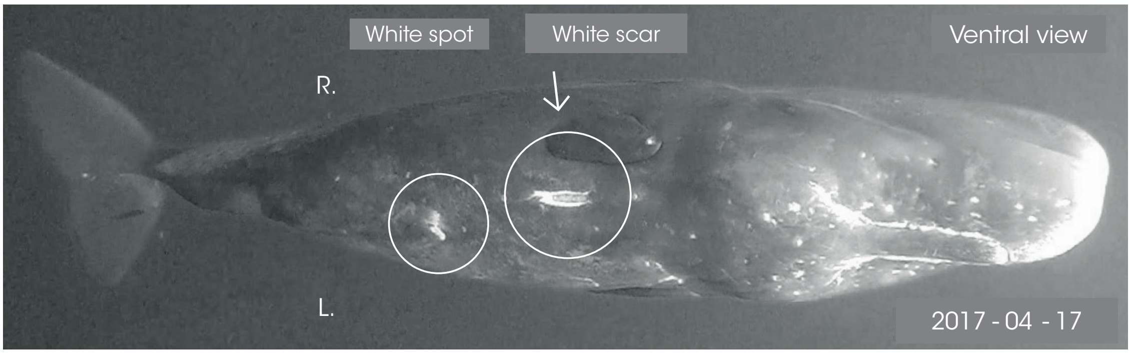

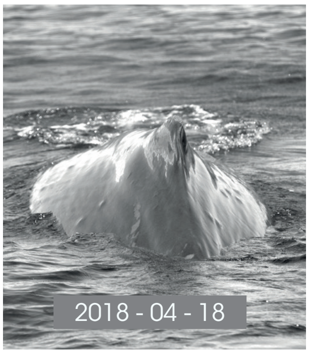

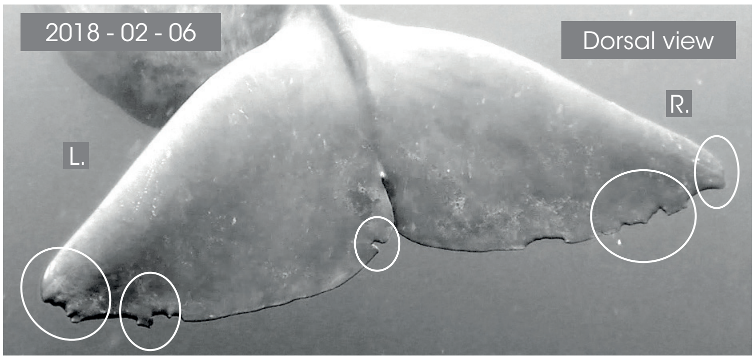

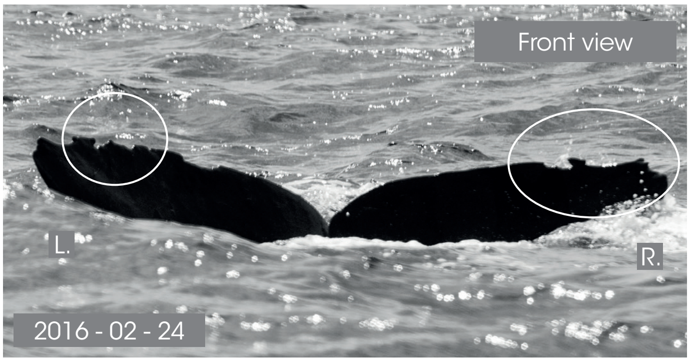

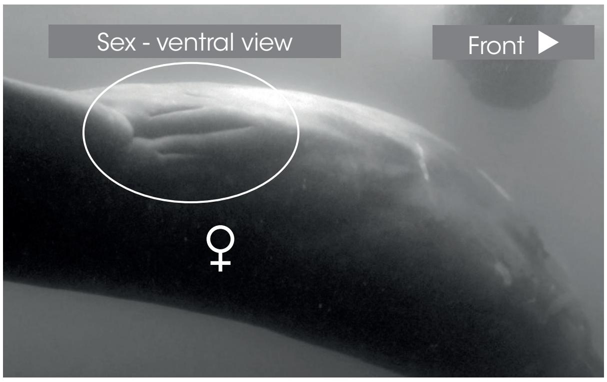

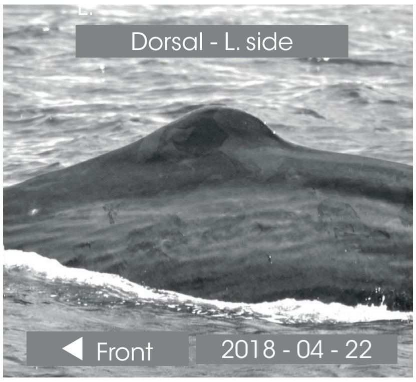

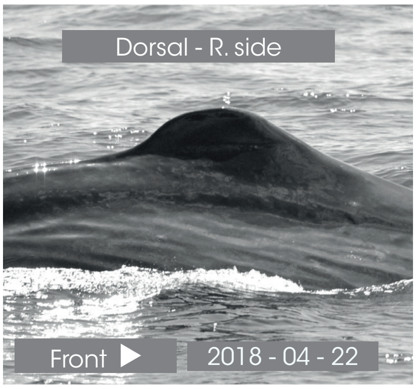

© Longitude 181 - Creation 2015 / Update 2020 - Concept : F. Sarano - Graphic design & illustrations : M. Sarano - © Pictures : S. Granzotto, F. Guérin, R. Heuzey, A. Preud'homme, V. & F. Sarano.

### JOUE BLANCHE

*1ST OBS. : 2009 - 01 - 27 // LAST OBS. : 2015 - 04 - 25*

| *ADULT* | *SEEN* | |
|---|---|---|
| - | ? - 2012<br>? - 2014<br>2015 - ?<br>? | *ADDITIONAL INFORMATION* |
| ♀ | | • White spot under the eye on the L. cheek.<br>• Wounds after Agatha's disappearance - 25th April 2015<br>• Convex caudal.<br>• Probable Agatha's mother. Parturition: 24th March 2014 |

L. pectoral

Joue blanche

◀ Front

L. pectoral

◀ Front

2015 - 05 - 02

R. pectoral

Joue blanche

Front ▶

R. pectoral.

Front ▶

Dorsal view // 2014 - 04 - 07

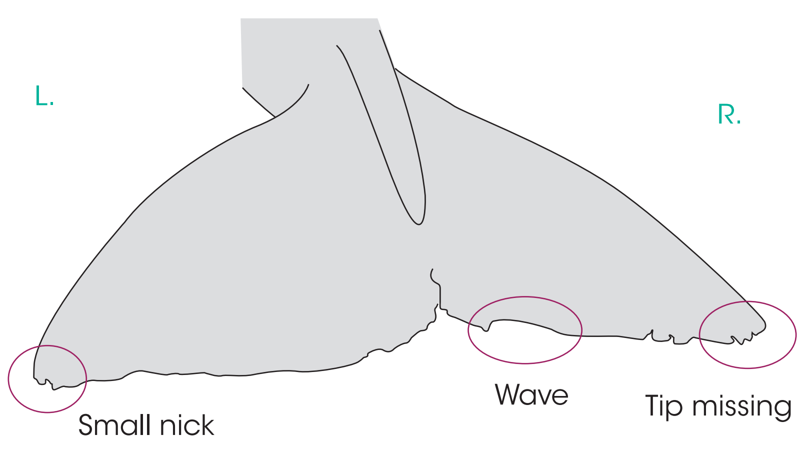

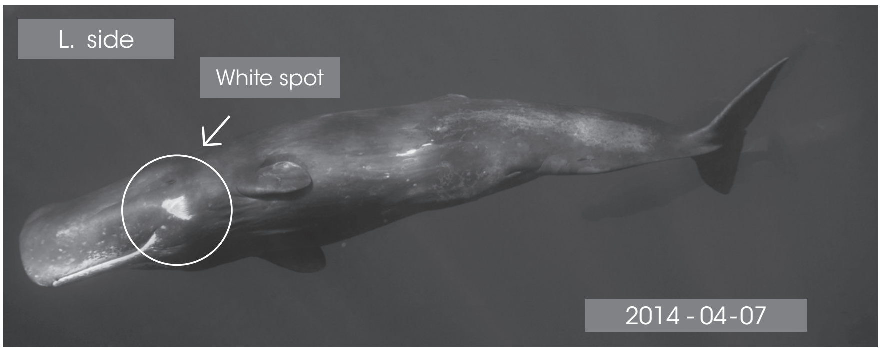

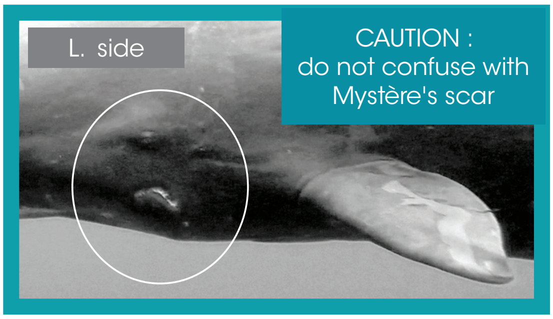

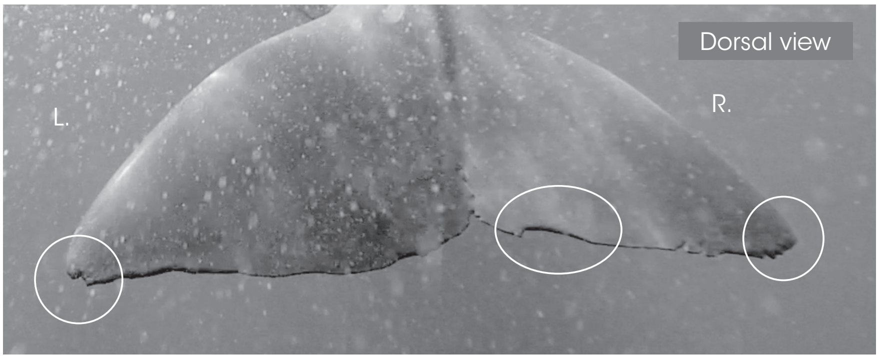

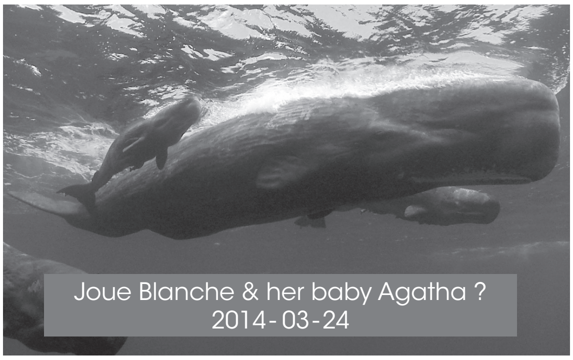

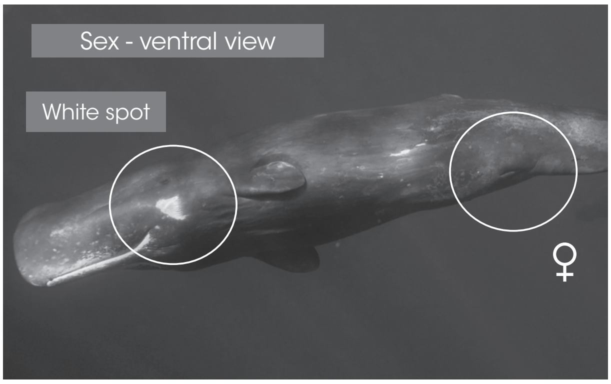

Dorsal - L. side

◀ Front

Dorsal - R. side

Front ▶

© Longitude 181 - Creation 2015 / Update 2020 - Concept : F. Sarano - Graphic design & illustrations : M. Sarano - © Pictures : S. Granzotto, F. Guérin, R. Heuzey, A. Preud'homme, V. & F. Sarano.

### LUCY

1ST OBS. : 2009 - 06 - 13 // LAST OBS. : 2020 - 03 - 11

| ADULTE | SEEN | |
|---|---|---|
| ADN | 2011 - 2012<br>2013 - 2014<br>2015 - 2016<br>2017 - 2018<br>2019 - 2020 | ADDITIONAL INFORMATION |
| ♀ | | • Very dark body color.<br>• 5 distinct nicks on the L. caudal lobe.<br>• Dos Calleux's daughter.<br>• Roméo's mother. Parturition: beginning March 2013.<br>• Daren's mother. Parturition:17? /04/2018 |

L. pectoral

Small nick

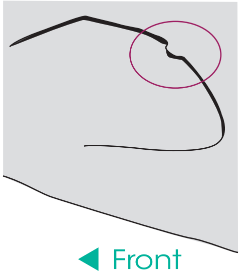

◀ Front

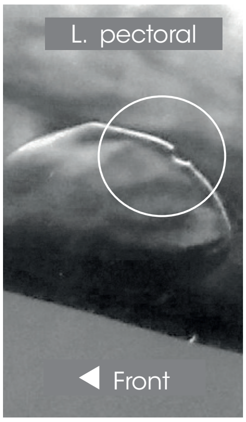

Dorsal view // 2014 - 04 - 26

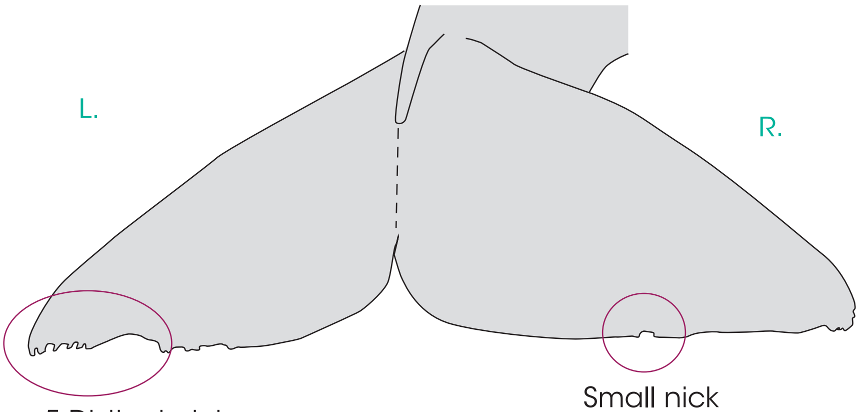

R. pectoral

Lucy

Front ▶

R. pectoral.

Front ▶

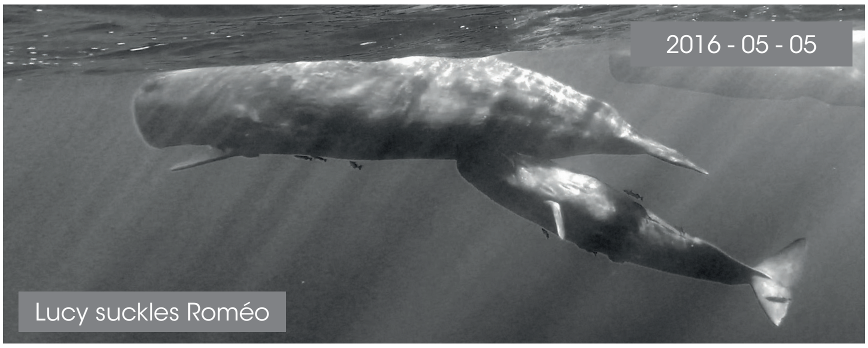

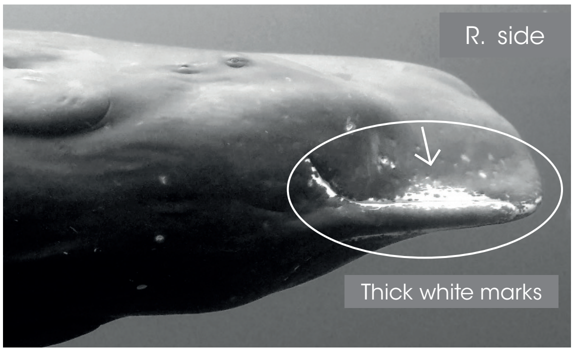

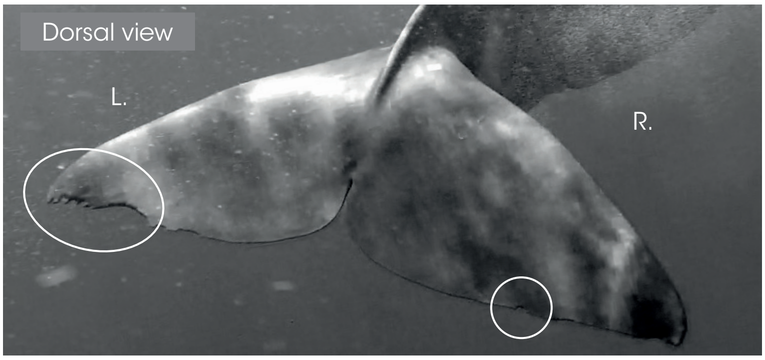

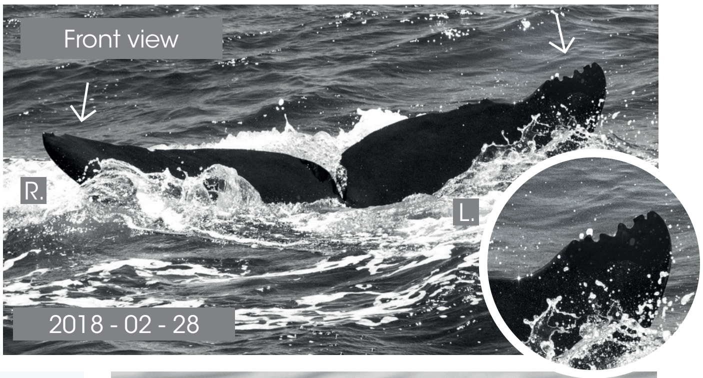

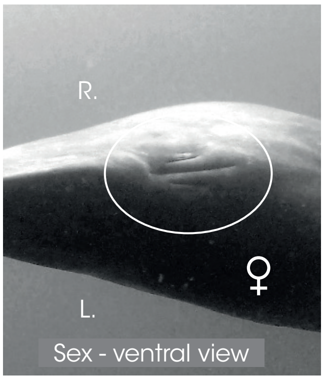

Dorsal - L. side

◀ Front

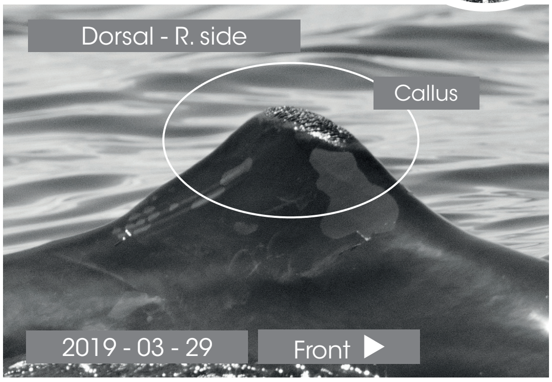

© Longitude 181 - Creation 2015 / Update 2020 - Concept : F. Sarano - Graphic design & illustrations : M. Sarano - © Pictures : S. Granzotto, F. Guérin, R. Heuzey, A. Preud'homme, V. & F. Sarano.

### MINA

**1ST OBS. : 2009 - 06 - 13 // LAST OBS. : 2020 - 03 - 12**

| ADULT | SEEN | |
|---|---|---|
| DNA | | ADDITIONAL INFORMATION |
| ♀ | 2011 - 2012<br>2013 - 2014<br>2015 - 2016<br>2017 - 2018<br>2019 - 2020 | • The 2nd scallop appeared in 2017.<br>• Concave caudal.<br>• Dos Calleux's daughter.<br>• Ali's mother - parturition: beginning of Feb.2018. |

L. pectoral

UNHARMED

◀ Front

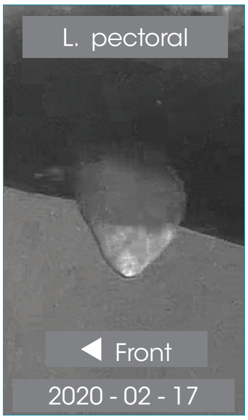

Dorsal view // 2017 - 03 - 14

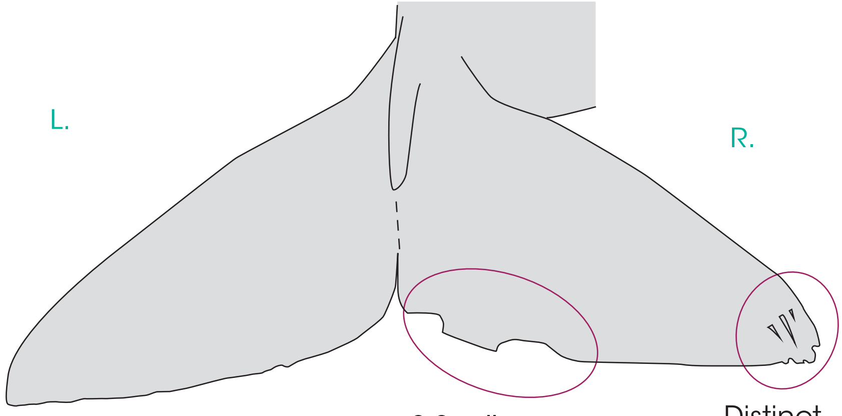

R. pectoral

Mina

Front ▶

R. pectoral.

Front ▶

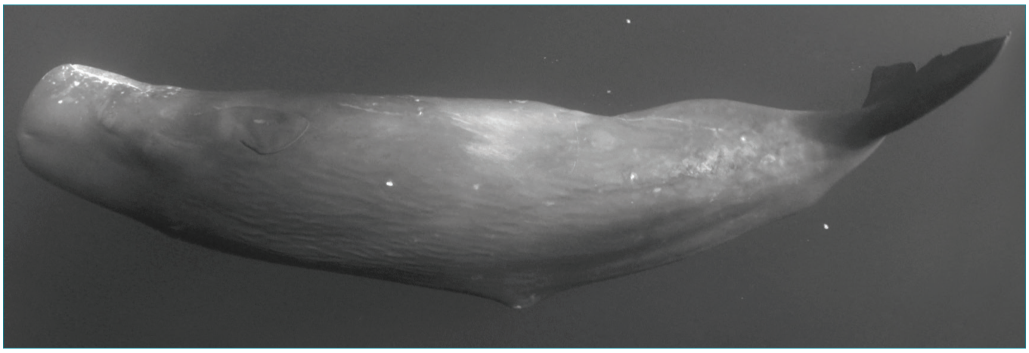

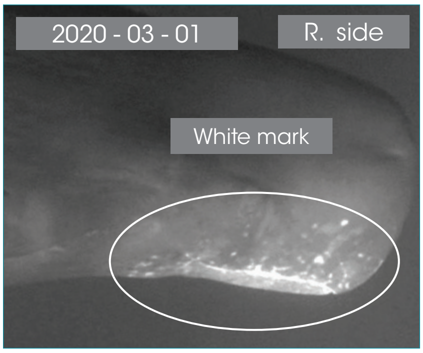

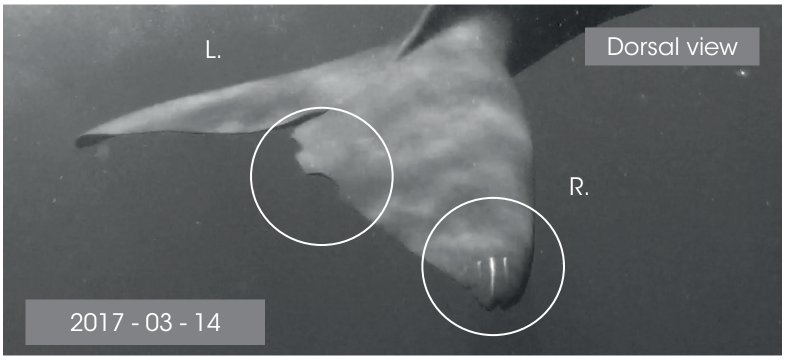

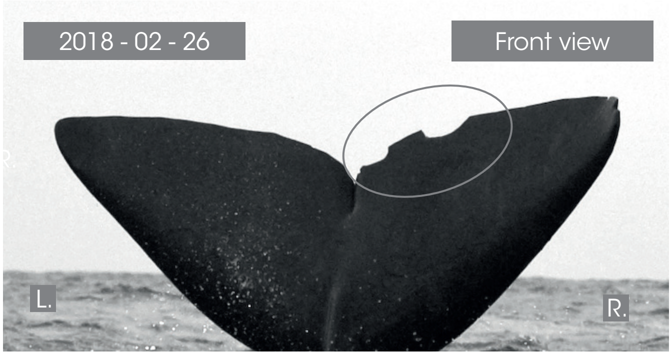

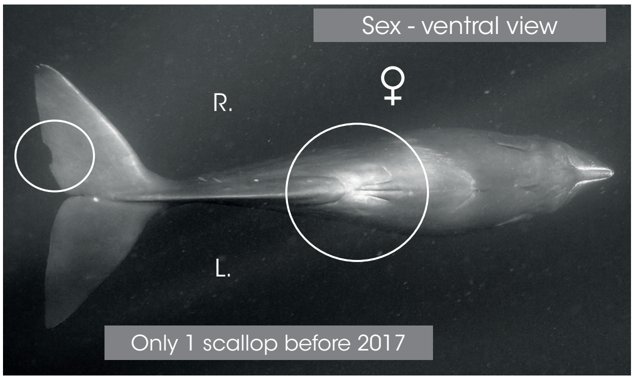

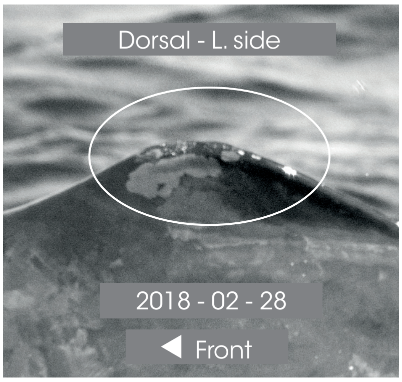

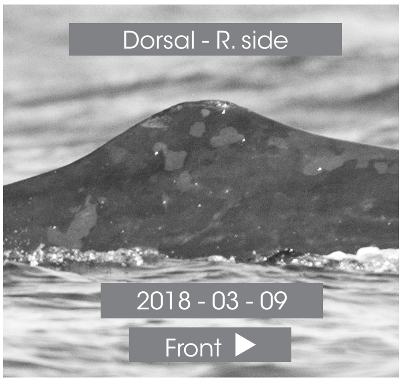

© Longitude 181 - Creation 2015 / Update 2020 - Concept : F. Sarano - Graphic design & illustrations : M. Sarano - © Pictures : S. Granzotto, F. Guérin, R. Heuzey, A. Preud'homme, V. & F. Sarano.

### MYSTÈRE

1ST OBS. : 2011 - 06 - 22 // LAST OBS. : 2020 - 02 - 17

| ADULT | SEEN | |
|---|---|---|
| DNA | 2011 - ?<br>? - ?<br>2015 - 2016<br>2017 - 2018<br>2019 - 2020 | ADDITIONAL INFORMATION |
| ♀ | | • 2 distinct nicks on the dorsal.<br>• White spot close to the genital slit.<br>• Scar on the L. cheek.<br>• Irène's true sister. |

L. pectoral

UNHARMED

◀ Front

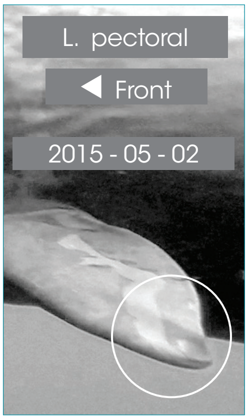

Dorsal view // 2016 - 05 - 06

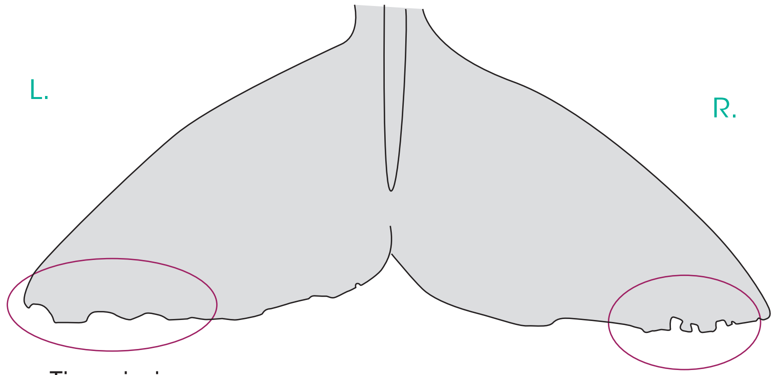

R. pectoral

Mystère

Front ▶

R. pectoral.

Front ▶

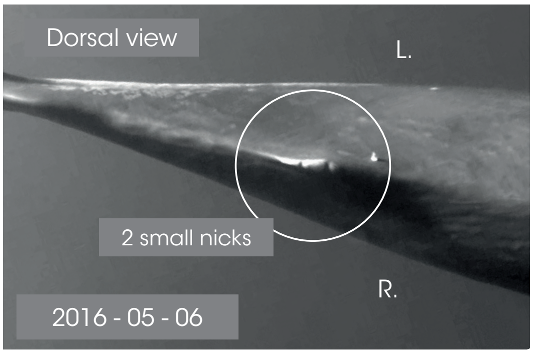

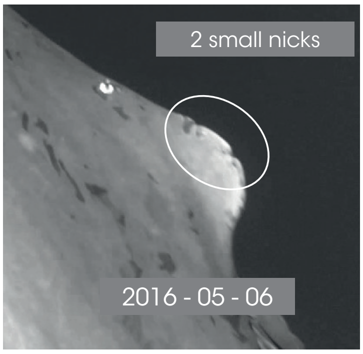

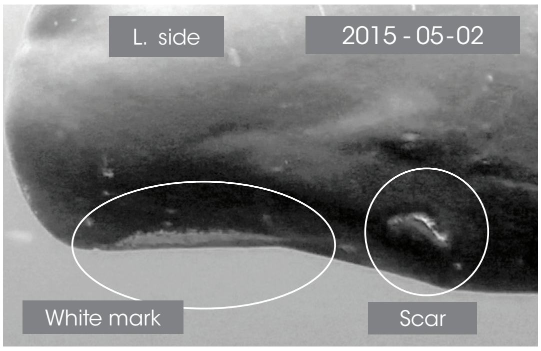

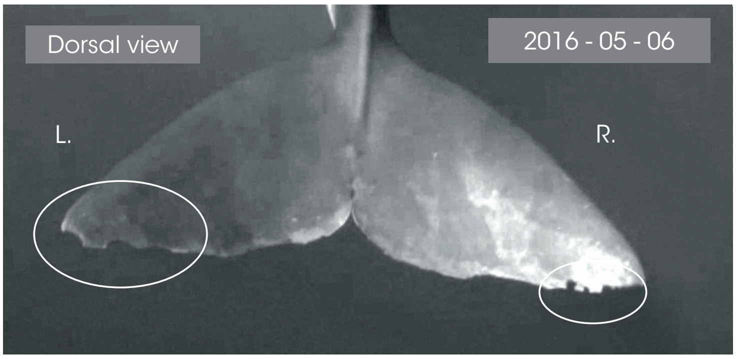

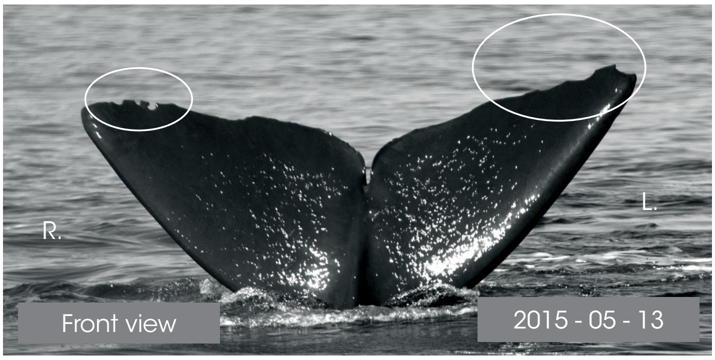

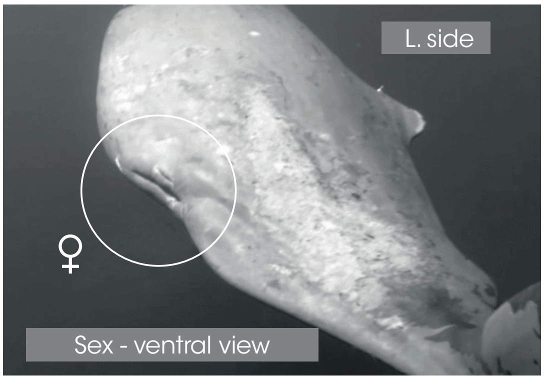

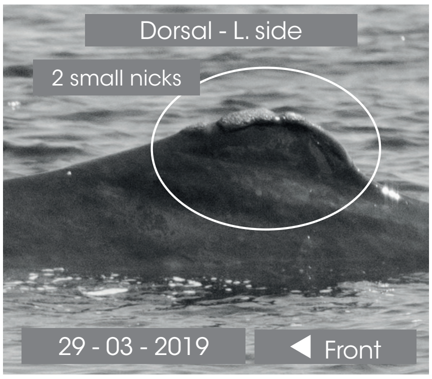

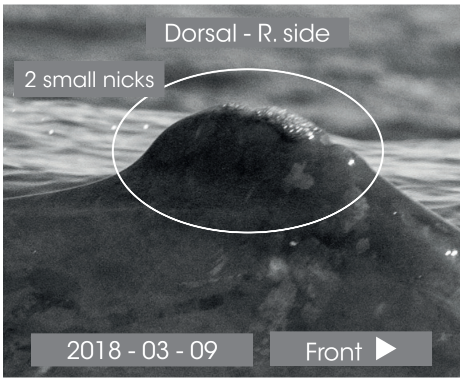

© Longitude 181 - Creation 2015 / Update 2020 - Concept : F. Sarano - Graphic design & illustrations : M. Sarano - © Pictures : S. Granzotto, F. Guérin, R. Heuzey, A. Preud'homme, V. & F. Sarano.

### SWASTEE

1ST OBS. : 2016 - 03 - 24 // LAST OBS. : 2019 - 04 - 26

| ADULT | SEEN | |
|---|---|---|
| - | ? - 2016<br>2017 - 2018<br>2019 | ADDITIONAL INFORMATION |
| ♀ | | • Huge bulge on the nape.<br>• Little hole in the L. caudal lobe. |

L. pectoral

UNHARMED

◀ Front

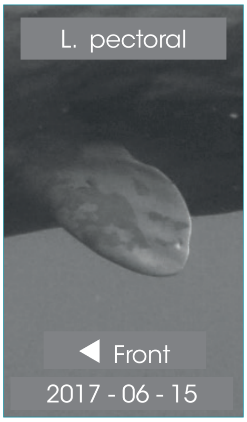

Dorsal view // 2017 - 04 - 16

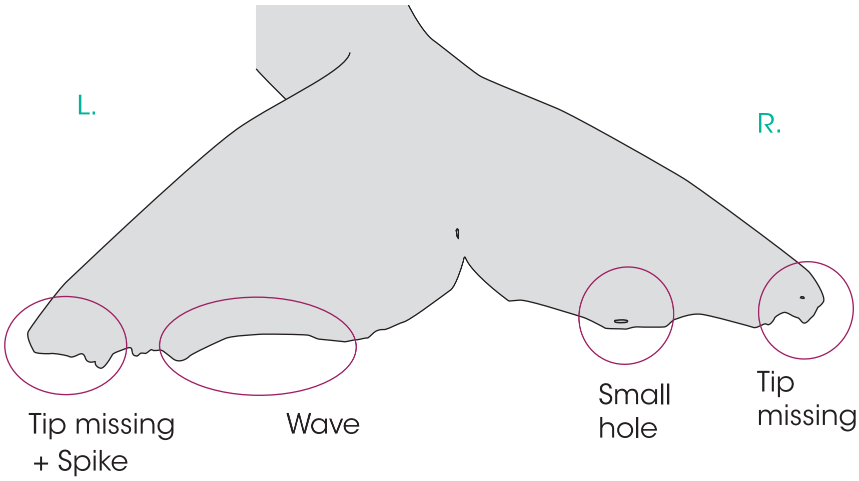

R. pectoral

Swastee

Front ▶

R. pectoral.

Front ▶

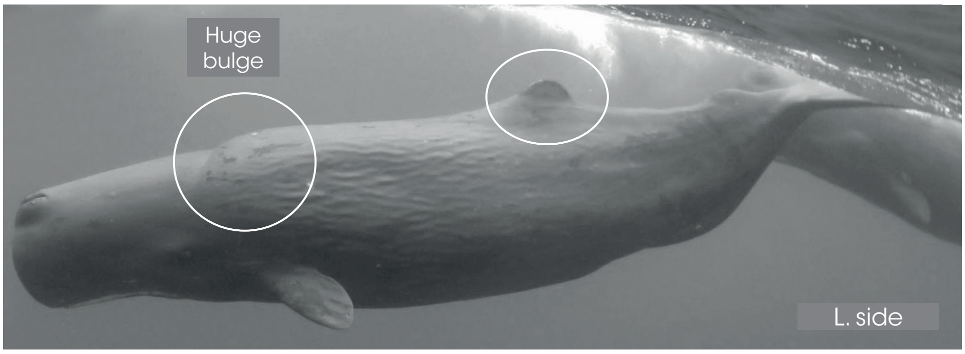

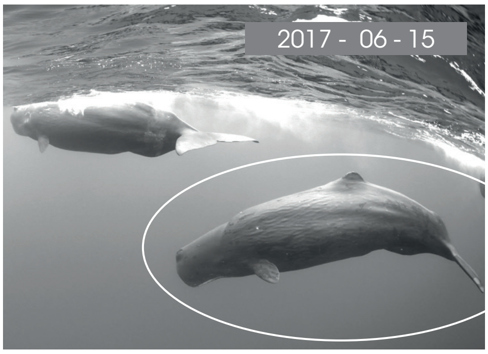

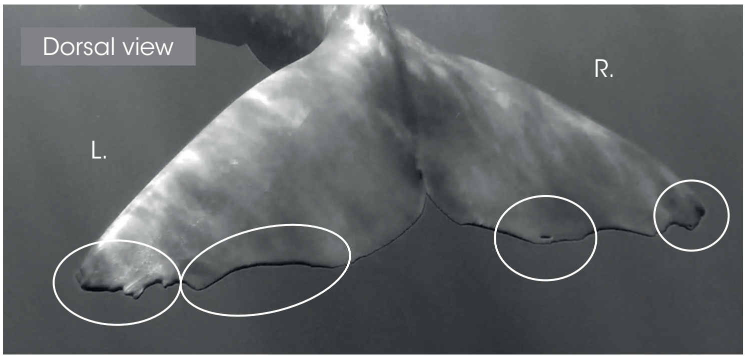

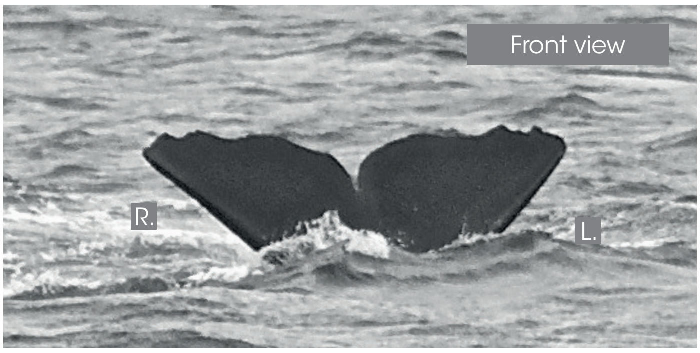

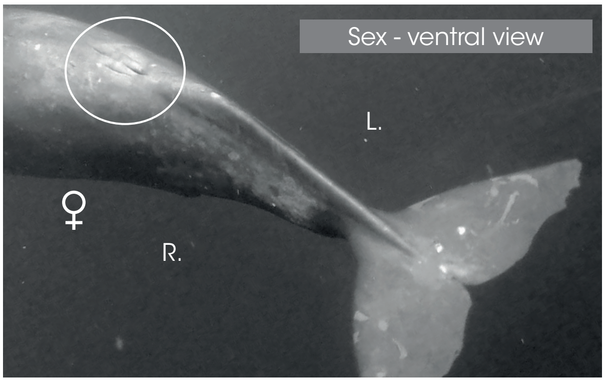

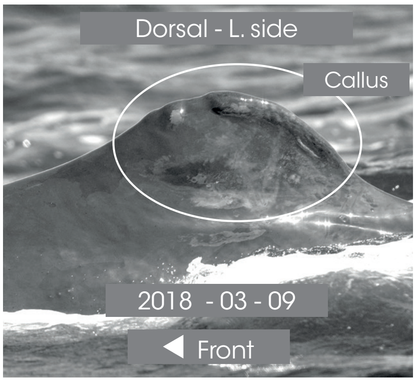

Dorsal - R. side

Front ▶

© Longitude 181 - Creation 2015 / Update 2020 - Concept : F. Sarano - Graphic design & illustrations : M. Sarano - © Pictures : S. Granzotto, F. Guérin, R. Heuzey, A. Preud'homme, V. & F. Sarano.

### VANESSA

**1ST OBS. : 2012 - 01 - 12 // LAST OBS. : 2020 - 03 - 18**

| ADULT | SEEN | |
|---|---|---|
| DNA | ? - ?<br>? - 2014<br>2015 - 2016<br>2017 - 2018<br>2019 - 2020 | ADDITIONAL INFORMATION |
| ♀ | | • White callus on the top of the dorsal, noticeable.<br>• Once known as Piton. |

Dorsal view // 2015 - 04 - 21

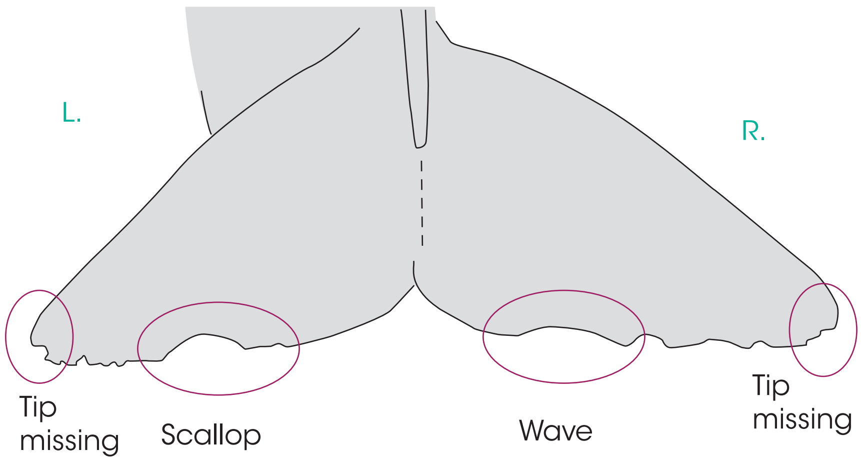

L. pectoral

UNHARMED

◀ Front

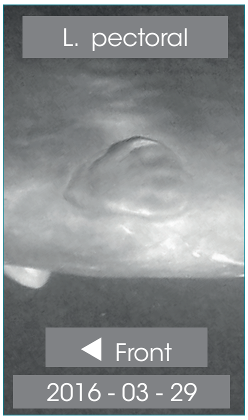

R. pectoral

UNHARMED

Front ▶

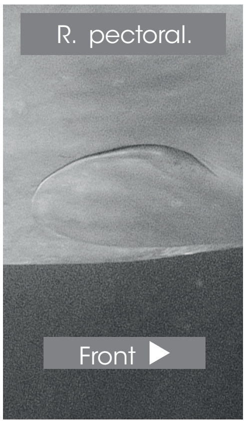

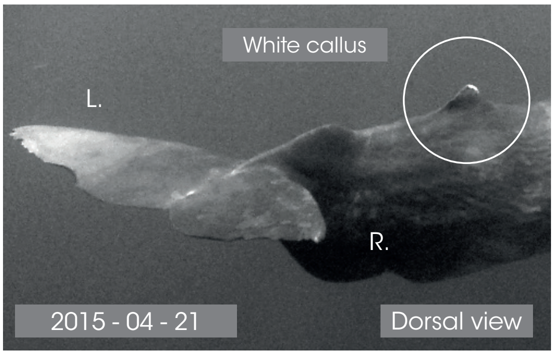

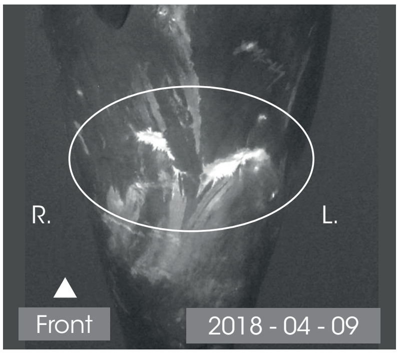

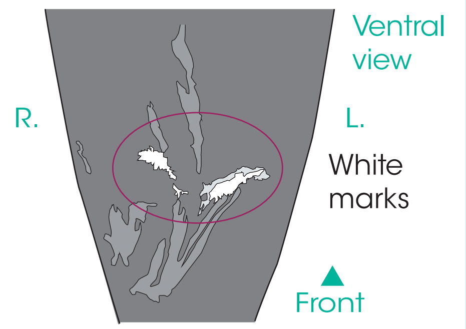

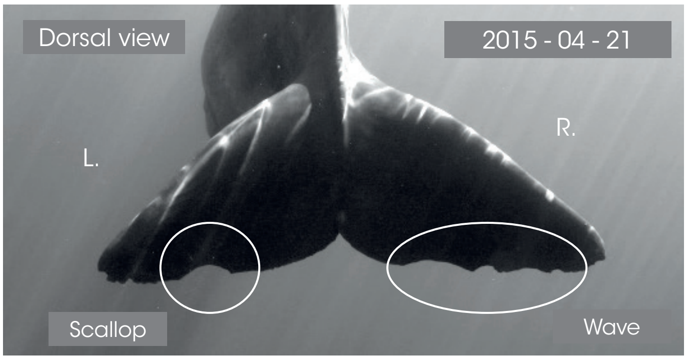

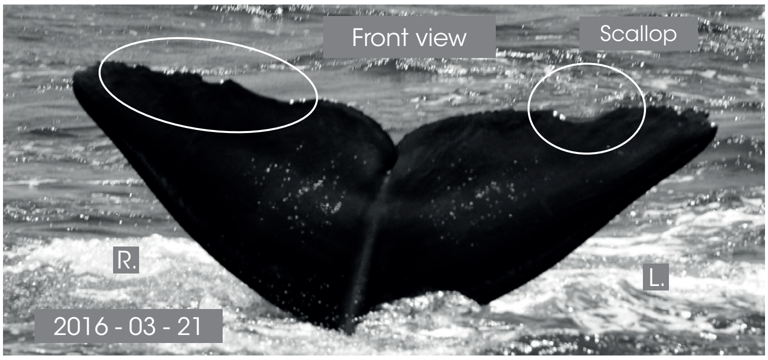

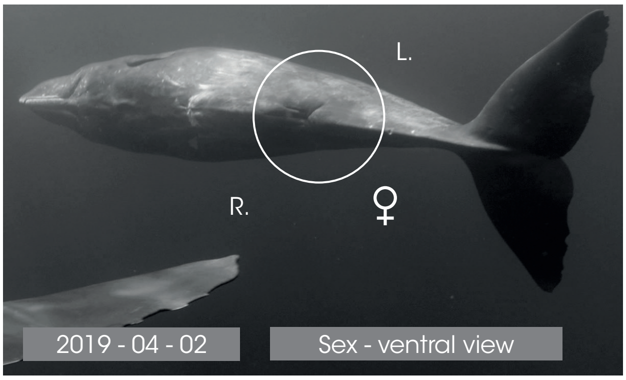

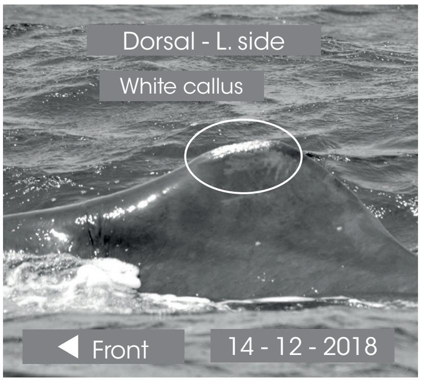

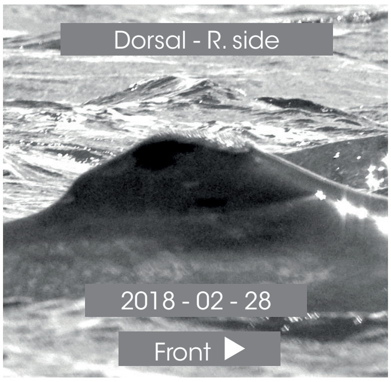

© Longitude 181 - Creation 2015 / Update 2020 - Concept : F. Sarano - Graphic design & illustrations : M. Sarano - © Pictures : S. Granzotto, F. Guérin, R. Heuzey, A. Preud'homme, V. & F. Sarano.

### YUKIMI

1ÈRE OBS. : 2011 - 03 - 14 // DERNIÈRE OBS. : 2020 - 03 - 10

| ADULT | SEEN | |
|---|---|---|
| DNA | 2011 - ?<br>? - ?<br>2015 - 2016<br>2017 - 2018<br>2019 - 2020 | ADDITIONAL INFORMATION |
| ♀ | | • Very dark mandibular zone, without white marks.<br>• Convex caudal.<br>• Ali's nurse in 2018.<br>• Lana's nurse in 2019, with evidence of milk. |

L. pectoral

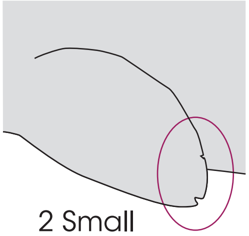

2 Small nicks

◀ Front

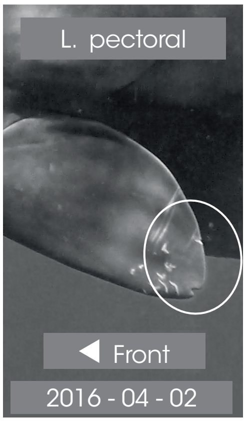

Dorsal view // 2016 - 02 - 24

L.

R.

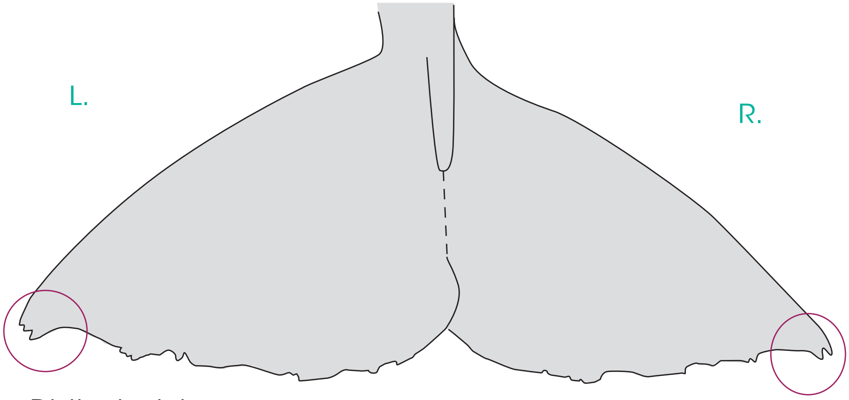

Distinck nicks

Distinck nick

R. pectoral

Yukimi

Front ▶

R. pectoral.

Front ▶

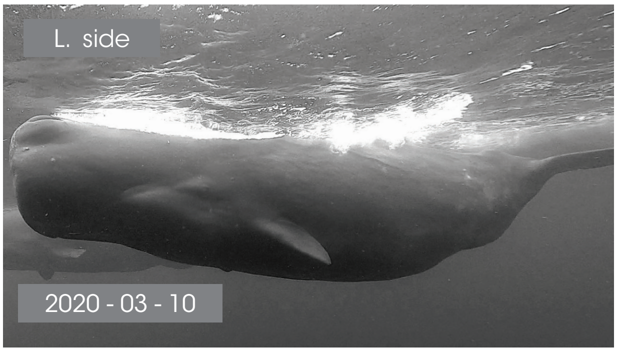

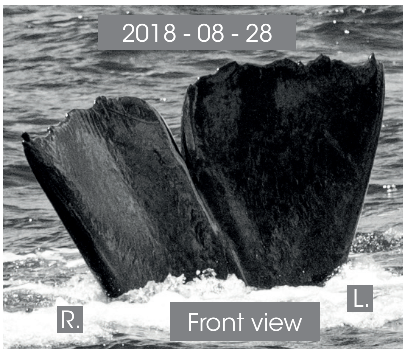

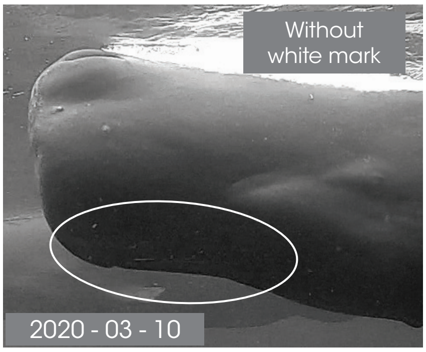

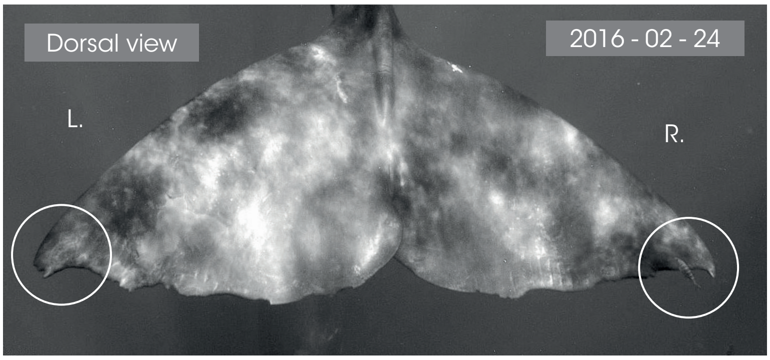

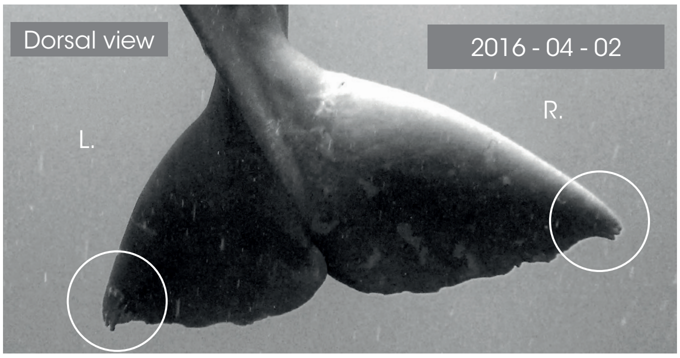

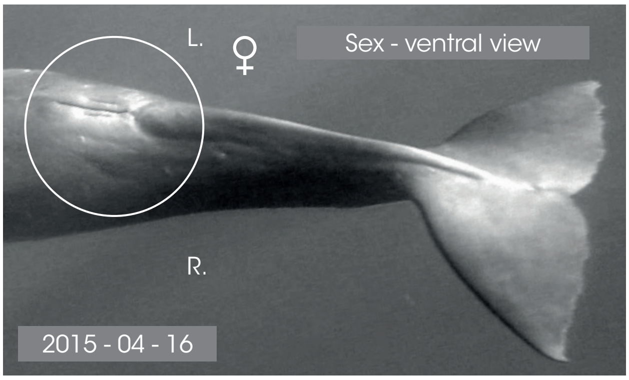

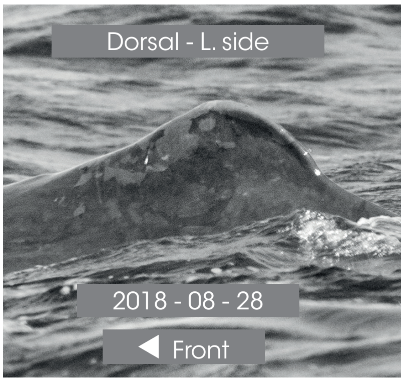

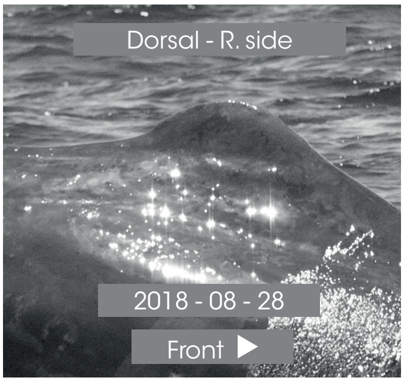

© Longitude 181 - Creation 2015 / Update 2020 - Concept : F. Sarano - Graphic design & illustrations : M. Sarano - © Pictures : S. Granzotto, F. Guérin, R. Heuzey, A. Preud'homme, V. & F. Sarano.

### JUVENILES

CONCEPT : François Sarano

GRAPHIC DESIGN & ILLUSTRATIONS : Marion Sarano

PICTURES : Stéphane Granzotto, Fabrice Guérin, René Heuzey, Daniel Jouannet, Bernard Kirchhofer, Vanessa Mignon, Axel Preud'homme, Véronique Sarano, François Sarano, Hugues Vitry.

© Longitude 181 - Creation 2015 / Update 2020

### IDENTITY CARDS
### SPERM WHALES OF MAURITIUS I.

#### AGATHA

*1ST OBS. : 2014 - 03 - 24 // LAST OBS. : 2015 - 04 - 17*

| JUVENILE | SEEN | B : 2014 - 20TH MARCH |
|---|---|---|
| DNA | 2014 - 2015 | *ADDITIONAL INFORMATION* |
| ♀ | DEAD ? | • R. caudal tip: missing portion<br>• L. caudal tip: tooth marks<br>• Still in the crèche on 17th April 2015, missing since. |

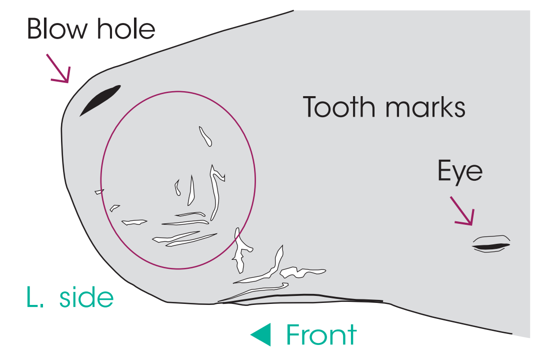

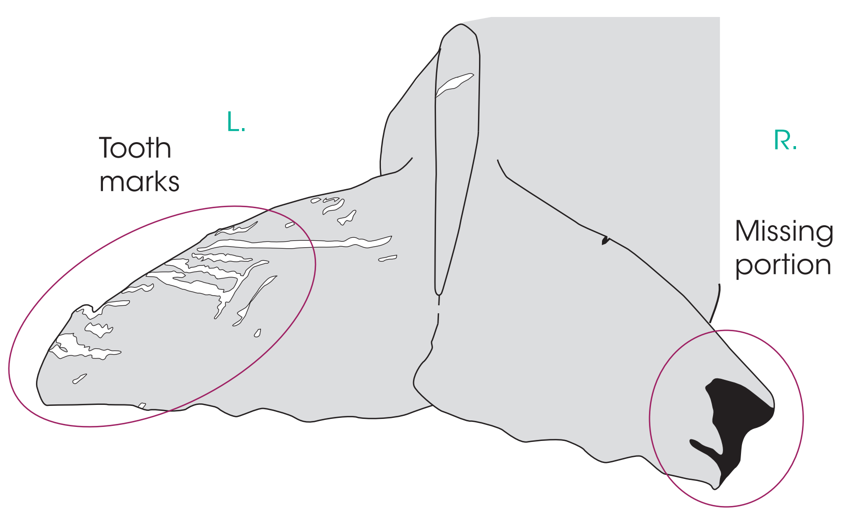

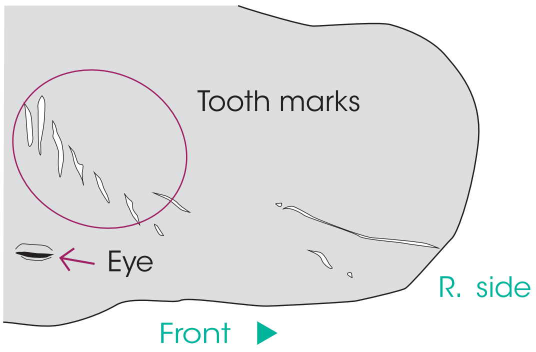

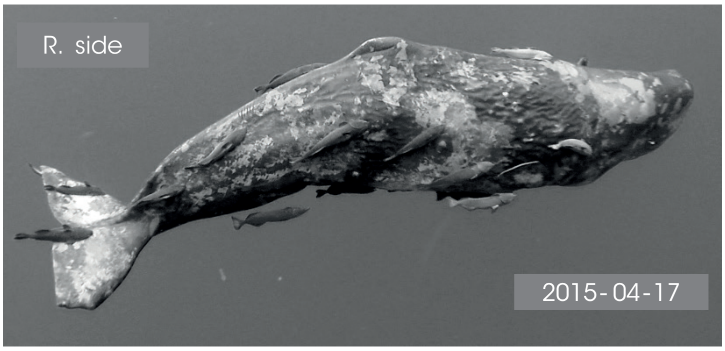

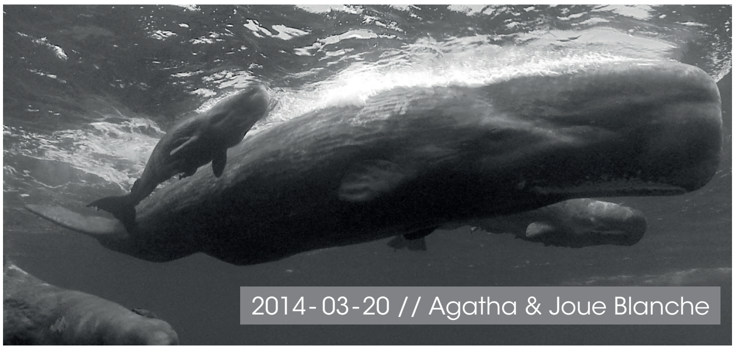

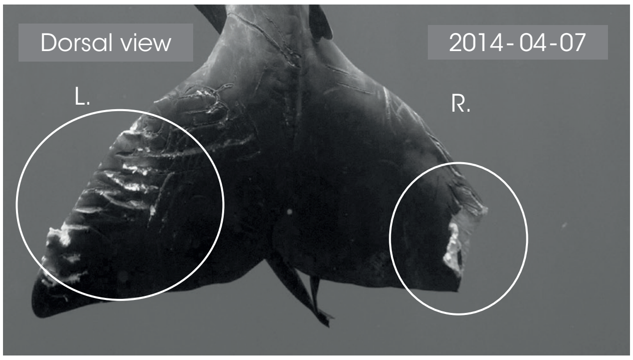

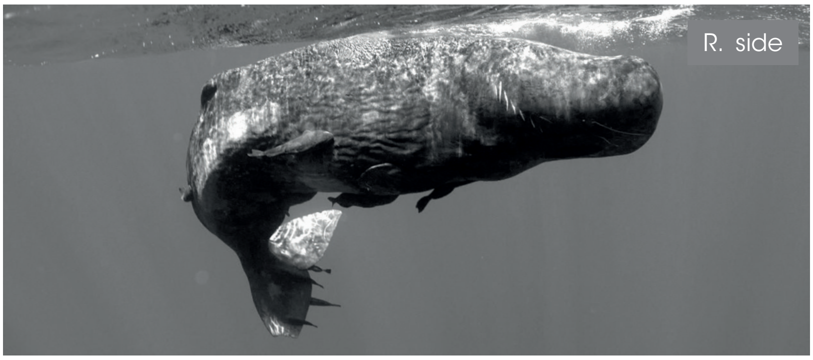

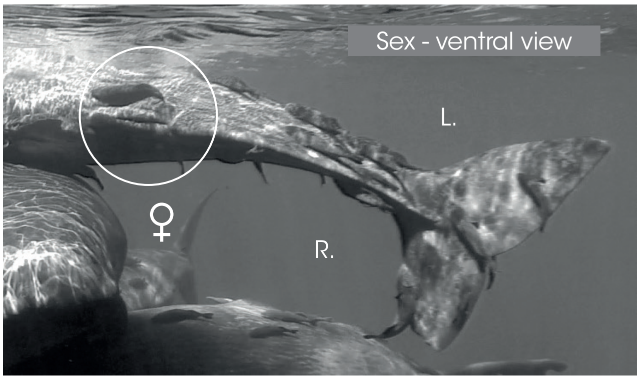

Dorsal - L. side

◀ Front

Dorsal - R. side

Front ▶

© Longitude 181 - Creation 2015 / Update 2020 - Concept : F. Sarano - Graphic design & illustrations : M. Sarano - © Pictures : S. Granzotto, F. Guérin, R. Heuzey, A. Preud'homme, V. & F. Sarano.

### ALEXANDER

*1ST OBS. : 2019 - 03 - 05 // LAST OBS. : 2020 - 03 - 18*

| JUVENILE | *SEEN* | B : 2019 - JAN |
|---|---|---|
| *DNA* | ... 2019 - 2020 | *ADDITIONAL INFORMATION* |
| ♂ | | • Thick white mark on the mandibular zone.<br>• Probably born : Dec. 2018 or Jan. 2019<br>• Caroline's son, Zoé's half-brother, Jonas's grandson<br>• Emy was its nurse in 2019 |

L. pectoral

**UNHARMED**

◀ Front

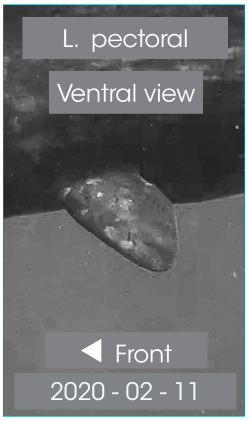

Dorsal view // 2020 - 03 - 02

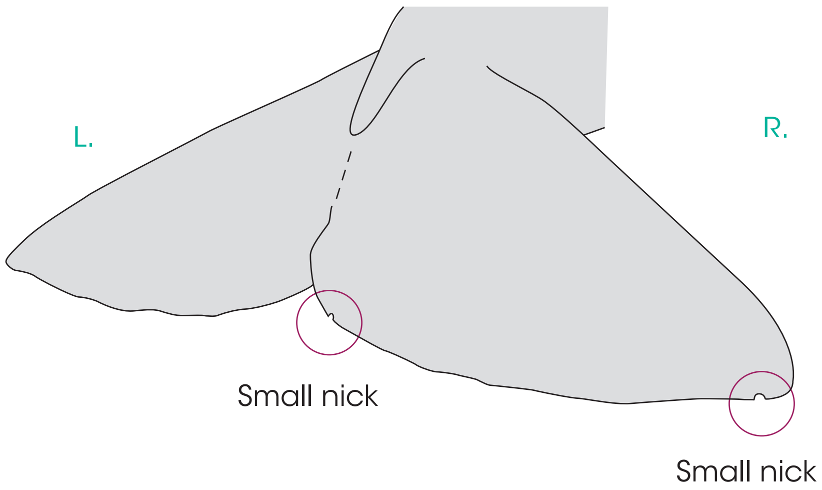

R. pectoral

Alexander

Front ▶

R. pectoral.

Front ▶

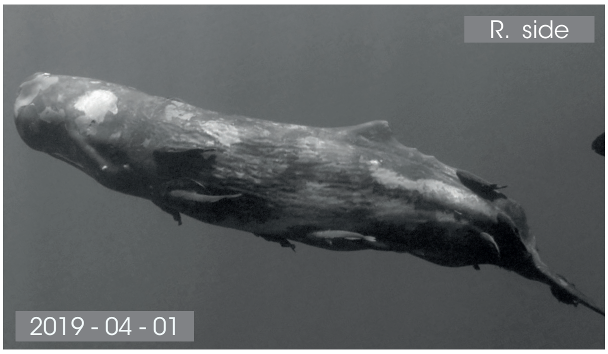

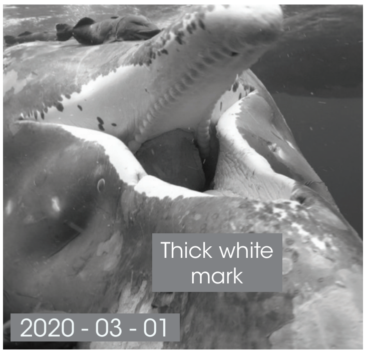

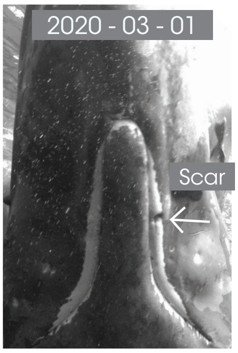

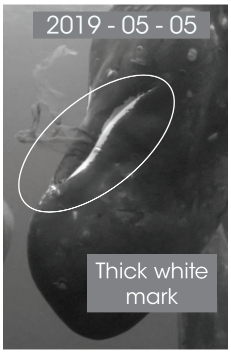

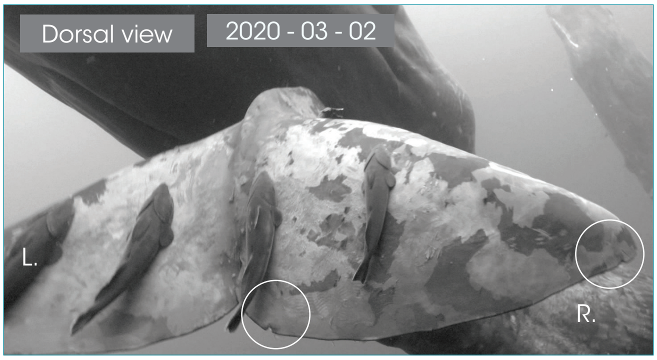

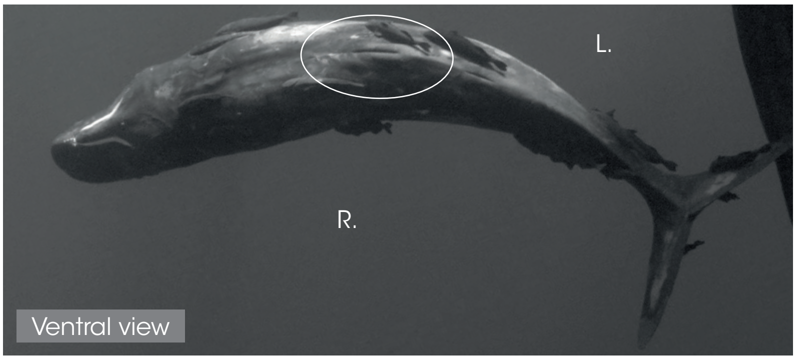

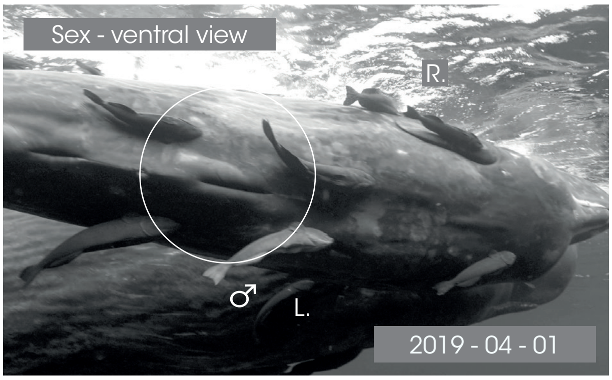

Dorsal - L. side

◀ Front

Dorsal - R. side

Front ▶

### ALI

#### 1ST OBS. : 2018-02-02 // LAST OBS. : 2020-03-12

| JUVENILE | SEEN | B : ≈ 2018 - FEB |
|---|---|---|
| DNA | - 2018<br>2019 - 2020 | ADDITIONAL INFORMATION |
| ♂ | | • White mandibular mark: black mottled.<br>• Large white stripes on the caudal.<br>• Mina's son.<br>• Dos Calleux's grandson. |

L. pectoral

**UNHARMED**

◀ Front

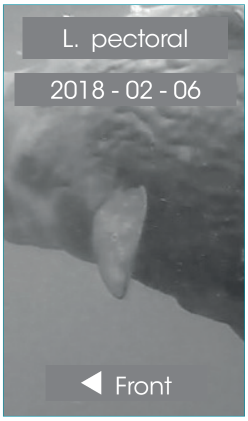

R. pectoral

**UNHARMED**

Front ▶

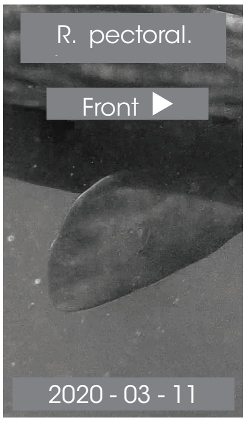

Dorsal view // 2018 - 04 - 07

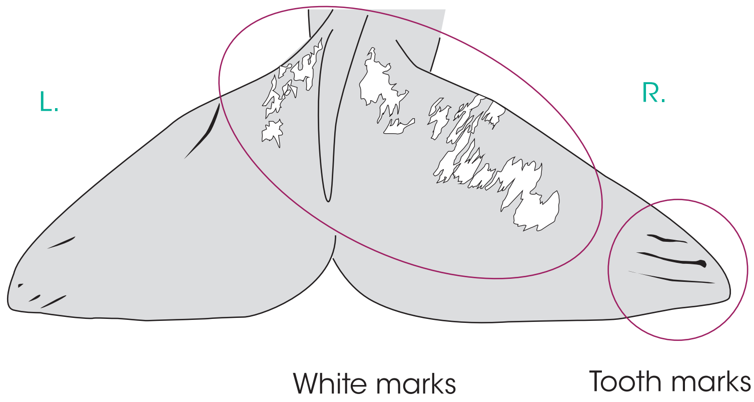

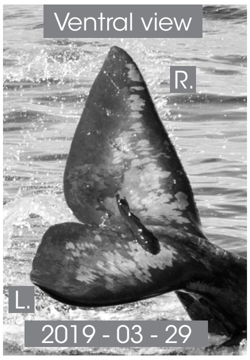

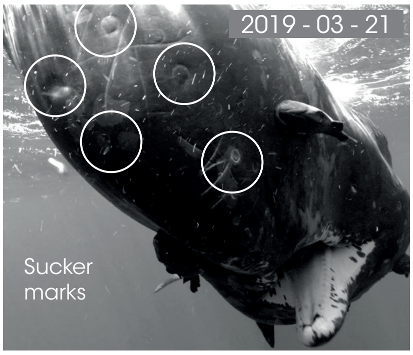

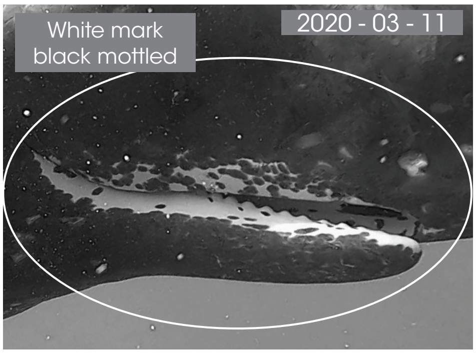

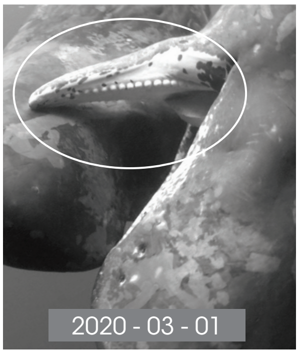

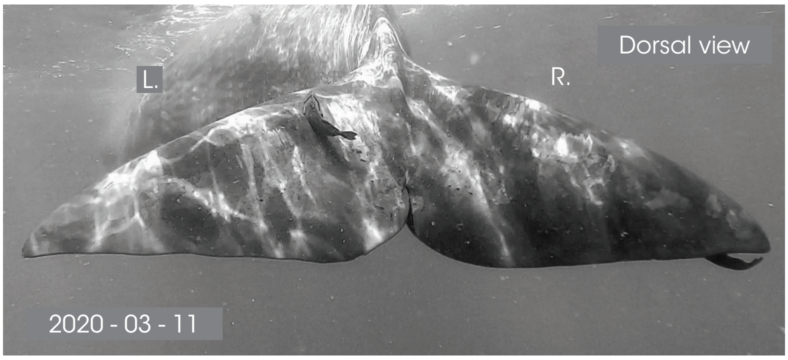

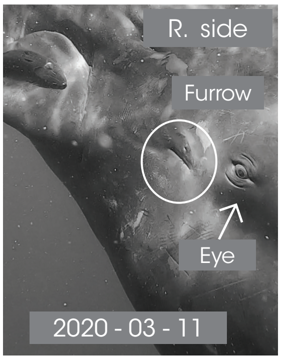

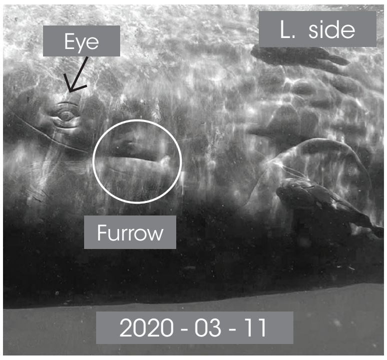

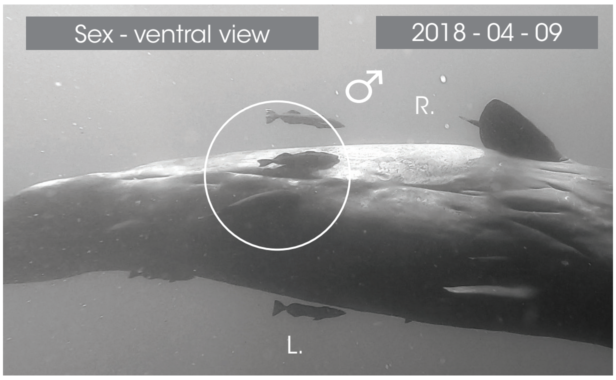

Dorsal - L. side.

◀ Front

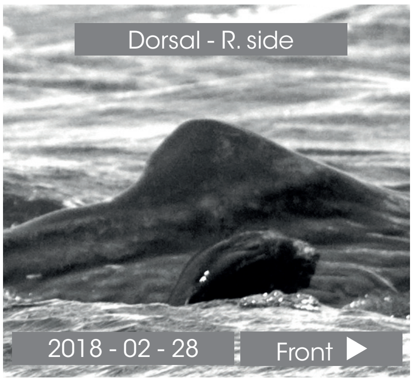

© Longitude 181 - Creation 2015 / Update 2020 - Concept : F. Sarano - Graphic design & illustrations : M. Sarano - © Pictures : S. Granzotto, F. Guérin, R. Heuzey, A. Preud'homme, V. & F. Sarano.

### ARTHUR

*1[ST] OBS. : 2013 - 04 - 05 // LAST OBS. : 2020 - 03 - 12*

| JUVENILE | *SEEN* | B : ≈ 2013 - 05[TH] APRIL |
|---|---|---|
| *DNA* | 2013 - 2014<br>2015 - 2016<br>2017 - 2018<br>2019 - 2020 | *ADDITIONAL INFORMATION* |
| ♂ | | • Bear ear shaped dorsal, protruding vertebras<br>• Ventral white escutcheon<br>• Irène's son.<br>• Lana's half-brother.<br>• Ker2007's great grandson. |

Dorsal view // 2018 - 04 - 12

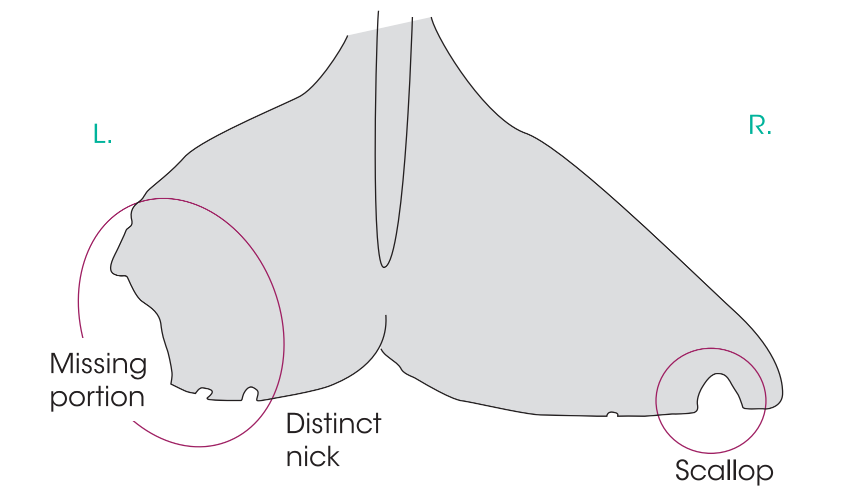

L. pectoral

UNHARMED

◀ Front

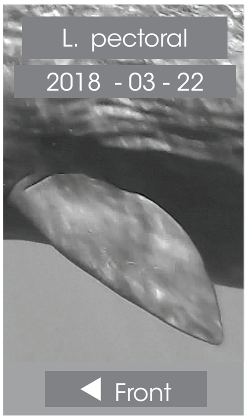

R. pectoral

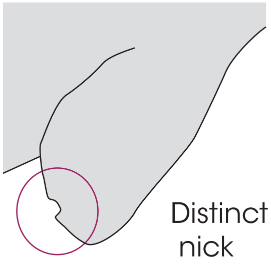

Front ▶

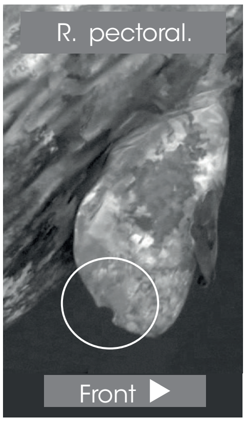

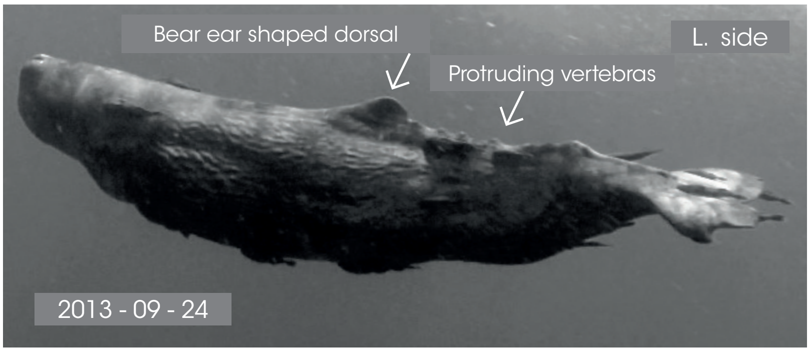

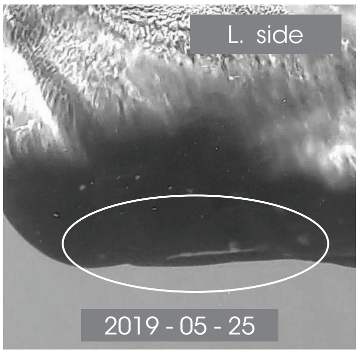

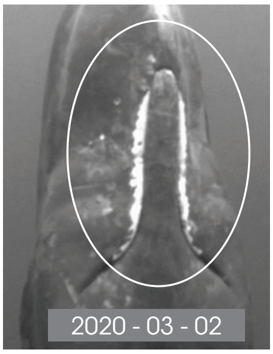

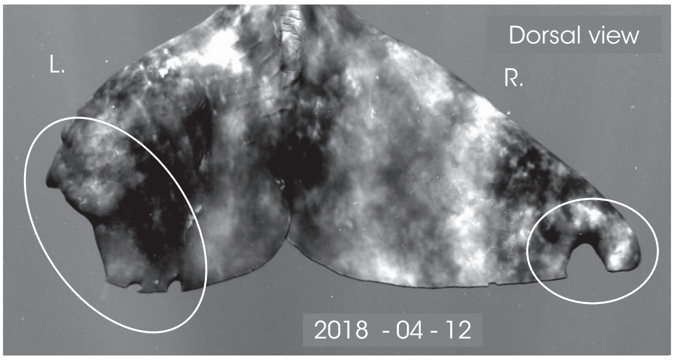

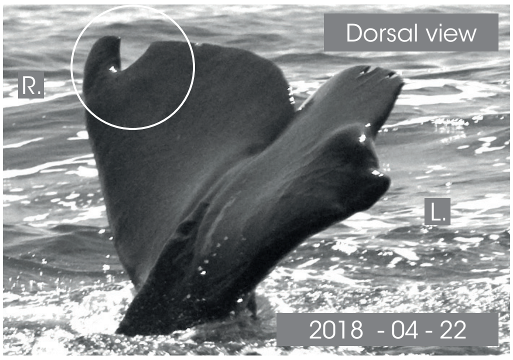

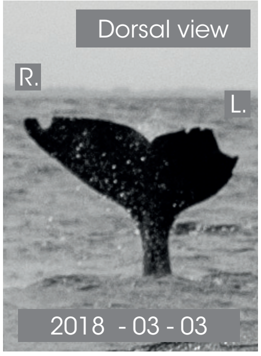

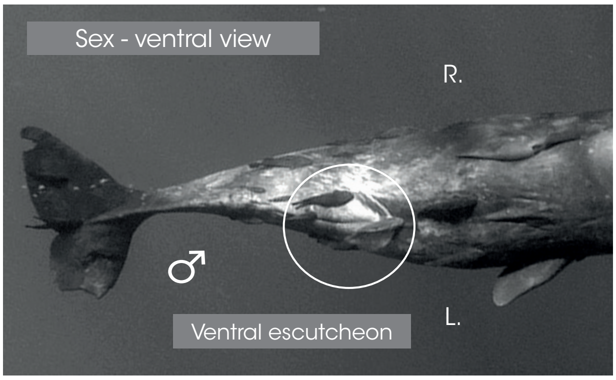

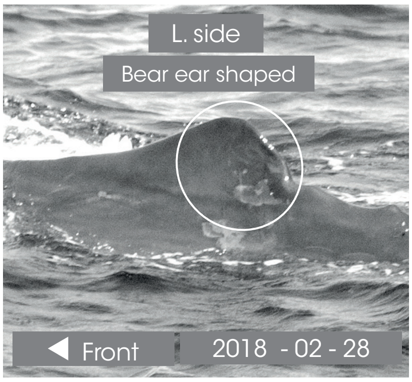

Dorsal - R. side

Front ▶

© Longitude 181 - Creation 2015 / Update 2020 - Concept : F. Sarano - Graphic design & illustrations : M. Sarano - © Pictures : S. Granzotto, F. Guérin, R. Heuzey, A. Preud'homme, V. & F. Sarano.

### BAPTISTE

1ST OBS. : 2017 - 03 - 11 // LAST OBS. : 2017 - 03 - 24

| | | |
|---|---|---|
| JUVENILE | SEEN | B : ≈ 2017 - 07TH MARCH |
| DNA | 2017 | ADDITIONAL INFORMATION |
| ♂ | DEAD ? | • 1 White spot on the back, in the front of the dorsal.<br>• Dos Calleux's son.<br>• Missing since 24th March 2017 |

L. pectoral

baptiste

◀ Front

L. pectoral

◀ Front

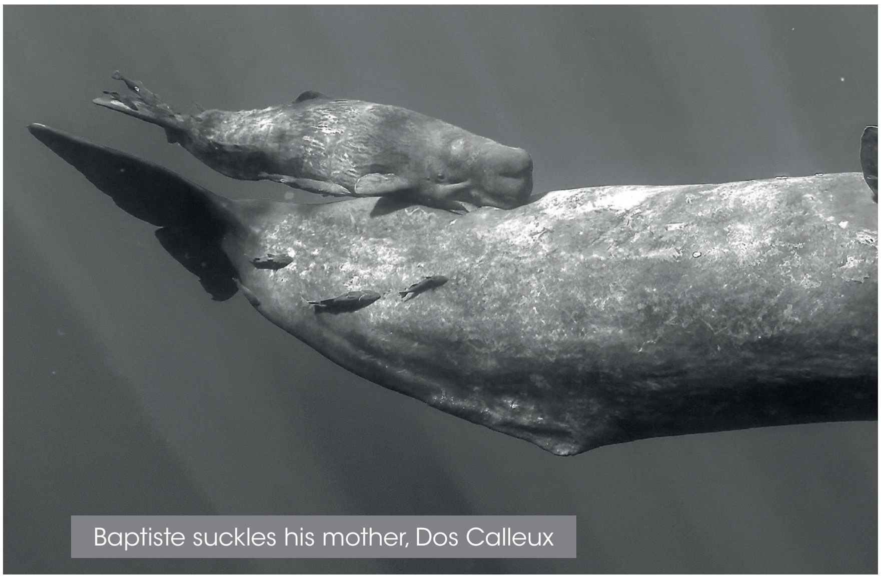
Baptiste suckles his mother, Dos Calleux

R. pectoral

UNHARMED

Front ▶

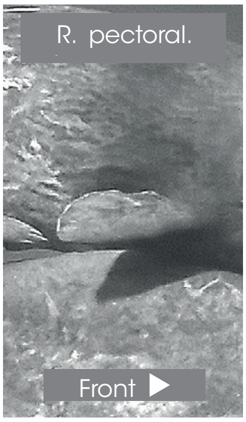

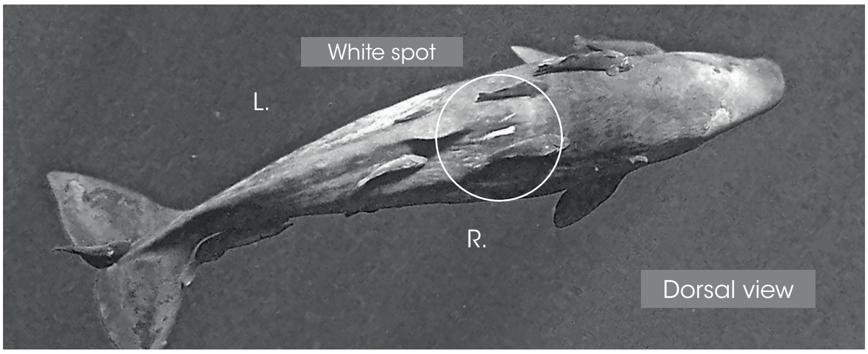

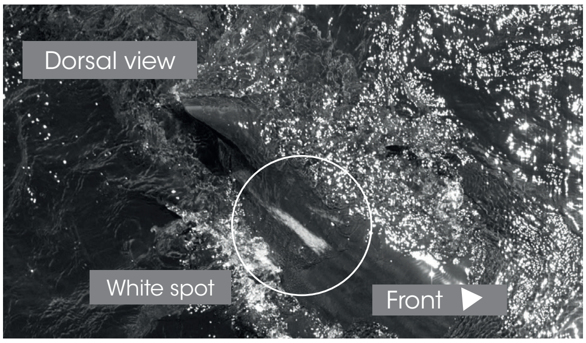

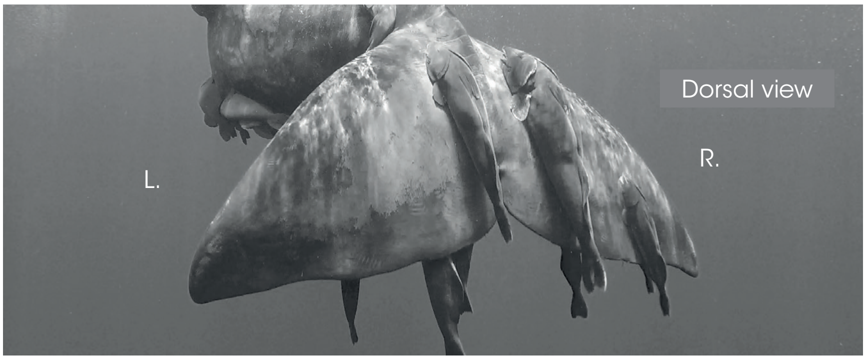

Dorsal view

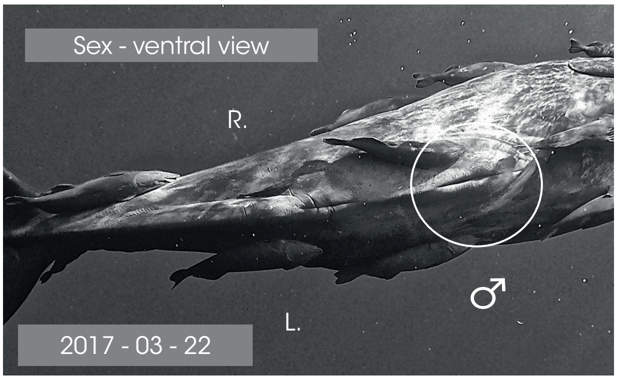

2017 - 03 - 22

Dorsal - L. side

◀ Front

Dorsal - R. side

Front ▶

© Longitude 181 - Creation 2015 / Update 2020 - Concept : F. Sarano - Graphic design & illustrations : M. Sarano - © Pictures : S. Granzotto, F. Guérin, R. Heuzey, A. Preud'homme, V. & F. Sarano.

LONGITUDE 181
La Voix de l'Océan

### IDENTITY CARDS
### SPERM WHALES OF MAURITIUS I.

#### CHESNA

1ST OBS. : 2018 - 03 - 02 // LAST OBS. : 2020 - 03 - 18

| JUVENILE | SEEN | B : ≈ 2018 - END FEB |
|---|---|---|
| DNA | - 2018<br>2019 - 2020 | ADDITIONAL INFORMATION |
| ♀ | | • L. caudal tip : missing portion (3-9th March 2018)<br>• Small nick on the R. pectoral (2nd April 2018)<br>• Delphine's daughter.<br>• Tache Blanche's half-sister. |

Dorsal view // 2018 - 04 - 01

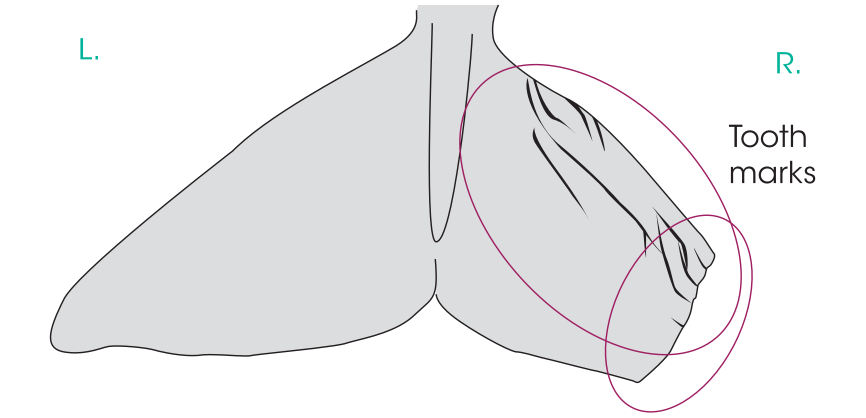

L. pectoral

UNHARMED

◀ Front

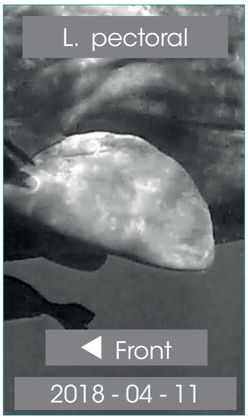

R. pectoral

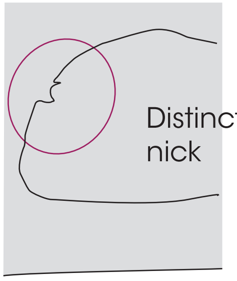

Front ▶

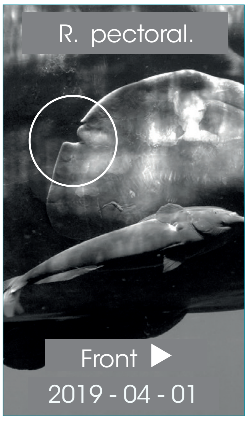

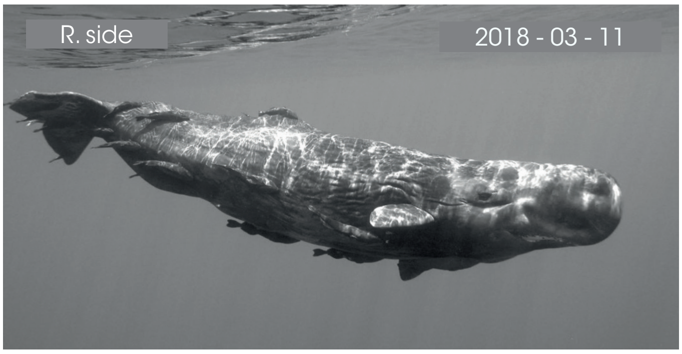

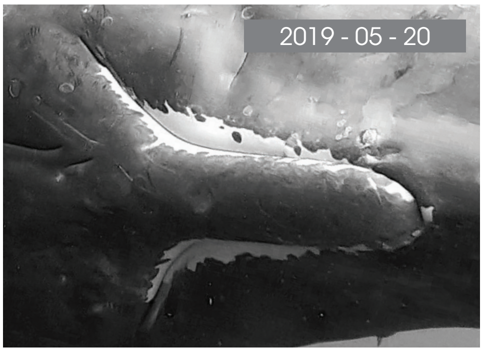

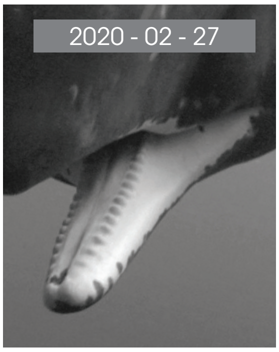

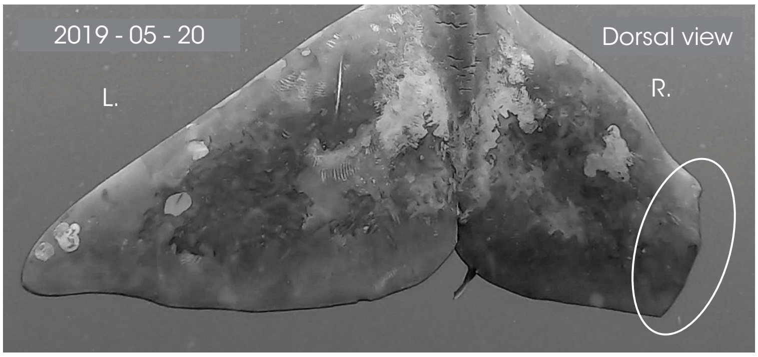

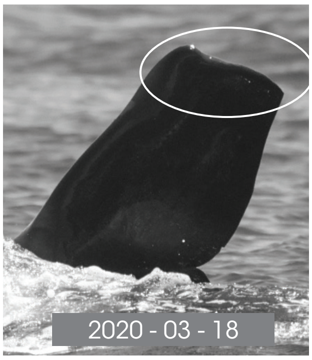

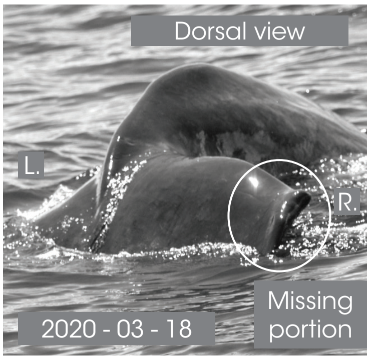

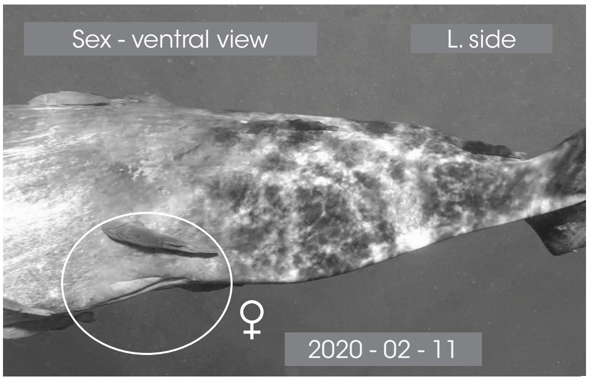

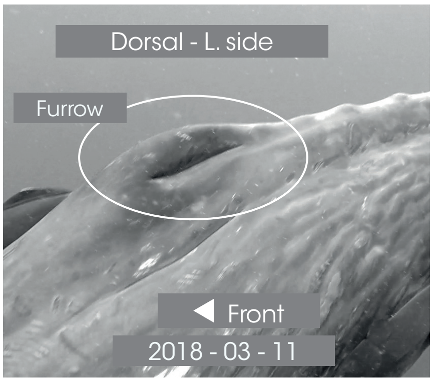

Dorsal - R. side

Front ▶

© Longitude 181 - Creation 2015 / Update 2020 - Concept : F. Sarano - Graphic design & illustrations : M. Sarano - © Pictures : S. Granzotto, F. Guérin, R. Heuzey, A. Preud'homme, V. & F. Sarano.

### DAREN

1ST OBS. : 2018 - 04 - 18 // LAST OBS. : 2020 - 03 - 12

| | | |
|---|---|---|
| JUVENILE | SEEN | B : 2018 - 17TH APRIL |
| DNA | - 2018<br>2019 - 2020 | ADDITIONAL INFORMATION |
| ♂ | | • Dark mandibular zone, without white marks, like his half-brother Roméo.<br>• Lucy'son & Jason's son.<br>• Roméo's half-brother |

Dorsal view // 2018 - 04 - 28

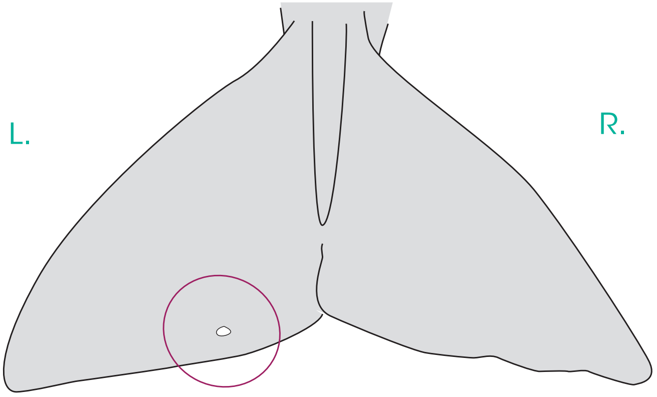

L. pectoral

Daren

◀ Front

L. pectoral

◀ Front

R. pectoral

UNHARMED

Front ▶

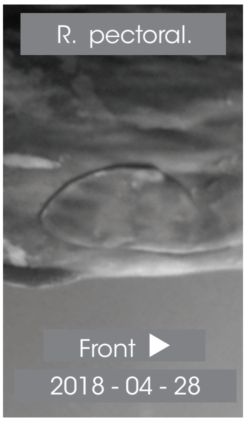

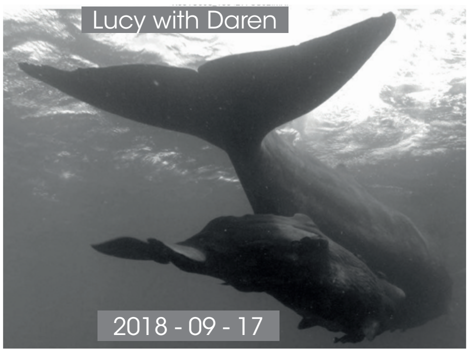

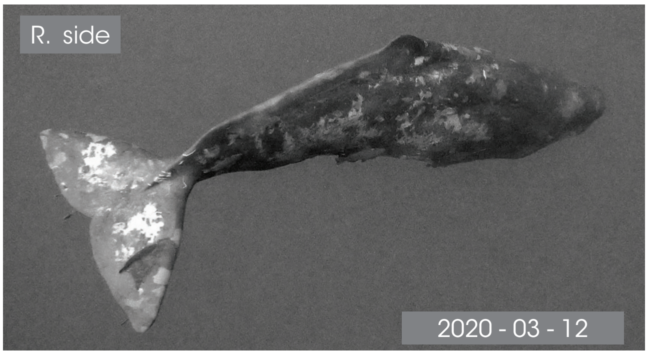

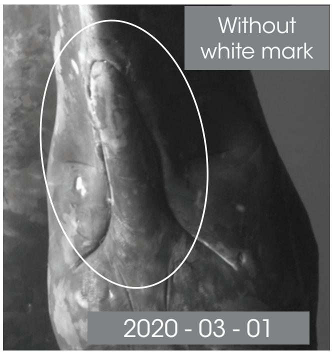

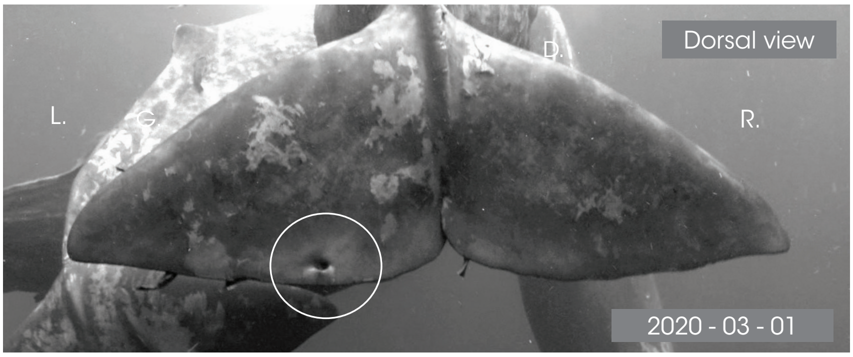

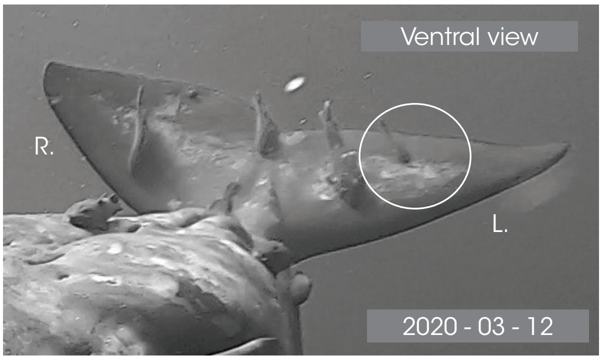

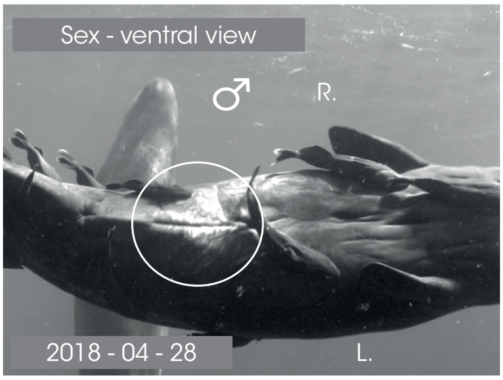

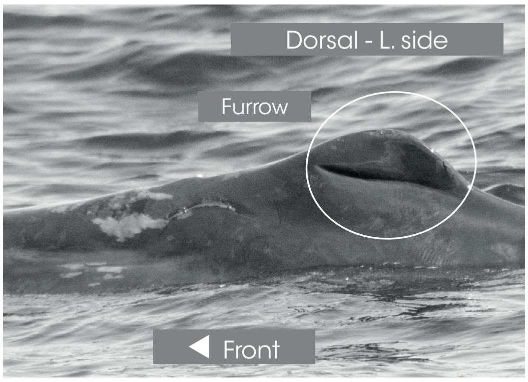

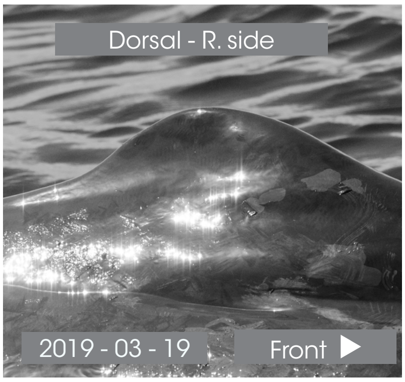

© Longitude 181 - Creation 2015 / Update 2020 - Concept : F. Sarano - Graphic design & illustrations : M. Sarano - © Pictures : S. Granzotto, F. Guérin, R. Heuzey, A. Preud'homme, V. & F. Sarano.

### ÉLIOT

**1ST OBS. : 2011 - 03 - 14 // LAST OBS. : 2020 - 03 - 19**

| JUVENILE | SEEN | B : 2011 - 14TH MARCH |
|---|---|---|
| DNA | 2011 - 2012<br>2013 - 2014<br>2015 - 2016<br>2017 - 2018<br>2019 - 2020 | ADDITIONAL INFORMATION |
| ♂ | | • Ventral white escutcheon.<br>• Dark mandibular zone, without white marks.<br>• New toothmarks on the caudal: end Sept. 2012.<br>• First teeth, May 2017.<br>• Adélie's son |

Dorsal view // 2017 - 03 - 16

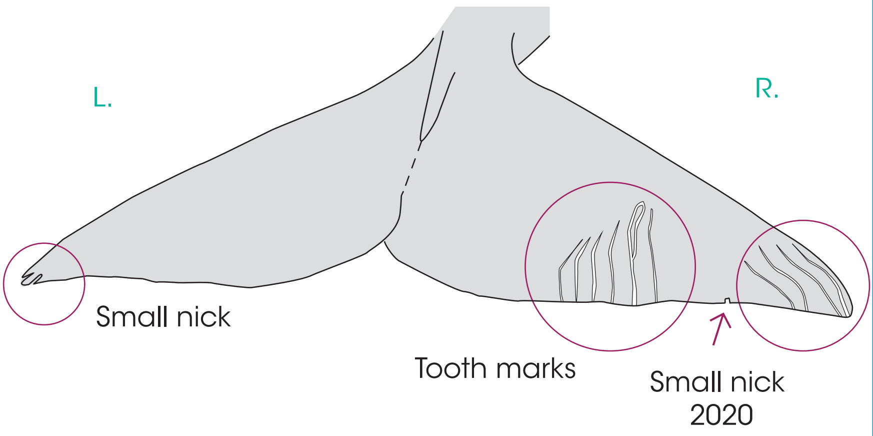

L. pectoral

UNHARMED

◀ Front

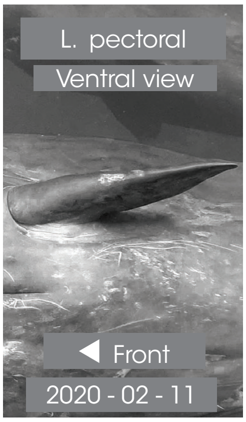

R. pectoral

UNHARMED

Front ▶

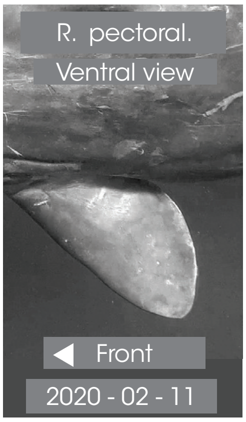

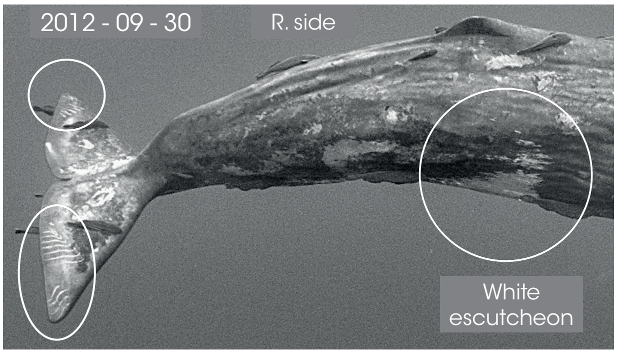

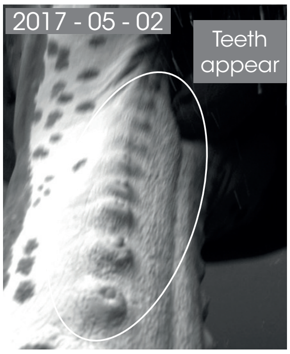

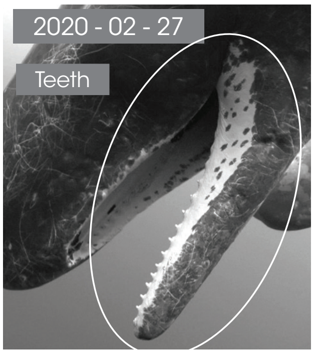

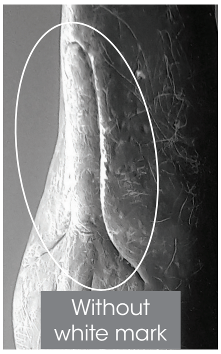

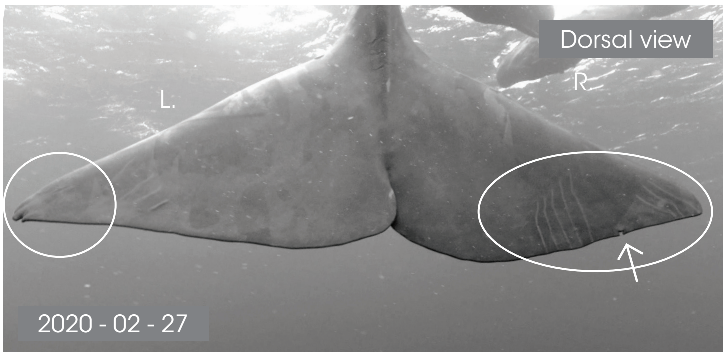

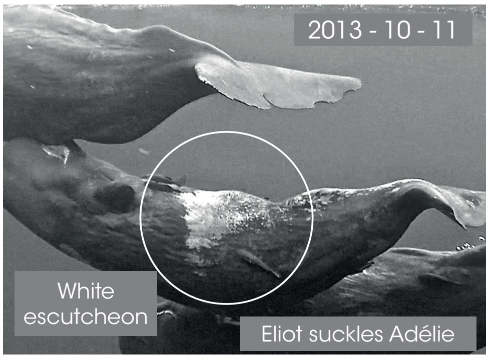

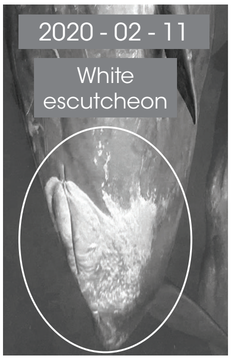

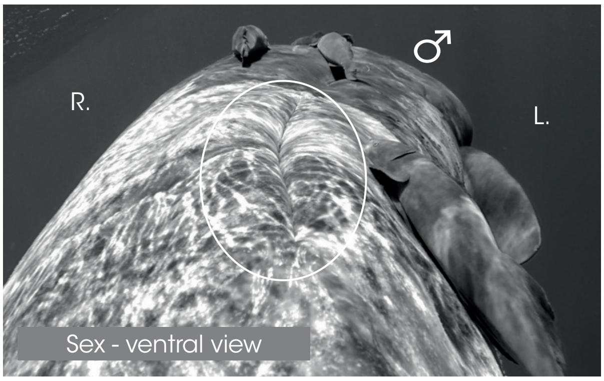

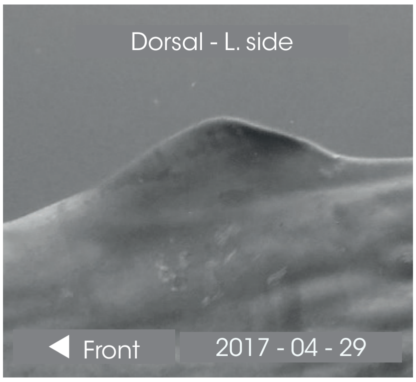

Dorsal - R. side

Front ▶

© Longitude 181 - Creation 2015 / Update 2020 - Concept : F. Sarano - Graphic design & illustrations : M. Sarano - © Pictures : S. Granzotto, F. Guérin, R. Heuzey, A. Preud'homme, V. & F. Sarano.

### LANA

1ST OBS. : 2019 - 02 - 21 // LAST OBS. : 2020 - 03 - 12

| JUVENILE | SEEN | B : 2019 - 20TH FEB |
|---|---|---|
| DNA | 2019 - 2020 | ADDITIONAL INFORMATION |
| ♀ | | • Bear ear shaped dorsal, protruding vertebras<br>• Irène's daughter & Noé's daughter.<br>• Arthur's half-brother.<br>• Ker2007's great granddaughter.<br>• Yukimi is its nurse (with evidence of milk). |

Dorsal view // 2020 - 03 - 01

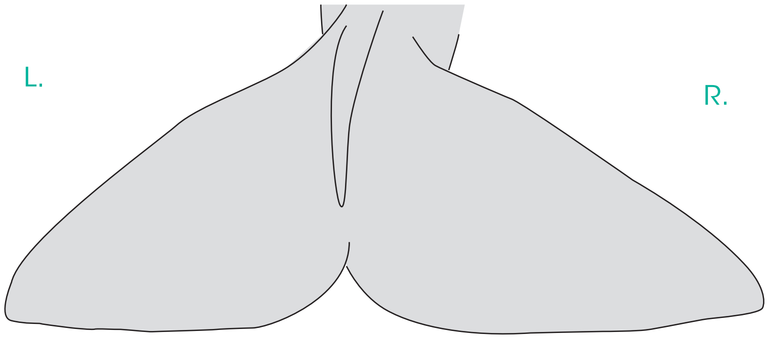

L. pectoral

UNHARMED

◀ Front

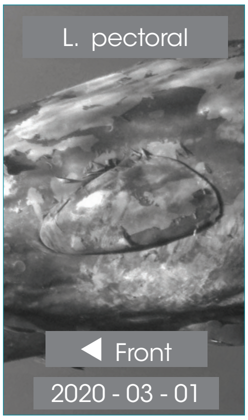

R. pectoral

UNHARMED

Front ▶

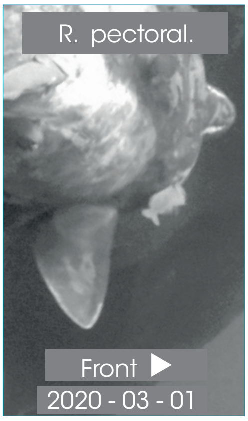

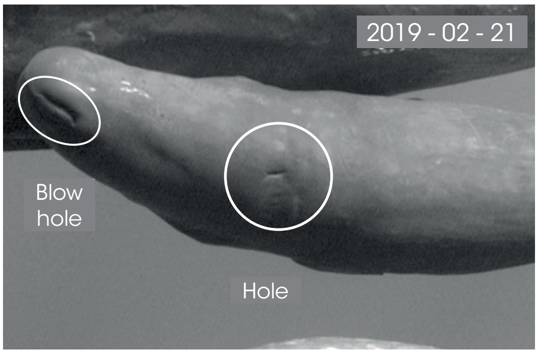

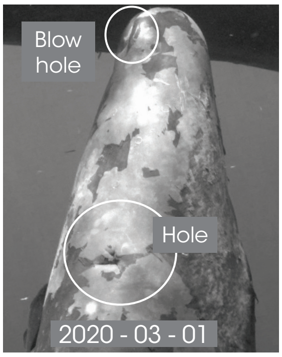

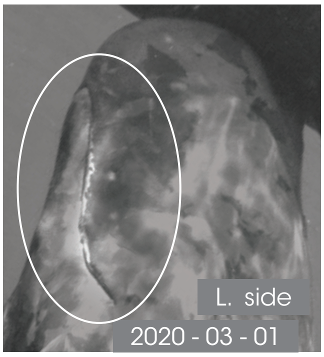

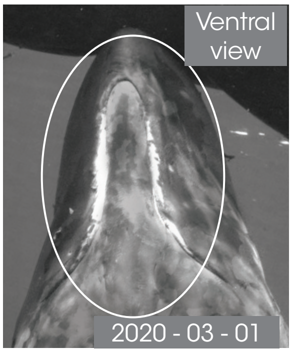

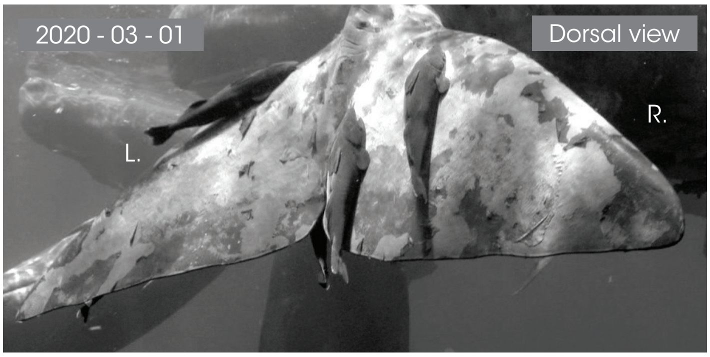

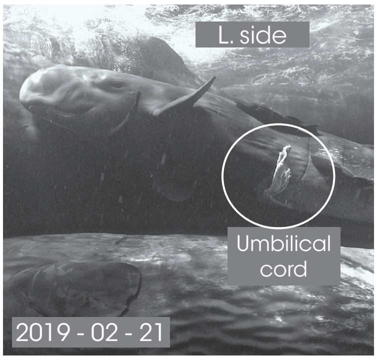

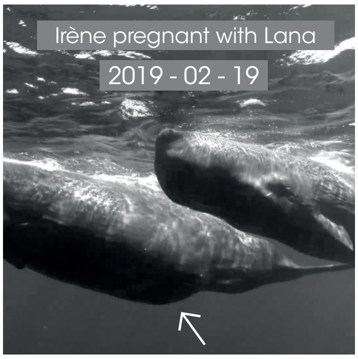

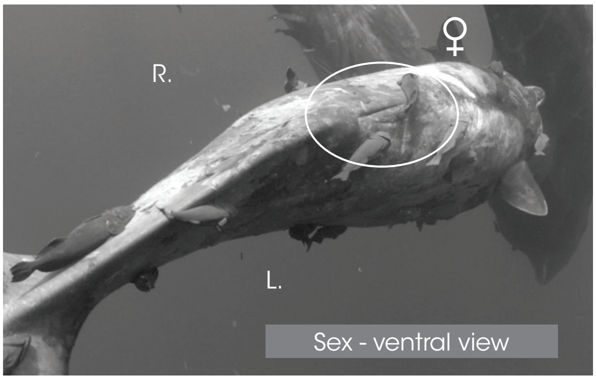

Dorsal - L. side

◀ Front

Dorsal - R. side

Front ▶

© Longitude 181 - Creation 2015 / Update 2020 - Concept : F. Sarano - Graphic design & illustrations : M. Sarano - © Pictures : S. Granzotto, F. Guérin, R. Heuzey, A. Preud'homme, V. & F. Sarano.

### MAURICE

1ST OBS. : 2011 - 03 // LAST OBS. : 2016-02-24

| JUVENILE | SEEN | B : 2011 - MAY ? |
|---|---|---|
| DNA | 2011 - 2012<br>2013 - 2014<br>2015 - ?<br><br>DEAD ? | ADDITIONAL INFORMATION |
| ♂ | | • Small triangular white spot close to the navel.<br>• Concave caudal.<br>• Missing since 24th Feb 2016. |

L. pectoral
MAURICE
◀ Front

L. pectoral
◀ Front

R. pectoral
MAURICE
Front ▶

R. pectoral.
Front ▶

Dorsal view // 2014 - 03 - 20

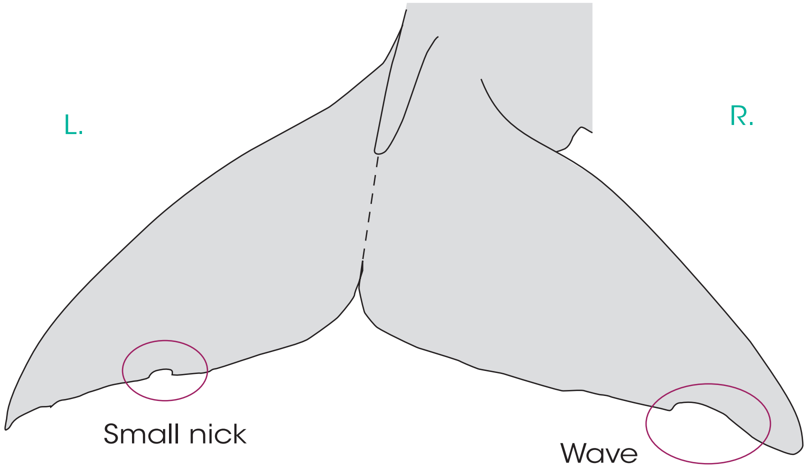

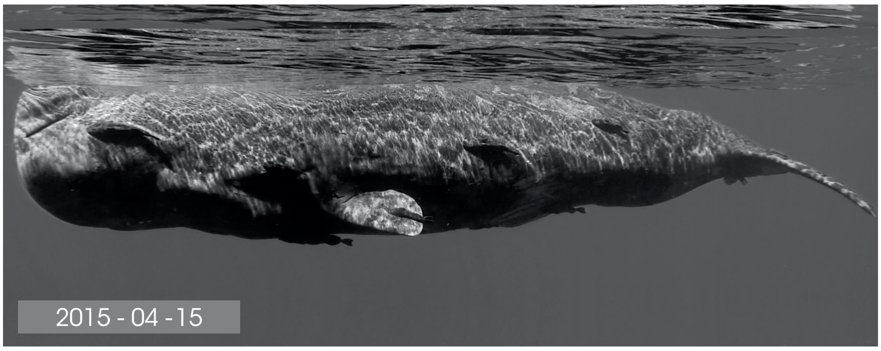

Genital slit

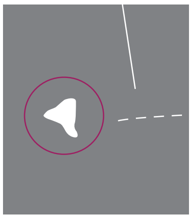

◀ Front Small white spot

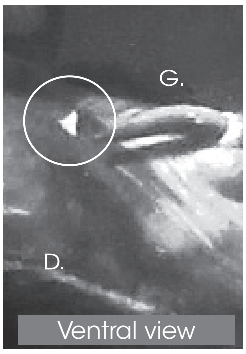

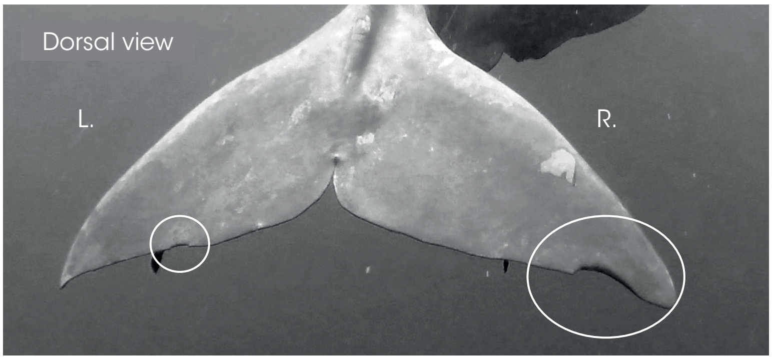

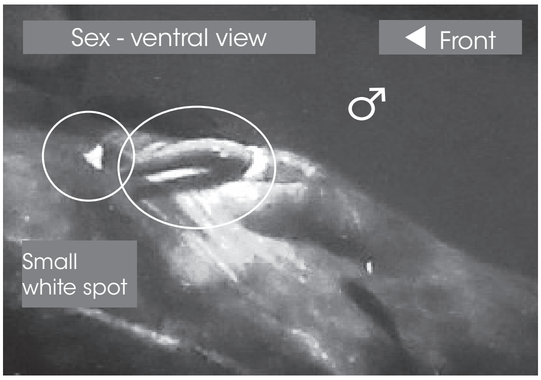

Dorsal - L. side
◀ Front

Dorsal - R. side
Front ▶

© Longitude 181 - Creation 2015 / Update 2020 - Concept : F. Sarano - Graphic design & illustrations : M. Sarano - © Pictures : S. Granzotto, F. Guérin, R. Heuzey, A. Preud'homme, V. & F. Sarano.

### MISS TAUTOU

1[ST] OBS. : 2016 - 02 - 24 // LAST OBS. : 2020 - 03 - 12

| JUVENILE | SEEN | B : ≈ 2016 - FEB |
|---|---|---|
| DNA | > 2016<br>2017 - 2018<br>2019 - 2020 | ADDITIONAL INFORMATION |
| ♀ | | • L. caudal tip : missing portion before 24/02/2016.<br>• White stripes close to the navel.<br>• Probably born beginning Feb, Issa's daughter.<br>• Germine's half-sister |

Dorsal view // 2016 - 03 - 18

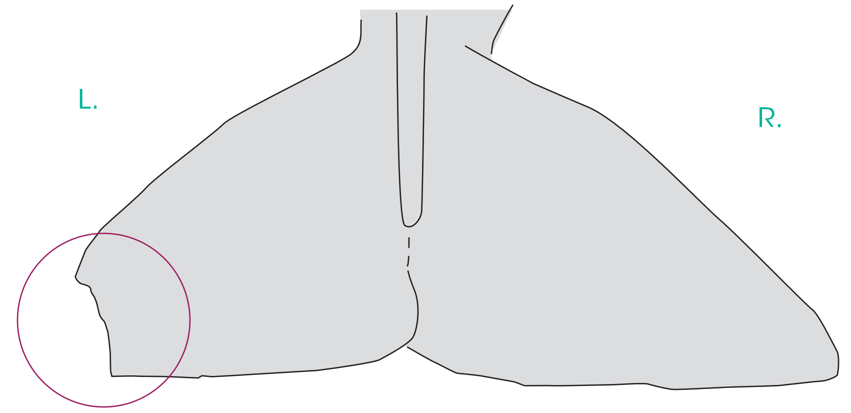

L. pectoral

Miss Tautou

◀ Front

L. pectoral

◀ Front

R. pectoral

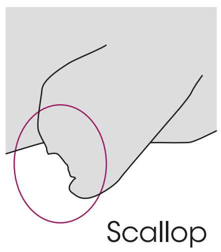

Front ▶

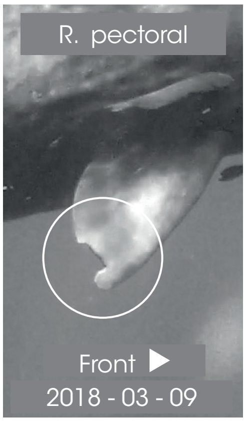

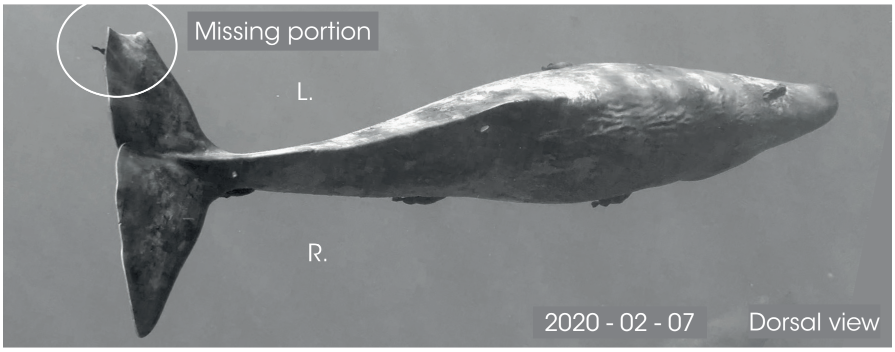

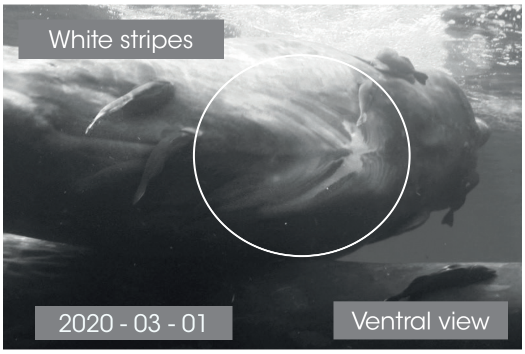

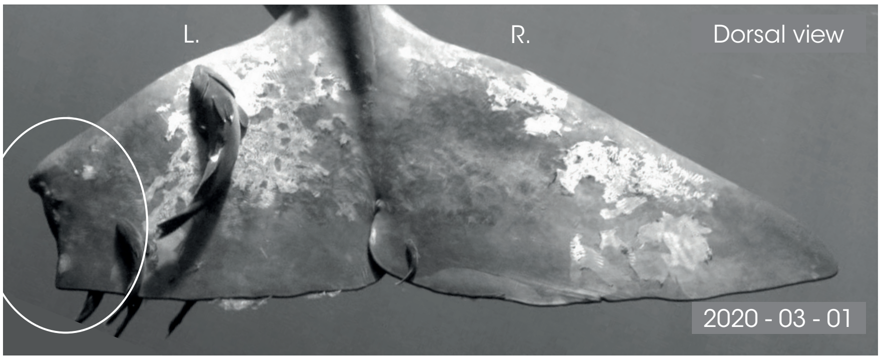

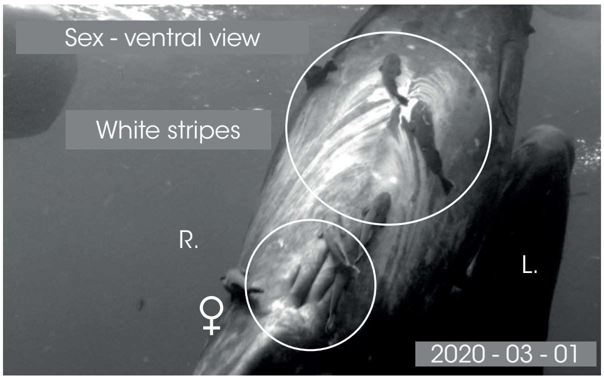

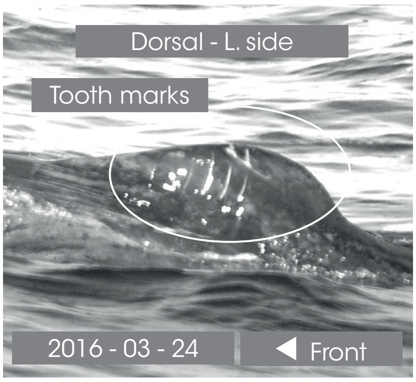

Dorsal - R. side

Front ▶

© Longitude 181 - Creation 2015 / Update 2020 - Concept : F. Sarano - Graphic design & illustrations : M. Sarano - © Pictures : S. Granzotto, F. Guérin, R. Heuzey, A. Preud'homme, V. & F. Sarano.

 p. 31

### ROMÉO

*1ST OBS. : 2013 - 03 - 13 // LAST OBS. : 2020 - 03 - 12*

| JUVENILE | *SEEN* | B : ≈ 2013 -3RD MARCH |
|---|---|---|
| *DNA* | 2013 - 2014<br>2015 - 2016<br>2017 - 2018<br>2019 - 2020 | *ADDITIONAL INFORMATION* |
| ♂ | | • Do not confuse with Maurice (scallop on the R. lobe).<br>• New small nick on the caudal L. lobe in 2020.<br>• Dark mandibular zone, without white marks, like his half-brother Daren.<br>• Lucy's son & Jonas's son.<br>• Daren's half-brother. |

Dorsal view // 2020 - 03 - 01

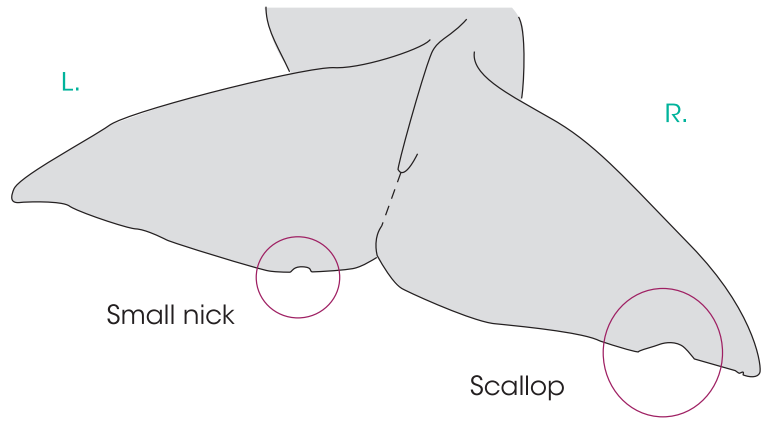

L. pectoral

roméo

◀ Front

L. pectoral

◀ Front

R. pectoral

roméo

Front ▶

R. pectoral

Front ▶

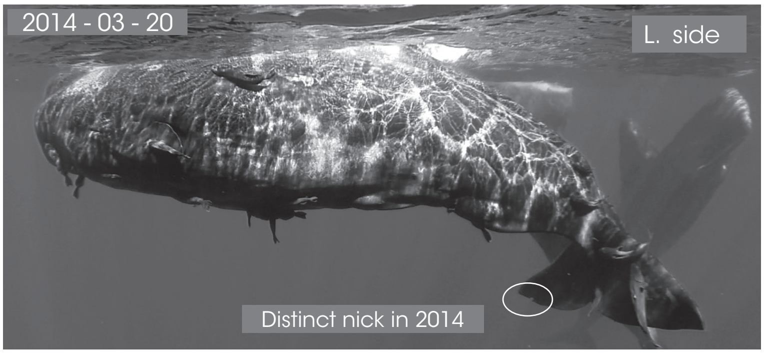

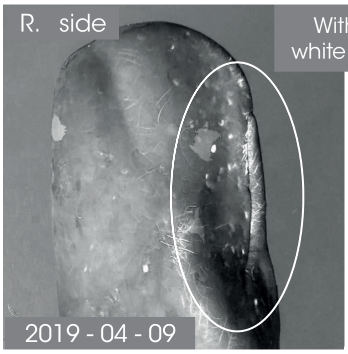

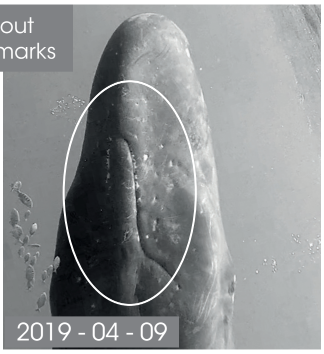

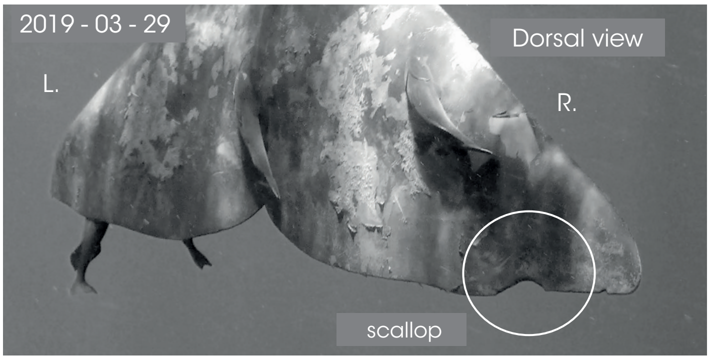

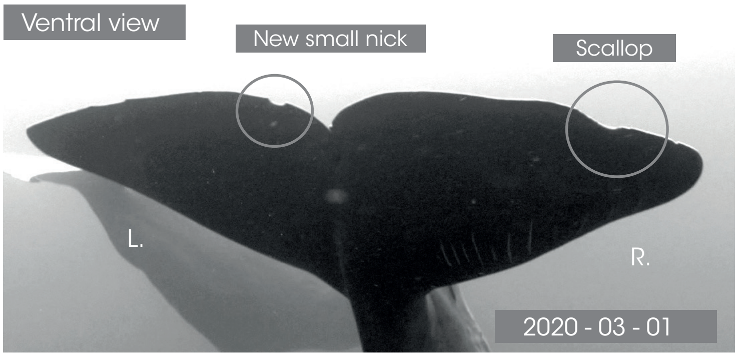

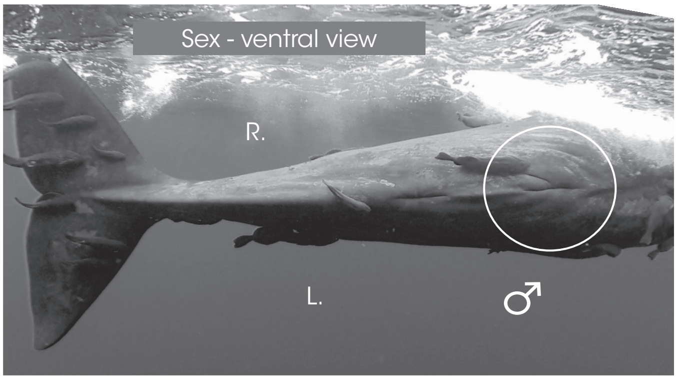

Dorsal - L. side

◀ Front

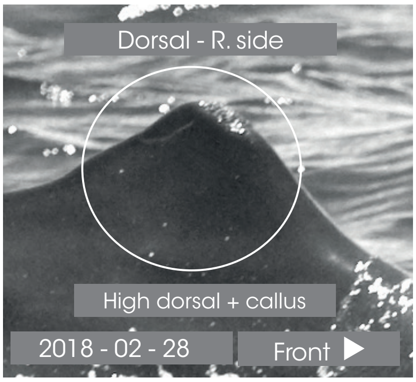

© Longitude 181 - Creation 2015 / Update 2020 - Concept : F. Sarano - Graphic design & illustrations : M. Sarano - © Pictures : S. Granzotto, F. Guérin, R. Heuzey, A. Preud'homme, V. & F. Sarano.

LONGITUDE 181
La Voix de l'Océan

IDENTITY CARDS
#### SPERM WHALES OF MAURITIUS I.

un océan de vie — www.unoceandevie.com — TERIA — Marine Megafauna Conservation Organisation

### TACHE BLANCHE

*1ST OBS. : 2011 - 06 // LAST OBS. : 2020 - 03 - 19*

| JUVENILE | *SEEN* | B : ≈ 2011 - JUNE |
|---|---|---|
| *DNA* | 2011 -2012<br>2013 - 2014<br>2015 - 2016<br>2017 - 2018<br>2019 - 2020 | *ADDITIONAL INFORMATION* |
| ♂ | | • White spot between navel and genital slit.<br>• Concave caudal.<br>• Delphine's son.<br>• Chesna's half-brother. |

L. pectoral

**UNHARMED**

◀ Front

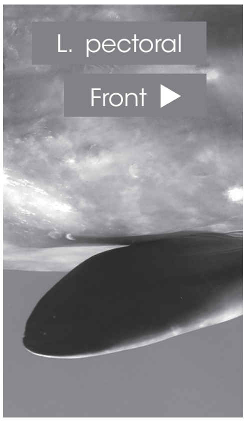

Dorsal view // 2018 - 04 - 18

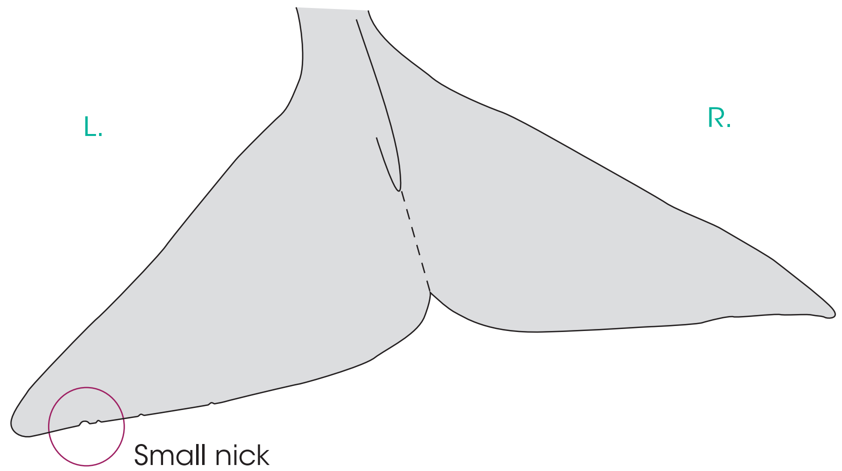

R. pectoral

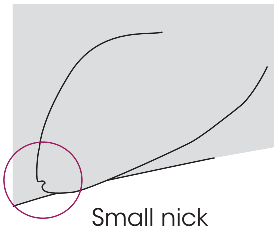

Front ▶

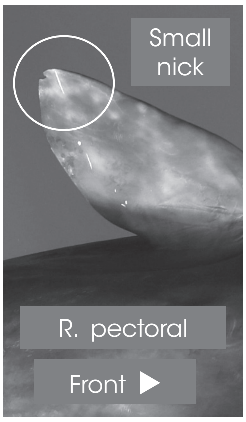

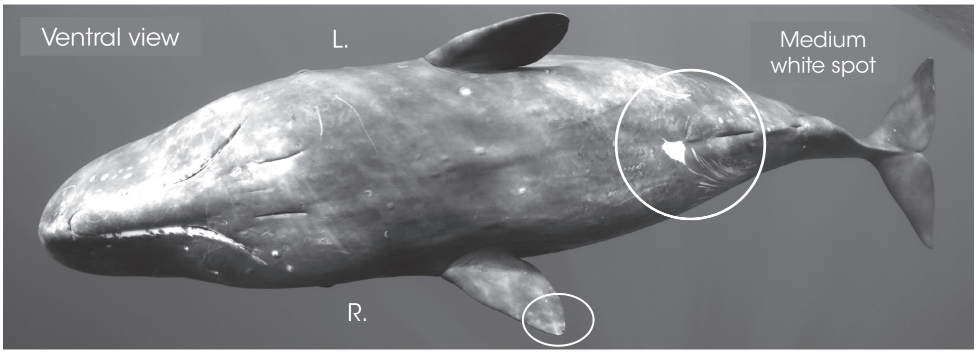

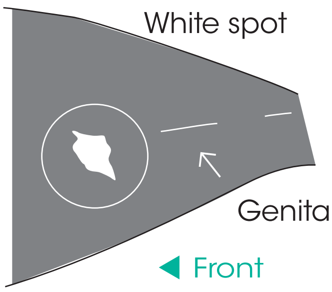

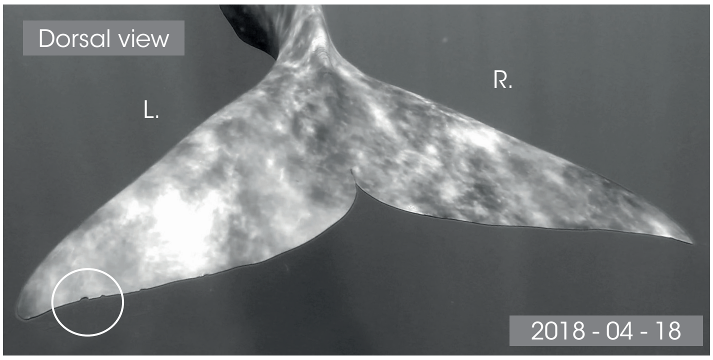

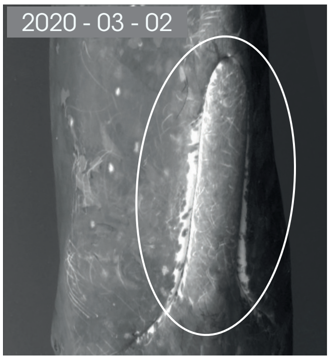

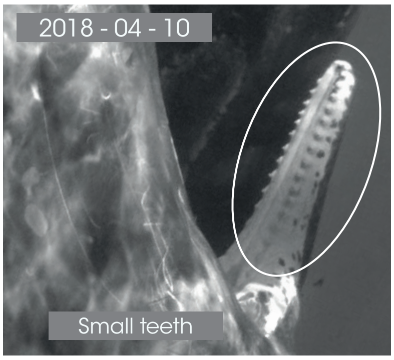

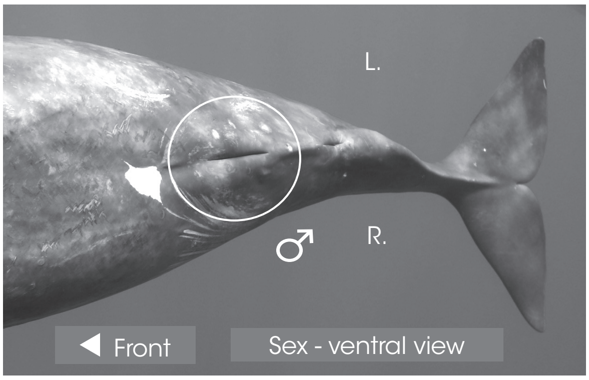

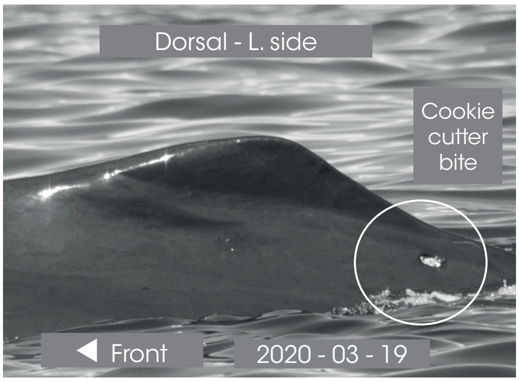

Dorsal - R. side

Front ▶

© Longitude 181 - Creation 2015 / Update 2020 - Concept : F. Sarano - Graphic design & illustrations : M. Sarano - © Pictures : S. Granzotto, F. Guérin, R. Heuzey, A. Preud'homme, V. & F. Sarano.

### ZOÉ

**1ST OBS. : 2013 - 12 - 16 // LAST OBS. : 2020 - 03 - 18**

| JUVENILE | *SEEN* | B : ≈ 2013 - DEC. ? |
| --- | --- | --- |
| *DNA* | 2013 - 2014<br>2015 - 2016<br>2017 - 2018<br>2019 - 2020 | *ADDITIONAL INFORMATION* |
| ♀ | | • Tooth marks on the L. lobe of the caudal.<br>• New hole on the R. lobe of the caudal in 2019.<br>• Concave caudal.<br>• Caroline's daughter.<br>• Alexander's half-sister. |

Dorsal view // 2019 - 05 - 05

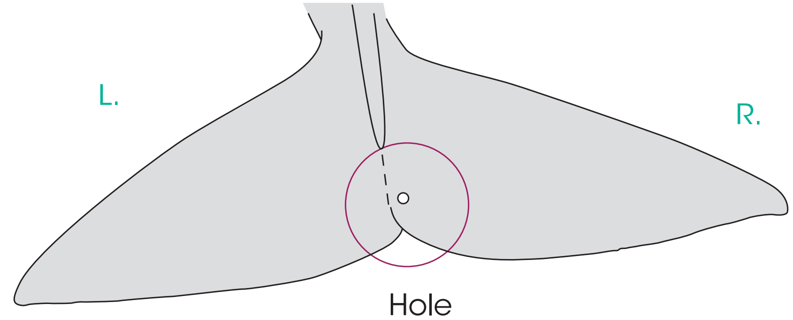

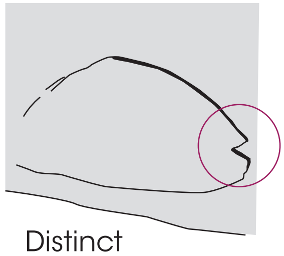

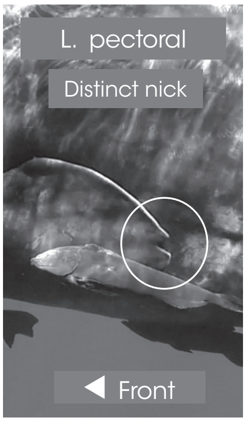

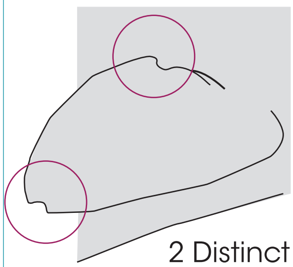

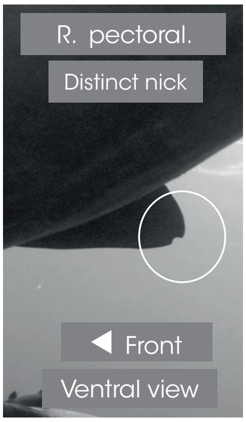

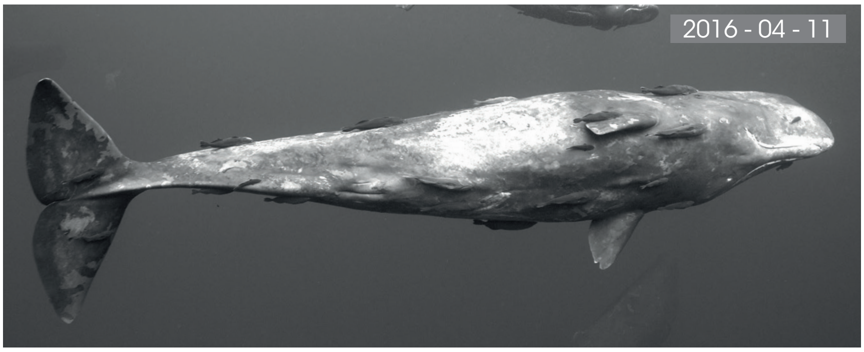

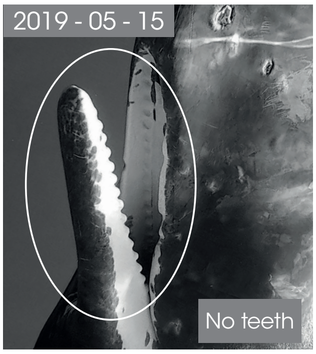

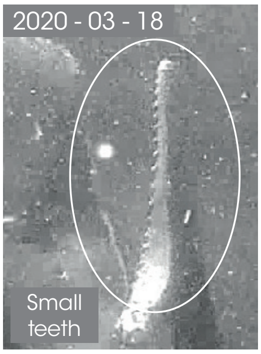

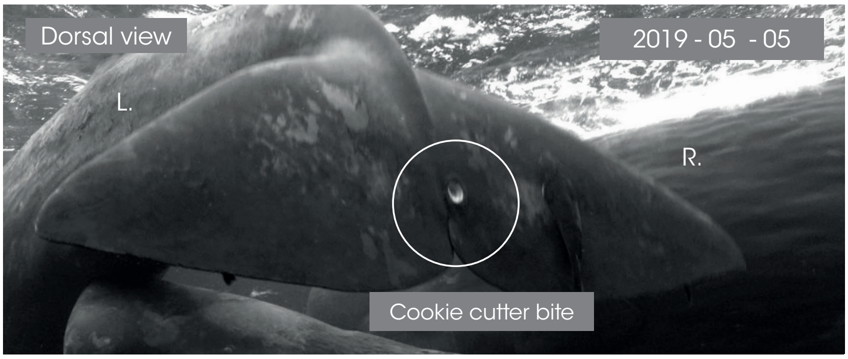

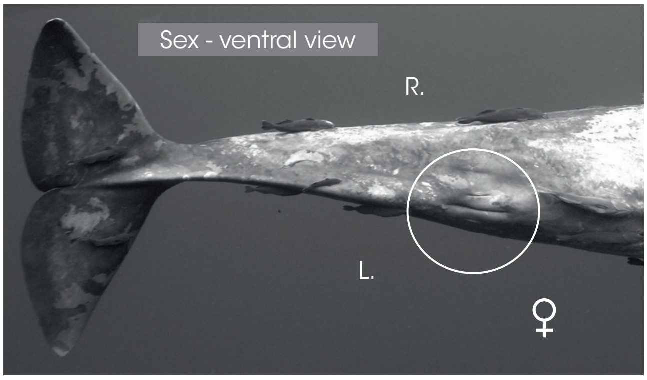

Dorsal - L. side

Front

Dorsal - R. side

Front

© Longitude 181 - Creation 2015 / Update 2020 - Concept : F. Sarano - Graphic design & illustrations : M. Sarano - © Pictures : S. Granzotto, F. Guérin, R. Heuzey, A. Preud'homme, V. & F. Sarano.
